# Spinal circuit regionalization diversifies motor output along the vertebrate body axis

**DOI:** 10.64898/2026.07.02.736051

**Authors:** Stavros Papadopoulos, August Winther, Florina Alexandra Toma, Kareem Elassy, Clara Julseth, Remi Ronzano, Ying Zhang, Tim P. Vogels, Didier Le Ray, Lora B. Sweeney

## Abstract

The evolution of the vertebrate limb-torso-limb body plan drove a diversification of motor behavior. How neural circuits are organized to generate distinct outputs across body regions, however, remains unknown. Here, we construct a cross-species, spatiotemporal atlas of spinal interneurons spanning the rostrocaudal and developmental axes of frogs and mice. We uncover a common framework of interneuron regionalization that mirrors each vertebrate’s body plan and arises through Hox-dependent regulation of rostrocaudal neurogenesis. Combining electrophysiology with computational modeling, we show that changing regional circuit composition is sufficient to respecify motor output. Together, these findings establish a causal link between developmental patterning, circuit architecture, and motor function, identifying interneuron regionalization as a fundamental organizational principle linking body-plan evolution to motor diversity.

## INTRODUCTION

Vertebrate evolution was transformed by two highly correlated events: the development of limbs and the transition from aquatic to terrestrial habitats.^1–3^ These events resulted in an explosion of new and highly coordinated motor patterns, as the body plan shifted from a largely uniform axial to a regionalized limb-torso-limb organization.^4^ This presented a neural control challenge: motor networks must change from regulating broad rostral-to-caudal undulatory waves to instead controlling highly regionalized, cross-body coordinated motor responses characteristic of limbed vertebrates.^5,6^ Despite the revolutionary impact of these motor innovations on how vertebrates navigate their environment, the underlying adaptive changes in spinal circuits that gave rise to these new localized patterns along the rostrocaudal axis remain unclear.

Spinal circuits are composed of distinct inhibitory and excitatory cardinal interneuron types that coordinate motor neuron output.^5,7,8^ For motor neurons, we have a clear understanding of how molecular identity and projection pattern vary along the rostrocaudal axis, and what specifies these differences.^9–13^ However, for the networks of interneurons that regulate their output, we do not.^5,8,9,14^ Motor neurons are regionalized into brachial, thoracic and lumbar subtypes via Hox and downstream effector gene expression.^15–20^ Losing expression of a single Hox gene, *hoxc9*, results in thoracic motor neurons assuming limb level-like characteristics.^18,21,22^

For interneurons, unlike motor neurons, our understanding of regional differences along the rostrocaudal axis is more limited. Only a few populations have been examined in detail, and for these, only select examples exist of differences in number, molecular identity and connectivity.^14,23–25^ The ventral V1 inhibitory neuron class, a primary regulator of motor neuron coordination,^26–28^ is increased two-fold at limb compared to thoracic levels, and additionally exhibits level-specific subtype variation.^29^ These differences, similar to motor neurons, have been linked to the thoracic expression of *hoxc9*, as have differences in spinocerebellar neuronal cell types along the rostrocaudal axis.^29–31^ Excitatory V2a neurons vary in their molecular identity, connectivity and relative distributions at fore- and hindlimb levels.^32^ Finer-scale, level-specific variation in subtype composition and settling position has also been suggested for ventral populations during development.^31,33^ Recently, in the adult spinal cord, single-cell and spatial transcriptomic analysis has also begun to identify coarse, level-specific enrichment of subtypes along the rostrocaudal axis, leaving open the question of when these differences emerge, how they are specified and what are their functional consequences.^34,35^ Collectively, this suggests an overarching, but as of yet largely untested, principle of spinal regionalization in which developmental and evolutionary specialization of motor output at each body region generates the immense diversity of tetrapod locomotion.

How spinal interneuron variation between limb-torso-limb levels relates to movement pattern differentiation along the body axis however is still unknown. At forelimb levels, integration of multimodal descending and sensory information produces fine-tuned grasping responses.^8^ In contrast, at thoracic levels, torso musculature supports postural control, with tightly controlled stabilization mechanisms mediated in large part by vestibular-driven spinal circuits.^36,37^ Such radical differences in motor control by level beg the question of whether a broad limb-torso-limb logic of neuron architecture governs the regional specialization of function. Are motor- and sensory-related interneurons differentially distributed? Do regional differences in neuron number and local circuit architecture influence output? Do Hox genes represent a developmental master regulator linking interneuron number, circuit connectivity and motor coordination?

Here, we determine that all interneuron networks in the spinal cord are regionalized, identify mechanisms driving regionalization, and define how these regional differences shape motor output. Exploiting the diversity of vertebrate movement, we provide a combined cell-type and level-specific atlas of spinal neuron architecture along the rostrocaudal axis of the frog and mouse, two species which last shared a common ancestor ∼360 million years ago.^38^ This cross-species approach reveals that tuning excitatory and inhibitory interneuron number and distribution by rostrocaudal level is a common means to regionalize spinal circuits and differentiate movement patterns during tadpole-to-frog metamorphosis and vertebrate evolution. Loss-of-function analysis of Hoxc9 demonstrates Hox genes serve as global regulators of rostrocaudal differences in interneuron number via regulation of neural progenitors and their proliferation. Employing a combination of computational modeling and electrophysiology, we identify network effects of changing one spinal region into another, directly linking regional architecture to function. Our findings show that regional variation in interneuron distributions is a developmentally reliable and evolutionarily pliable means to differentiate motor output along the vertebrate body axis.

## RESULTS

### Interneurons transition from a uniform axial to a limb-torso-limb regional architecture during frog metamorphosis

Amphibians, such as the *Xenopus* frog, transition from tail to limb locomotion during metamorphosis, recapitulating the terrestrialization of vertebrate behavior in their development. This results in a drastic remodeling of the body plan along the longitudinal axis from repeated axial segments to a limb-torso-limb arrangement.^39^ Such remodeling provides the ideal starting point to determine how the body plan is reflected in the neural architecture of the spinal cord.

Capitalizing on the well-defined molecular classification of cardinal classes in the mammalian spinal cord,^7,40–42^ and its recently established conservation in both transcriptional and neurotransmitter profile between frog and mouse,^43^ we evaluated the rostrocaudal distribution of eight interneuron classes from tadpole to frog: motor-related excitatory V0v, V3v, V2a and dI3; inhibitory V1 and dI6; and sensory-related excitatory dI1 and dI5 interneurons. We targeted our analysis to three key stages (**Figure 1A**): Nieuwkoop and Faber (NF) stage 47, the most mature swim stage before limb circuit expansion; stage NF55, when limb-innervating motor neuron and spinal inhibitory V1 interneuron numbers peak and the first spontaneous limb movements are observed; and stage NF66, the first post-metamorphic frog stage with fully ambulatory behavior.^39,44–47^ At each stage, the number and distribution of each interneuron population was scored using established molecular markers at brachial, thoracic and lumbar levels (**Figure 1B-C, S1A-B, S2A-B, S5A**).^42,48–56^ Limb-innervating lateral motor neuron columns (LMCs) were used as landmarks to demarcate levels (**Figure S4A**),^39,57,58^ enabling side-by-side analysis of motor and interneurons.

**Figure 1.**
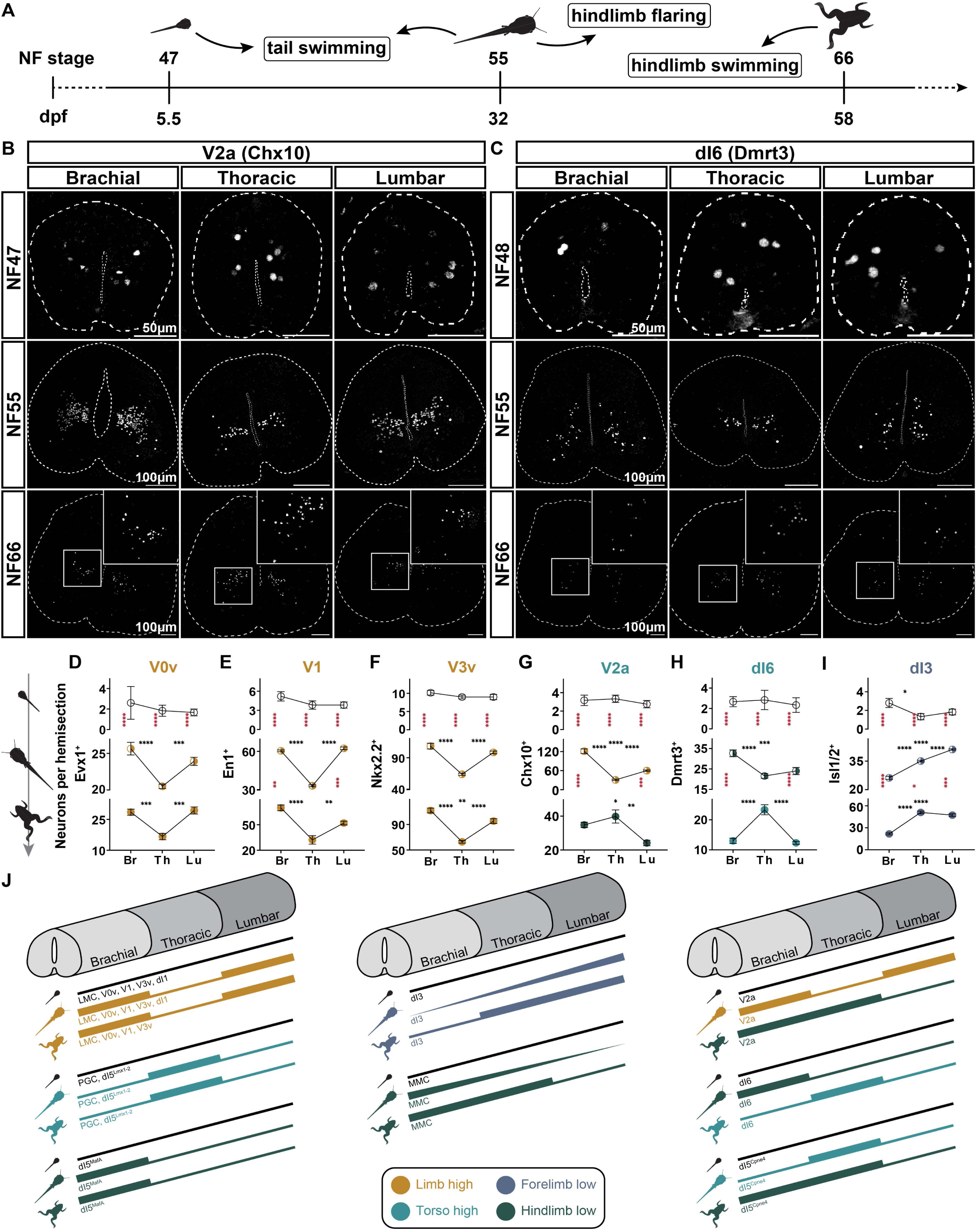
The swim-to-limb transition of neuronal architecture. **A.** Examined time points representing the swim-circuit (NF47), the metamorphic limb-circuit (NF55), and juvenile (NF66) limb circuit, with their respective main locomotor behaviors. **B-C.** V2a (Ch×10^+^) and dI6 (Dmrt3^+^) neurons along the brachial-thoracic-lumbar axis, over three locomotor modes. Scale bars; 50μm for NF47, 100μm for NF55 and NF66. **D-I.** Neuron distribution along the brachial-thoracic-lumbar axis is uniform in the swim-circuit, but gets regionalized with the emergence of the limb-circuit, acquiring a neuron class-distinct pattern. V0v, Evx1^+^; V1, En1^+^; V3v, Nkx2.2^+^; V2a, Ch×10^+^; dI6, Dmrt3^+^; dI3, (dorsal)Isl1/2^+^. All data reported as mean ± SEM. One-way ANOVA and Tukey’s multiple comparisons test; * p<0.1, ** p<0.01, *** p<0.001, **** p<0.0001. Black asterisks signify significant differences across rostrocaudal levels of a single developmental stage; red asterisks signify significant differences across developmental stages at the same rostrocaudal level. For each animal, 3-6 transverse hemisections per level were scored, with NF55; N=12-14, NF47 and NF66; N=2-5 animals. **J.** Schematic summary of rostrocaudal distribution patterns, unique or shared between developmental stages/locomotor modes, for all neuron classes and subpopulations examined in this study.

This analysis revealed striking level-specific changes in interneuron number and distribution during the tail-to-limb transition. At NF47 swim stages, all examined populations including motor and interneurons were present in low numbers (**Figure 1D-I, S4A-F**). These neurons exhibited largely uniform rostrocaudal distributions and acquired broad ventral or dorsal settling positions across the transverse plane of the spinal cord (**Figure 1B-C, S1A-D**). These results extend previously characterized knowledge of larval tadpole and zebrafish V0v, V1 and V2a interneuron architecture.^59–65^

With the emergence of limbs, all initially uniform spinal interneuron distributions diverged across the body axis (**Figure 1D-J**). V0v, V1 and V3v interneurons shared a common and developmentally stable high-limb/low-torso pattern (*Limb high*; **Figure 1D-F**). This pattern matched that of limb motor neurons (**Figure S4A**), implicating these regional changes in local regulation of limb-level motor output. Other interneuron populations had differing and developmentally dynamic rostrocaudal patterns. V2a interneurons exhibited a dynamic distribution that was first evenly distributed at tadpole stages, then high at limb levels during the tail-plus-limb stages (*Limb high*), and finally, high at both brachial and thoracic levels at frog stages (*Hindlimb low*; **Figure 1B, 1G**). dI6 interneurons transitioned from a *Hindlimb low* to a *Torso high* distribution as tadpoles matured to frogs. Excitatory dI3 neurons showed a *Forelimb low* distribution, with consistently high numbers at hindlimb levels and increasing numbers of neurons at thoracic levels over developmental time (**Figure 1C, 1H**).

Dorsal excitatory neurons also varied in their rostrocaudal patterns. dI1 interneurons had a *Limb high* pattern, paralleling the distribution of LMC motor neurons and suggesting a heightened regulatory role at limb regions (**Figure S4C**). For dI5 interneurons, we examined three partially overlapping subpopulations each expressing the canonical dI5 markers Lmx1-2, MafA and Cpne4, respectively.^42,55,56^ These subpopulations were detected in increasingly higher numbers as development progressed. Each exhibited a different rostrocaudal pattern: enriched either at only thoracic levels (*Torso High*), or at brachial-and-thoracic or brachial levels (*Hindlimb Low*; **Figure S4D-F, 1J**). Notably, along the rostrocaudal axis and between stages, the changes in interneuron numbers we observed did not match global changes in the per-section neuron number, cell density or volume of the spinal cord (**Figure S4H-J**).

These findings show that neuron distributions along the spinal cord vary over metamorphosis in a cell-type specific manner, suggesting a region-by-region, circuit-to-function correspondence as behavior changes from tadpole to frog. To begin to query the functional implications of these changing neuron distributions, we assumed differences in the inhibition/excitation balance of spinal regions across rostrocaudal levels and developmental phases would diversify motor output. Across all stages and levels, excitatory neurons comprised three-quarters of the motor-related circuit and inhibitory neurons, one-quarter (**Figure S4K-M**). Rostrocaudal excitation and inhibition, subdivided into ipsilateral or commissural populations based on projection patterns in mice,^7^ changed over time: first, uniform in the NF47 tadpole, positioned to drive the rostral-to-caudal undulatory waves of tail swimming (**Figure S4K**) and surprisingly also uniform in the NF55 tadpole, despite the strong regionalization of individual neuron distributions (**Figure 1J**). Only when the limb circuit was fully functional in NF66 froglets did cross-level differences emerge between each ipsilateral/commissural population (**Figure S4L-M**). This result underscores the strong effect cumulative changes in neuron distribution across the rostrocaudal axis can exert on circuit-level properties (**Figure 1D-I, S4K-L**). Circuit regionalization thus parallels output regionalization, implicating the fine-tuning of local cell-type numbers in changing locomotor responses.

### Conservation and divergence of interneuron architecture between four-limbed vertebrates

The emergence of rostrocaudal enrichments in interneuron numbers with the addition of the limb circuit in the frog raises the question of whether, and if so what, functional differences these regional differences confer. To examine the significance of rostrocaudal specialization, we extended our analysis to a second four-limb vertebrate with highly variant motor behavior to that of the frog: the mouse. Whereas frog locomotion is dominated by bilaterally synchronous kicking, mouse locomotion is notable for its various speed-dependent, alternating gaits and increased repertoire of dexterous behaviors.^5,8,66,67^ Such a comparison between distant tetrapods simultaneously tests the evolutionary questions of whether shared rostrocaudal principles define a conserved ambulatory circuit, and whether rostrocaudal patterns evolved to facilitate differences in locomotor patterns (**Figure 2A-B**).

**Figure 2.**
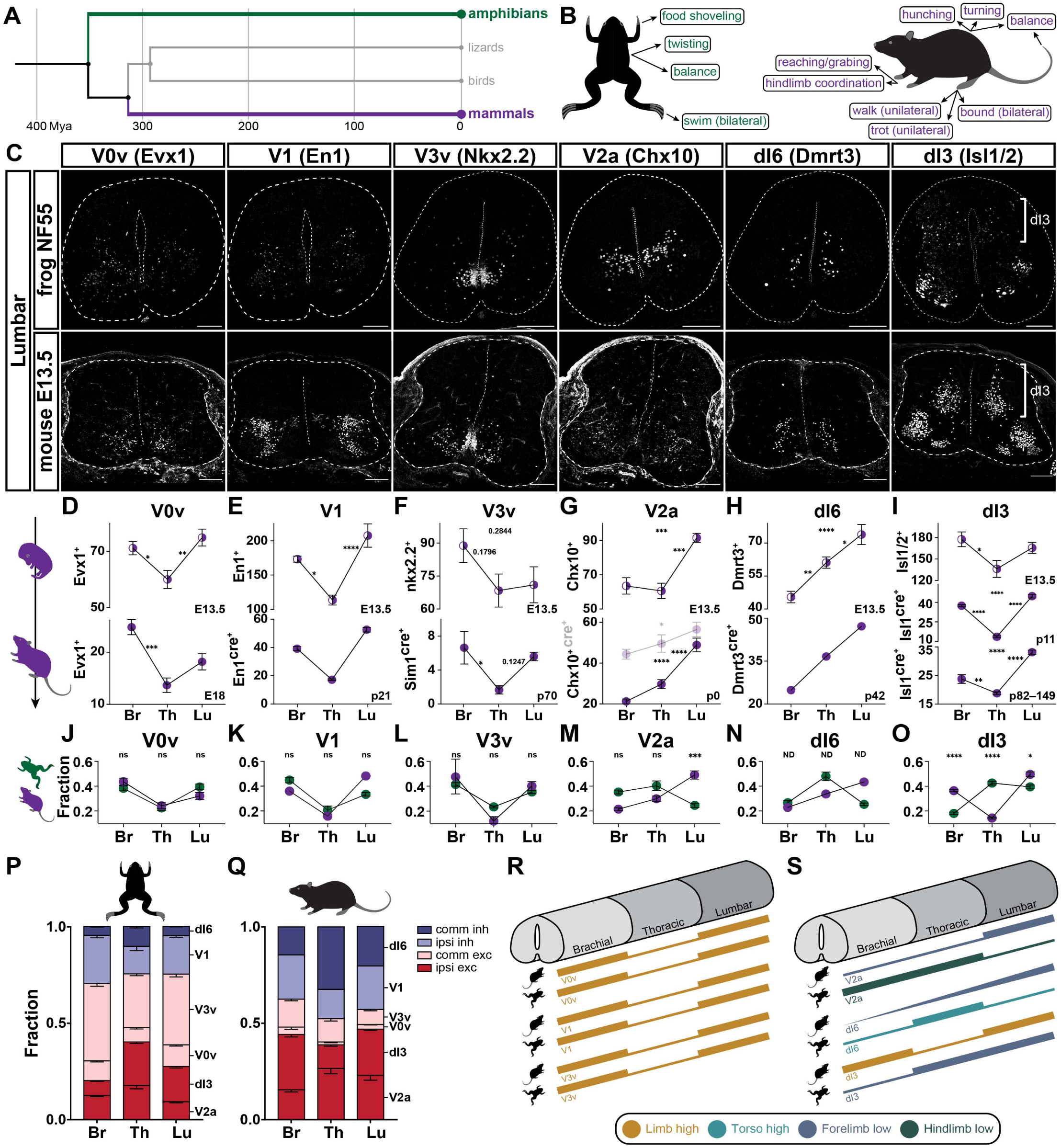
Conservation and divergence of rostrocaudal neuron architecture between mouse and frog. **A.** Tetrapod phylogenetic tree depicting the last common ancestor of amphibians and mammals lived approximately 360 million years ago.^38^ **B.** Movements and locomotor functions at different rostrocaudal levels in mice and frogs. **C.** Neurons with frog-mouse conserved rostrocaudal patterns V0v (Evx1^+^), V1 (En1^+^), V3v (Nkx2.2^+^), and neurons with divergent rostrocaudal patterns V2a (Ch×10^+^), dI6 (Dmrt3^+^), dI3 (Isl1/2^+^_dorsal_), shown at the lumbar levels. Scale bars, 100μm. **D-I.** Neuron distribution patterns along the brachial-thoracic-lumbar axis get cemented as development progresses but vary between classes. Top row, E13.5; bottom row, V0v, E18; V1, p21 (En1^cre/wt^; adapted from *Sweeney et al. 2018*);^29^ V3v, p70 (Sim1^cre/wt^, only ventral population); dI3, p82-149 (Isl1^cre/wt^;Ai14^fl/wt^, excluding motor neurons); dI6, p42 (Dmrt^cre/wt^; adapted from *Perry et al. 2019*),^68^ V2a, p0 (Ch×10^cre/wt^; adapted from *Hayashi et al. 2018*)^32^; middle row, dI3, p11 (Isl1^cre/wt^;Ai14^fl/wt^, excluding motor neurons). All data reported as mean ± SEM. One-way ANOVA and Tukey’s multiple comparisons test; * p<0.1, ** p<0.01, *** p<0.001, **** p<0.0001. For each animal a minimum of three (n=3) transverse hemisections per level were scored for two to nine (N=2-9) animals. **J-O.** Conserved (**J-L**) and divergent (**M-O**) neuron distribution patterns along the brachial-thoracic-lumbar axis between juvenile frogs and mature mice, depicted as rostrocaudal fractions – the cell count of a neuron class at each rostrocaudal level is divided by the sum at all three rostrocaudal levels for this neuron class, e.g. for a brachial fraction, X_Br_/(X_Br_+X_Th_+X_Lu_). Data reported as mean ± SEM. One-way ANOVA and Tukey’s multiple comparisons test; ns, p>0.1; *, p<0.1; **, p<0.01; ***, p<0.001; ****, p<0.0001; ND, not determined. For each animal a minimum of three (n=3) transverse hemisections per level were scored for two to six (N=2-6) animals. **P, Q.** Fractions of motor-related interneurons along the brachial-thoracic-lumbar axis for frog (**P**) and mouse (**Q**), showing rostrocaudal regionalization in both circuits, but with the inhibitory fraction being ∼0.25 and ∼0.5 for frog and mouse, respectively. Fractions are calculated per rostrocaudal level – the cell count of each neuron class at a rostrocaudal level is divided by the sum of cell counts for all six neuron classes at that rostrocaudal level, e.g. for the brachial fraction of an X_1_ neuron class, Br_X1_/(Br_X1_+Br_X2_+Br_X3_+Br_X4_+Br_X5_+Br_X6_). Data are sorted and colour-coded by projection pattern and function: blue, commissural inhibitory; light blue, ipsilateral inhibitory; pink, commissural excitatory; red, ipsilateral excitatory. All data reported as mean with SD. **R, S.** Schematic summary of neuron rostrocaudal distribution patterns, conserved (**R**) or divergent (**S**) between post-metamorphic frog and mature mouse.

In mice, we thus created a rostrocaudal atlas of excitatory and inhibitory spinal interneurons at two developmental stages: embryonic stage E13.5, representing a molecularly complex but still expanding spinal circuit, and post-natal and/or adult stages, proxies of a mature circuit.^42^ Paralleling our analysis in frog, we targeted six major cardinal interneuron classes – ventral excitatory V0v, V3v, and V2a; ventral inhibitory V1; dorsal excitatory dI3; and dorsal inhibitory dI6 interneurons – primary regulators of locomotor pattern in mice.^5,7^ At each stage, we either mined previously published data for rostrocaudal distributions of spinal neuron populations,^29,32,68^ or used a combination of genetically labeled Cre lines and immunochistochemistry to determine their number and settling position (**Figure 2C, S5A**). Over time, we detected a ubiquitous decrease in population size within each segment (**Figure 2D-I**). However, unlike in frogs in which rostrocaudal patterns change over development, in mice they were stable from embryonic to adult stages and notably, the same pattern was observed with antibody staining and Cre lineage tracing (**Figure 2D-I**).

This frog-mouse comparison reveals a species-conserved *Limb high* distribution for the motor-associated ventral interneuron cell types – V0v, V1, V3v (**Figure 2J-L**), in line with the hypothesis that these populations directly and proportionally shape level-specific motor output in a species-ubiquitous manner.^5,7^ However, further analysis of these classes demonstrated species-specific variation in settling position: a ventral V1 subpopulation is observed in mice, but absent from frogs (**Figure S6A-D**). This contrasted with the other classes, V0v and V3v, which were largely conserved in their position across species (**Figure S5A-D**). Furthermore, as in the frog, slight variations in settling position across rostrocaudal levels resulted in different brachial, thoracic and lumbar interneuron neighborhoods (**Figure S2C-F, S3B-E, S5B-E**). Such spatial divergence along the rostrocaudal axis and between species, even when the numerical differences between levels were conserved, suggests finer scale subtype divergence or variant connectivity could further differentiate motor function.^23,31,69,70^

The remaining cardinal classes – V2a, dI6 and dI3 – differed in their rostrocaudal numbers between species (**Figure 2M-O**). V2a interneurons were *Forelimb low* in mouse but *Hindlimb low* in the frog (**Figure 2G, 2M**); dI6 interneurons were *Forelimb low* in mouse but *Torso high* in frog (**Figure 2H, 2N**); and dI3 interneurons were *Limb high* in mouse but *Forelimb low* in frog (**Figure 2I, 2O**). Despite this, analysis of the settling position of these classes, however, showed the spatial distributions of dI3 and dI6 were highly conserved, with only V2a interneurons diverging in their location between mouse and frog (**Figure S6A-D**).

As in the frog, we evaluated the regionalization of inhibition and excitation along the rostrocaudal axis of the mouse spinal cord. We used the neuron number distributions we defined, in combination with the known projection pattern of these populations.^7^ In mice, like frogs (**Figure S4K-M**), regionalized inhibitory and excitatory circuits also emerged when the ambulatory circuit became functional (**Figure S6E**). At this stage however, uniquely in the mouse, a much larger fraction of neurons was inhibitory – totaling ∼50% (**Figure 2P-Q**). Amongst these, and in agreement with the frog, commissural inhibitory neurons were enriched in proportion at thoracic levels, whereas the ipsilateral inhibitory population was higher at limb levels (**Figure 2Q**), aligning with the differential motor output of these regions.^5^ Only in the mouse however, and not the frog, were ipsilaterally and commissurally projecting excitatory neurons both segmentally uniform in their proportions (**Figure 2P-Q**). These findings suggest conserved principles in circuit maturation across distant tetrapods that can be co-opted to differentiate and regionalize inhibitory-excitatory circuits (**Figure 2R-S**).

Taken together, these data support that rostrocaudal regionalization of the spinal cord is a fundamental characteristic of tetrapod ambulation, with a base sensorimotor circuit computation at a local level that is shared between members of the tetrapod clade. The circuit differences we identify across species additionally demonstrate the adaptability of rostrocaudal patterning within the limb-torso-limb framework that could allow for the evolutionary fine-tuning of locomotor output and sensory relay.

### Global Hox patterning of rostrocaudal neuron architecture

Our cross-species analysis of neuron architecture demonstrates rostrocaudal regionalization in spinal organization parallels regionalized function (**Figure 1J, 2R-S**). The ubiquity of interneuron regionalization raises the questions of how such differences arise, and whether they are regulated in a region-specific manner. Previous studies have shown Hox transcription factors play a key role in motor neuron specification along the body axis, with level-specific motor neurons expressing Hoxc6 at the forelimb and Hoxc9 at torso levels.^13,18,20,22^ More recently, Hox genes have also been demonstrated to play a role in rostrocaudal specification of two interneuron types in mice, V1 inhibitory and spinocerebellar neurons.^29,31,71^ We thus sought to test whether Hox genes act as global determinants of interneuron number along the rostrocaudal axis.

*Xenopus* frogs express all four families of Hox genes (a, b, c, d), including the tetrapod motor neuron thoracic-determinant *hoxc9*.^72^ As a first step to determine whether Hox gene expression drives regionalization of interneurons, we evaluated the protein expression of Hoxc6 and Hoxc9. We found Hoxc6 to be expressed in the brachial but not in the thoracic spinal cord, and Hoxc9 conversely, to be absent at brachial but present at thoracic levels (**Figure 3C**), indicating strong conservation of Hox expression between amphibians and mammals. The expression of Hoxc6 and Hoxc9 proteins was also detected in both motor and interneurons at brachial and thoracic regions, respectively (**Figure 3C**).

**Figure 3.**
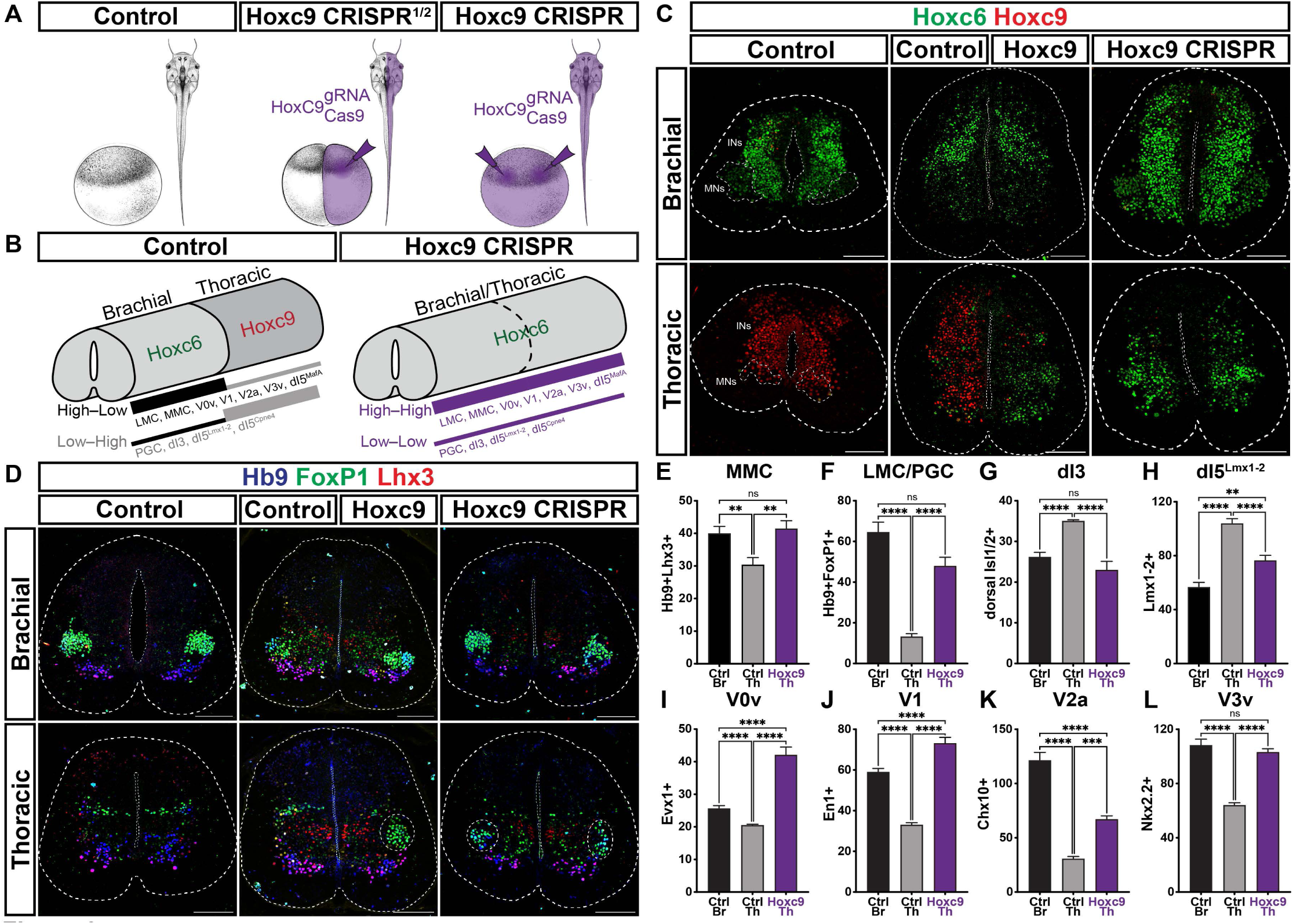
*hoxc9* loss-of-function transforms thoracic to brachial-like architecture. **A.** Schematic of CRISPR *hoxc9* loss-of-function experiment, depicting control-uninjected, and half- and whole-body Hoxc9 gRNA/Cas 9 injected animals, at one- or two-cell, and a later tadpole stage. Drawings adapted from *Zahn et al. 2022*.^182^ **B.** Summary of Hoxc6 and Hoxc9 protein expression in control and whole-body CRISPR animals, followed by a schematic summary of wild type and transformed neuron rostrocaudal distribution patterns, respectively. **C.** Thoracic-to-brachial transformation of Hox protein expression at NF55 stage frogs. Brachial Hoxc6 expression in control, half- and whole-body CRISPR animals, thoracic Hoxc9 expression only in control and the un-injected side of half-body CRISPR animals, and ectopic thoracic Hoxc6 expression in whole-body and the injected side of half-body CRISPR animals. Ventral horn motor neurons are delineated at brachial and thoracic levels of control sections based on Isl1/2^+^ expression. **D.** Thoracic-to-brachial transformation of motor neuron architecture. Brachial Hb9 and FoxP1 co-expression in control, half- and whole-body CRISPR animals marking ventrolaterally settled LMC neurons, thoracic Hb9 and FoxP1 co-expression only in control and the un-injected side of half-body CRISPR animals marking medially settled PGC neurons, and ectopic thoracic Hb9 and FoxP1 co-expression in whole-body and the injected side of half-body CRISPR animals marking ventrolaterally settled LMC-like neurons (delineated). Hb9 and Lhx3 co-expression marks MMC neurons throughout. **E-L.** Thoracic-to-brachial transformation of neuron numbers, showing motor (**E**-**F**), dorsal (**G**-**H**), and ventral (**I**-**L**) interneurons in control animals at brachial and thoracic levels, and whole-body CRISPR animals at thoracic levels of stage NF55 animals. All data reported as mean ± SEM. One-way ANOVA; ns, p>0.1; **, p<0.01; ***, p<0.001; ****, p<0.0001. For each animal three to six (n=3-6) transverse hemisections per level were scored, with N=2-14 for control, and N=3-5 for *hoxc9* mutant animals. Scale bars, 100μm. MNs, motor neurons; INs, interneurons.

Recent advances in CRISPR/Cas gene editing in *Xenopus* frogs,^73,74^ provided a high-throughput method to evaluate Hox gene function. To assess whether Hoxc9 determines the torso-level distributions of spinal neurons, we used CRISPR to target the *hoxc9* gene. By restricting our injections to either the one- or two-cell stage of the early *Xenopus* development, we generated whole- and half-body mutants, respectively (**Figure 3A**). In the thoracic spinal cord, this resulted in loss of Hoxc9 protein expression and ectopic expression of Hoxc6 (**Figure 3B-C**). Brachial expression of Hoxc6 was unaffected under all conditions (**Figure 3C**).

Given the transformation of Hoxc expression in the thoracic spinal cord of CRISPR mutant animals, we evaluated whether Hox mutation also affected thoracic motor and interneuron numbers. In *hoxc9* mutants, we observed a caudal extension of the limb-specific ventrolateral Hb9^+^FoxP1^+^ motor neuron column (**Figure 3D, 3F**), recapitulating the phenotype in mouse and chick embryos.^18,21,22^ Furthermore, this population-level analysis revealed a role of Hoxc9 in patterning axial motor neurons (MMC), which ectopically increased to brachial-level numbers (**Figure 3E**). Strikingly, a similar thoracic-to-brachial-like transformation was also evident for all interneuron populations examined: V0v, V1, V2a, V3v, and dI5^MafA^ increased; dI3, dI5^Cpne^^4^, and dI5^Lmx^^1–2^ decreased (**Figure 3F-L, S7A-D**). This analysis reveals a class-ubiquitous patterning role of Hox genes, and further supports number variation as a key principle in rostrocaudal regionalization in the spinal cord.

### Variant neurogenesis shapes rostrocaudal neuron architecture

The existence of Hox-dependent rostrocaudal patterns raises the question of what underlying developmental processes drive differences in neuron number. Molecularly distinct neural progenitors, the mitotically active precursors of mature spinal neurons, vary in their proliferative and neurogenic activity along the dorsoventral axis.^75,76^ Yet, along the rostrocaudal axis, whether neurogenesis shapes these broad-scale differences in cardinal class neuron number, and if so, what mechanisms regulate the differential proliferative potential of each class, is still unclear.

To test whether mitotic expansion underlies the establishment of regional patterns, we traced the generation of the spinal neurons stage-by-stage during frog metamorphosis using the thymidine analog EdU (5-ethynyl-2’-deoxyuridine) (**Figure 4A, S8A**). As previously shown,^57,76^ metamorphosis is characterized by a massive expansion in cell number (**Figure S9C, 9E**). Along the dorsoventral axis, the generation of ventral and dorsal neurons temporally overlapped more in frogs than in mice (**Figure S8B, S9D**), suggesting species-specific developmental timing. Along the rostrocaudal axis, cell proliferation was synchronous but varied in magnitude (**Figure S9C**), without the rostral-to-caudal neurogenesis delay described in mammals.^32,77–79^ Between limb and thoracic levels, population level analysis demonstrated differences in the timing and magnitude of dorsoventral neurogenesis by stage (**Figure S9A-B, S9E-F**), providing initial support for rostrocaudal regionalization via neurogenesis. In addition, the expansion of neuron populations coincided with an increase in progenitor domain size (**Figure S9D, S9G-I**), further implicating progenitor expansion in the regulation of neuron number.^75,76^

**Figure 4.**
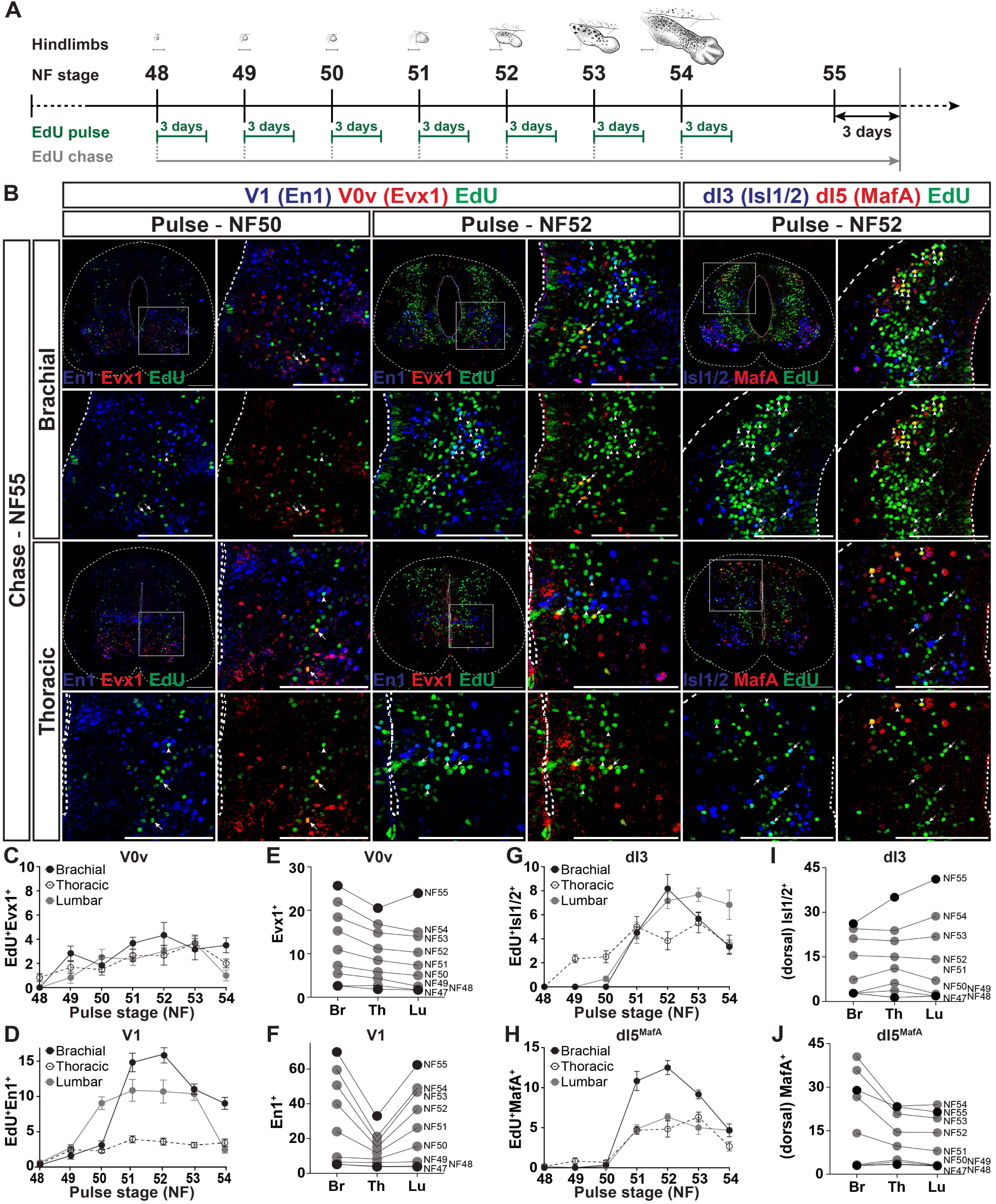
Neurogenesis diversifies and regionalizes neuron classes across rostrocaudal levels. **A.** Schematic of EdU pulse-chase experiment. Each pulse represents a 3-day injection regiment. Drawings adapted from *Zahn et al. 2022*.^182^ **B.** Different numbers of V0v, V1, dI3, and dI5^MafA^ interneurons are born at NF50 and NF52 stages at brachial and thoracic levels, marked by EdU and colabeled with Evx1 (arrows), En1 (arrowheads), Isl1/2_dorsal_ (arrows), and MafA_dorsal_ (arrowheads). Scale bars, 100μm. **C**-**D**, **G**-**H.** Number of V0v, V1, dI3, and dI5^MafA^ neurons born at each pulse based on EdU uptake. V1 and dI3 interneurons exhibit a burst-like generation pattern at limb, but not thoracic levels, V0v and dI5^MafA^ neurogenesis is uniform along the BTL axis, but time-wise stable or burst-like, respectively. All data reported as mean ± SEM. For each timepoint three (n=3) transverse hemisections per level were scored in two (N=2) animals. **E**-**F**, **I**-**J.** Regionalization of rostrocaudal patterns. Black points depict the mean for the indicated stage (as in Figure 1 or **S1**), and gray points represent the NF47 mean plus the mean of neurons born up to the indicated stage (shown in **C**-**D**, **G**-**H**).

To compare the spatiotemporal dynamics and magnitude of neuron type generation along the spinal cord, we utilized cardinal class markers in combination with EdU. We first evaluated motor neurons, tracing the addition of each motor neuron column (**Figure S10A-B, S10E-H**). Limb-innervating LMC motor neurons exhibited a burst-like neurogenesis pattern (**Figure S10C-D**). The addition of the lateral (LMC_l_) followed that of the medial LMC (LMC_m_) (**Figure S10B**), a characteristic inside-out generation sequence observed in other species.^80–83,16^ It coincided with the dorso-ventral bifurcation of the limb-invading nerve bundle into flexor and extensor divisions (**Figure S10I**),^39,84^ supporting conservation of limb-level motor subtype generation programs. Axial MMC motor neurons, in contrast to LMC, exhibited burst-like generation at brachial levels but continuous, gradual addition at lumbar and thoracic levels (**Figure S10C-F**, **S10D**), in line with graded MMC rostrocaudal distribution **(Figure S10G)**. PGC motor neurons, like the MMC, also exhibited a continuous, gradual expansion at thoracic levels (**Figure S10C-D**). The time course and mode of addition thus differed by rostrocaudal level for motor neurons, matching their regional distributions in cell number.

To determine whether heterogeneity in proliferation patterns along the rostrocaudal axis drove variations in interneuron number, we extended our EdU assay to excitatory V0v and inhibitory V1 interneurons (**Figure 4B, S11A-B**). As for motor neurons, neurogenesis of both interneuron classes varied between rostrocaudal levels. V0v interneurons were added continuously and gradually at all levels (**Figure 4C, 4E)**. Similarly, thoracic V1 interneurons were born at a steady, low rate (**Figure 4D**). Limb-level V1 interneurons instead exhibited burst-like, regionalized neurogenic expansion (**Figure 4D**), consistent with the two-fold greater limb versus torso V1 population size (**Figure 4F**).

Such distinct patterns of neurogenesis between neighboring ventral classes beg the question of whether regulation of progenitor expansion is a general mechanism of spinal neuron circuit specification. To address this question, we applied our EdU neurogenesis assay to two excitatory dorsal interneuron populations: Isl1/2^+^Otp^+^ dI3 interneurons and MafA^+^ dI5 interneurons. Both dorsal populations exhibited differences in proliferation by level. dI3 generation matched that of V1 interneurons, bursting at brachial and lumbar levels but not at thoracic (**Figure 4G**). A burst-like neurogenesis pattern was observed for dI5 interneurons at all rostrocaudal levels, however the magnitude was far greater at brachial than at either thoracic or lumbar regions (**Figure 4H**). For dI3 and dI5, like V0v and V1, interneurons, differences in generation pattern and/or magnitude between rostrocaudal levels aligned with their end-stage architecture (**Figure 4I-J**).

For other cardinal classes, we combined spatial probability EdU (**Figure S8B**) and marker gene expression maps (**Figure S2C-D**) to predict their generation pattern. We used the neuron classes that were experimentally measured as controls, confirming predicted results matched our *in situ* data (**Figure S11C-F, S11H-I**). Our predictions found variant addition patterns: V3v neurogenesis was burst-like at all levels; V2a, dI1, dI5^Lmx1+^ and dI6 generation was burst-like at limb but gradual at thoracic levels; and dI5^Cpne4+^ interneuron generation was burst-like only at forelimb and gradual at other levels (**Figure S11G, S11J**).

Taken together, these findings demonstrate that rostrocaudal differences in spinal interneuron number can be traced back to level-specific proliferation patterns, raising the question of how spinal neurogenesis is orchestrated in a level-specific manner to produce such regionalized neuronal patterns.

### Hox-dependent regulation of neurogenesis along the rostrocaudal axis via progenitor dynamics

As we have demonstrated, Hoxc9 plays a key role in determining the rostrocaudal regionalization of spinal distributions across neuron types. Progenitor domain sizes change over time,^75,76^ and correspond to neuron population expansion dynamics (**Figure S9G-I**), which means that a single regulatory cue per rostrocaudal region could result in the diverse neurogenesis patterns of different neuron populations of the same level. In mice, Hoxc9 is expressed by torso-level progenitors, and necessary for torso-specific motor neuron identity.^22^ In the ventral nerve cord of the fly, Hox genes restrict proliferation; in human stem cells and neuroblastomas, Hoxc9 likewise arrests the cell cycle.^85–90^ However, in the vertebrate nervous system, the function of Hoxc9 in progenitors is unknown. The central role neurogenesis holds in shaping rostrocaudal regionalization, along with the large differences in neuron number in *hoxc9* CRISPR mutants, raise the possibility that Hox genes actively regulate progenitors to mold regional neuron distributions in the tetrapod spinal cord.

To test this hypothesis, we examined the effects of CRISPR-mediated *hoxc9* loss-of-function on spinal cord neurogenesis using EdU pulse-chase labeling (**Figure 5A**). In the absence of Hoxc9, the proliferative output of the thoracic spinal cord was increased to brachial numbers (**Figure 5B-C**). Individual motor and interneuron populations were scored using class markers, as in wildtype animals (**Figure 5B, S12A**). Both axial MMC and level-specific LMC/PGC motor neuron generation reached forelimb levels (**Figure 5B, 5D-E**). This thoracic-to-brachial transformation of neurogenesis extended to ventral V1 (En1^+^) and dorsal dI5 (MafA^+^) interneurons, which were also born in significantly higher numbers at thoracic levels in *hoxc9* mutants (**Figure 5F, 5H, S12A**). However, mutating *hoxc9* did not uniformly cause an increase in neuron number: the Isl1/2^+^ dI3 population remained unaltered, as would be expected for a class where wildtype forelimb- and torso-level generation is the same at this stage (**Figure 4G, 5G, S12A**). These results demonstrate that the loss of Hoxc9 protein transforms thoracic into brachial-like neurogenesis in a class-specific manner (**Figure 5I**).

**Figure 5.**
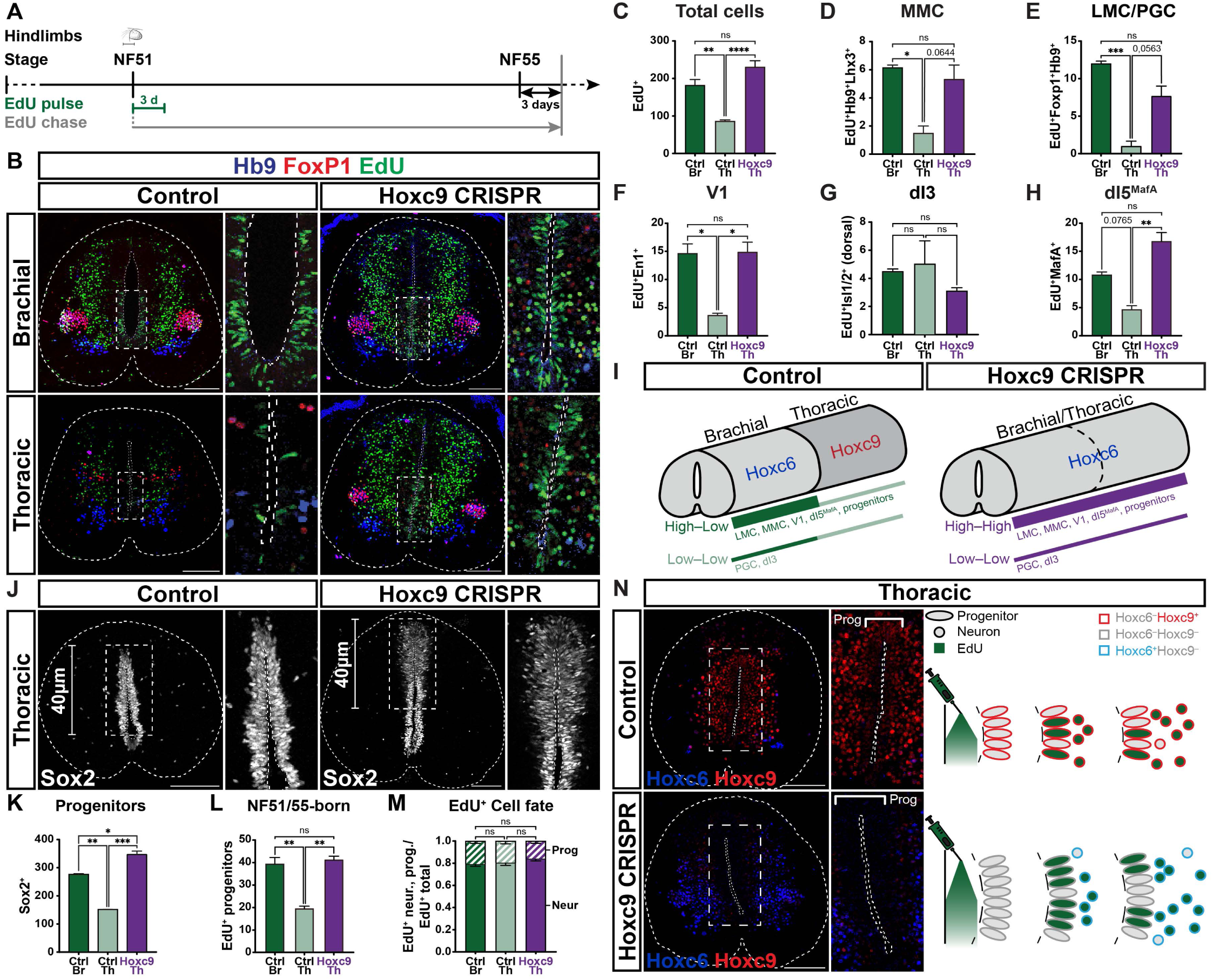
A Hox-dependent mechanism drives the regionalization of progenitors and neurogenesis. **A.** Schematic of EdU pulse-chase experiment in CRISPR *hoxc9* mutant animals. A pulse represents a 3-day injection regiment. Drawing adapted from *Zahn et al. 2022*.^182^ **B.** Thoracic-to-brachial transformation of neurogenesis, progenitor expansion and neural architecture at NF51 to NF55 stage frogs. Cells born at stage NF51 at brachial and thoracic levels in control and whole-body CRISPR animals, marked by EdU and colabeled with LMC/PGC motor neuron markers Hb9 and FoxP1. Insets depict EdU uptake in mitotically active progenitors. Scale bars, 100μm. **C**-**H.** Thoracic neurogenesis becomes brachial-like; as a whole (**C**), in motor (**D-E**), ventral (**F**) and dorsal (**G**-**H**) interneurons at stage NF51. **I.** Schematic summary of *hoxc9* loss-of-function induced transformation of neurogenesis and progenitor expansion compared to control. **J.** Transformation of thoracic progenitors in *hoxc9* CRISPR mutants compared to control animals. Insets depict progenitor expansion in Hoxc9 lacking animals. Scale bars, 100μm. **K-L.** Thoracic progenitors become brachial-like: the whole population based on Sox2 expression (**K**), and those born between NF51 and NF55 based on EdU incorporation (**L**). **M.** Cell fate determination of progenitors between NF51 and NF55 remains unchanged in *hoxc9* CRISPR mutants, calculated by dividing the number of EdU^+^ neurons and the EdU^+^ progenitors by the total number of EdU^+^ cells. **N.** Hoxc9 expression in thoracic progenitors (control) constricts both progenitor expansion and neurogenesis, while lack of Hoxc9 expression in thoracic progenitors (*hoxc9* CRISPR mutants) promotes the production of both progenitors and neurons. All data reported as mean ± SEM. One-way ANOVA; ns, p>0.1; *, p<0.1; **, p<0.01; ***, p<0.001; ****, p<0.0001. For each animal 3-6 transverse hemisections per level were scored, N=2-6 animals.

To further dissect the regulatory role exerted by Hoxc9 on progenitor function, we directly assessed the effects of *hoxc9* loss-of-function on the thoracic progenitor population. In control animals, brachial-level progenitors, as marked by Sox2 expression,^75^ were two-fold more populous than those in the torso region (**Figure 5J-K**). *hoxc9* CRISPR mutants had a two-fold increase of torso-level progenitors compared to control animals, as measured by both Sox2 expression as well as independent quantification of cells directly lining the central canal (**Figure 5L, S12B**). To directly evaluate whether Hoxc9 biases a proliferating cell to adopt a neuron or progenitor cell fate, we compared EdU incorporation in progenitors and neurons during peak limb-circuit neurogenesis. At thoracic levels, two-fold more progenitors were born in the *hoxc9* CRISPR mutants as compared to controls, indicated by the number of EdU^+^ progenitors (**Figure 5B**-insets, **5L**). EdU uptake in *hoxc9* mutant thoracic matched that of the brachial spinal cord (**Figure 5I**). However, the balance of progenitor and neuron generation was found to be stable across levels and genotypes (**Figure 5M, S12C-D**). Progenitor expression of Hoxc9 thus limits both progenitor pool size and output, without influencing the cell fate decision at the time of division (**Figure 5N**).

Taken together, these results indicate that Hoxc9 acts as a simple and robust molecular switch that constricts both progenitor expansion and neurogenesis in the thoracic spinal cord. In its absence, progenitor abundance is increased, elevating production of both progenitors and neurons at limb levels. Our findings thus expand the known roles of Hox genes in the vertebrate nervous system, and deepen our understanding of axis formation and neural patterning in the spinal cord.

### Thoracic network model predicts functional impact of spinal cord regionalization

To understand and predict how regional variations in interneurons may impact motor responses across species, we constructed two spatially embedded network models of the frog and mouse spinal cord (**Figure 6**, **S13, S14**). For both species, a wildtype spinal cord volume was populated with ventral cell types MMC, V0v, V1, V2a, V2b, V3v and dI6, and also V0d in mouse, in the proportions and spatial positions found at thoracic levels at NF55 in frog and E12.5-13.5 in mouse (**Figure 6A-B**, **S14A-B,** *also see* **STAR Methods**). In both models, we then sampled synaptic connections by assigning mouse-like spatial projection rules to each cell type, based on recent advances demonstrating how spinal network rhythms emerge from asymmetric connectivity patterns along the rostrocaudal axis (**Figure 6C**, **S13A-B**).^91^ This yielded network matrices we could analyze in a conventional firing-rate model framework (**Figure 6D-E**, **S14C**, *also see* **STAR Methods**). To investigate thoracic motor responses using our models, we utilized two different stimulation paradigms to elicit network responses. First, to evaluate the rhythm and pattern generation capacity of the networks, we applied a global tonic input to all spinal neurons and measured motor output. We found that both networks fired synchronously on the left-right side in rostral-to-caudal propagating waves (**Figure 6G-H, S14E-F, S14H**), despite differing in their regional composition, length, and thoracic root segmentation.

**Figure 6.**
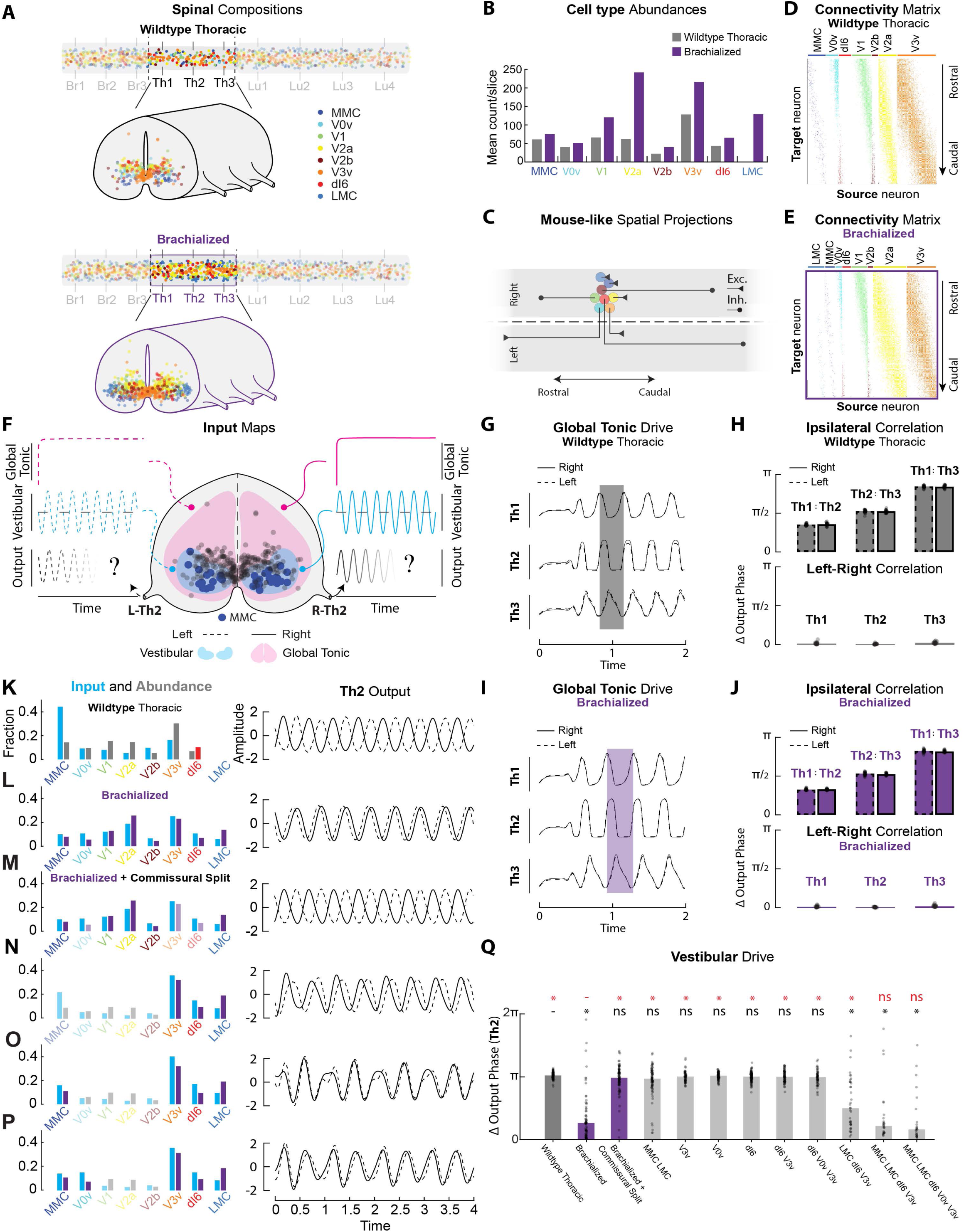
Hoxc9 thoracic network model predicts shift from alternating to bilateral coupling of vestibular reflex. **A.** Schematic of thoracic wildtype and brachialized model volumes, populated with eight ventral cell types. Compared to wildtype (black, top), the brachialized model (purple, bottom) integrates our experimentally mapped changes to the numbers and spatial regionalization of the ventral types. **B.** Mean slice abundances for each cell type for the two models. Brachialized cell types are denoted with purple bars. **C.** Assigned mouse-like spatial projection biases for each cell type along the rostrocaudal axis, independent of Hoxc9 genotype. Axon terminals signify the mean projection distance along the relative distance space, where the synapse probability is the highest. Around this point, synapse probability decays Gaussianly. **D, E.** Connectivity matrices for the wildtype (top) and brachialized (bottom) networks, resulting from combining the cell type composition and projection biases for the two models. Source neurons (columns) are sorted according to cell type identity. Target neurons (rows) are sorted caudo-rostrally. **F.** Transversal view of the wildtype model with highlighted MMCs (dark blue, enlarged). Remaining cells in gray. Two input modalities are illustrated: Tonic global (pink) and vestibulospinal (light blue) drive. In the input/output curves, dashed and solid denote left and right hemicords, respectively. Network response was measured through the middle thoracic roots, L-Th2 (left) and R-Th2 (right). **G.** Example steady state activity of the three ventral roots in the wildtype thoracic (**H**) and brachialized (**I**) network in response to global, tonic drive as illustrated in **F.** Width of shaded rectangle corresponds to a single oscillatory period. **H.** Phase correlation between steady state rhythms along and across the cord in the wildtype thoracic network in response to global tonic drive. N=20 instantiations for each network. The rhythm is left-right synchronous but propagates rostro-caudally along the cord. **I.** As in **G**, but for the brachialized network. The rhythm is also left-right synchronous in the brachialized network and propagates rostro-caudally along the cord. **J**. As in **H**, but for the brachialized network. **K-P.** Cell type and vestibular input composition for model networks along with example output traces (median network sample, as in **G**). Blue bars denote fraction of vestibular inputs received by the cell type in question, colors (purple) denote wildtype (brachialized) population fraction of the cell type in question. Th2 output amplitude in arbitrary, standardized units (left: dashed, right: solid). **Q.** Distribution of left-right phase differences for wildtype thoracic, fully and partially brachialized model networks in response to vestibulospinal input. Bar height denotes median. Statistical comparison between distributions: Mann-Whitney U Test with bonferroni corrected α = 0.0045. Wildtype:Condition notation (* or ns) in black. Brachialized:Condition notation in red. Wildtype networks (dark gray) maintain alternating responses to alternating vestibular input (Δθ=3.19 rad) compared to predominantly left-right synchronized responses (Δθ=0.82 rad) in brachialized networks (purple) (p < e-23). Commissural splitting in brachialized networks re-establishes alternating response (Δθ=3.08 rad, p_Wildtype : Commissural Split_ > 0.1, p_Brachialized : Commissural Split_ < e-20). Brachializing LMCs, MMCs, V3vs and dI6s together is sufficient to induce the synchrony response seen in the brachialized model (p_Wildtype : [LMC MMC V3v dI6]_ < e-10, p_Brachialized : [LMC MMC V3v dI6]_ > 0.8). Remaining wildtype thoracic comparisons p_Wildtype : x_ : [_LMC MMC_, p>0.006; _V3v_, p>0.01; _V0v_, p>0.4; _dI6_, p>0.02; _V3v dI6_, p>0.1, _V3v V0v dI6_, p>0.01; _LMC V3v dI6_, p<e-11; _MMC LMC V3v dI6 V0v_, p<e-10]. Remaining brachialized comparisons p_Brachialized : x_ : [_LMC MMC_, p<e-17; _V3v_, p<e-17; _V0v_, p<e-23; _dI6_, p<e-22; _V3v dI6_, p<e-19, _V3v V0v dI6_, p<e-17; _LMC V3v dI6_, p<0.002; _MMC LMC V3v dI6 V0v_, p>0.4]. n is the number of data points out of 100 instantiations.

Next, to evaluate the ability of the network to recapitulate local thoracic behaviors, we took advantage of the vestibular righting reflex, a motor function that selectively engages the thoracic network to attain balance in both frogs and rodents.^37,92–94^ In the model, we applied a local, alternating stimulus to the input region of the vestibular tract. This input region was defined here as the region occupied by MMC motor neurons and surrounding interneurons, based on anatomical tracing and synaptic connectivity mapping.^37,93,95^ It was constructed as a transversal probability map based on the distribution of MMC neurons, resulting in variant fractional inputs to each neuronal class (**Figure S2D, 6F, S14D**). Network output, measured at the mid-thoracic root, was found to be left-right alternating: root output was in-phase with its ipsilateral input sinewave and anti-phase with the contralateral root output (**Figure 6K, 6Q, S14G**), in accordance with the cross-coordination of the vestibular righting reflex.^37,94,96,97^ These two spatially-faithful networks of frog and mouse spinal cord can thus, with comparable stimuli, robustly generate two stereotyped activity patterns *in silico*.

Equipped with these models, we were now positioned to test how changes in thoracic network composition, as observed in Hox mutant frogs, changed network output. Loss of Hoxc9 induces a brachial-like transformation of the thoracic spinal cord. To simulate this transformation, we populated a mutant - or “brachialized” - thoracic volume with the numerical and spatial ventral cell type compositions we uncovered at brachial levels in developing frogs (**Figure 6A**, **6B**). Tonic input to this brachialized network still drove synchronous, wave-like activity as in the wildtype thoracic network (**Figure 6I-J**), indicating baseline pattern generation was unaffected. However, when a vestibulospinal stimulus was applied to the same input region in a brachialized as a wildtype network, the fractional input to each neuron population correspondingly changed and the correlation between the left and right output was dramatically altered (**Figure 6F, 6L, 6Q**). The activity was predominantly bilaterally synchronous instead of alternating (**Figure 6L, 6Q**). This transformation from alternation to synchrony led us to examine which network features respecified output, an investigation which is difficult to execute *in vivo*.

The most obvious candidate neuronal populations to regulate left-right coordination are the commissural interneurons.^5,7,98^ To probe the impact of commissural projections on network synchronization, we removed the commissural synapses in the brachialized network. This manipulation reestablished the alternating response of the wildtype model (**Figure 6M, 6Q**). Moreover, synchrony could also be achieved with only the brachialization of commissural projecting populations and motor neurons (**Figure 6P, 6Q**). To identify the source of the synchronization effect in the brachialized network, we made partial cell-type specific brachializations. Surprisingly, no single brachialized cell type was sufficient to induce a synchronized response (**Figure 6Q, S13D-F, S13I**). Additionally, neither any two-type combination of motor or interneurons, nor any combination of interneurons alone, reproduced the phenotype (**Figure 6Q**, **S13G-I**). Simply brachializing motor neurons was also insufficient to cause a synchronous response (**Figure 6Q, S13C**). However, a divergence from the wildtype thoracic response was observed when a motor neuron population was transformed together with the V3v and dI6 classes (**Figure 6N**, **6Q**, **S13I**). Further brachializing both MMC and LMC motor neurons in combination with V3v and dI6 best recapitulated full-circuit brachialization (**Figure 6O**, **6Q**). Thus, our model predicts that differences in rostrocaudal circuit output are dictated by the number and distribution of, not one, but combinations of spinal neurons.

Our computational work predicts that loss of Hoxc9 and subsequent brachialization of thoracic networks induces a population-level synchronization effect on the vestibulospinal response, supporting a central role of neuron distributions in influencing regional output.

### Hoxc9 expression defines thoracic morphology and function

To test experimentally whether thoracic function is influenced by changes in rostrocaudal neuron distributions, we performed ventral root recordings on wildtype and *hoxc9*-mutant frog spinal cords, in which the thoracic neuron number and distribution was brachialized **(Figure 3**). Post-metamorphic, wildtype *Xenopus* frogs have three thoracic roots on either side of the body that innervate somatic muscles and viscera (**Figure 7A**).^99^ Dissection in preparation for electrophysiological recordings revealed a dramatic anatomical change: in *hoxc9* mutants, the first and third thoracic roots (Th1 and Th3) were redirected to the fore-and hindlimb plexus, respectively (**Figure 7B**). This indicated a complete Hox-dependent transformation of the Th1/3 thoracic motor neuron projection pattern, previously suggested in Hoxc9 mice but not observed to this extent.^22^

**Figure 7.**
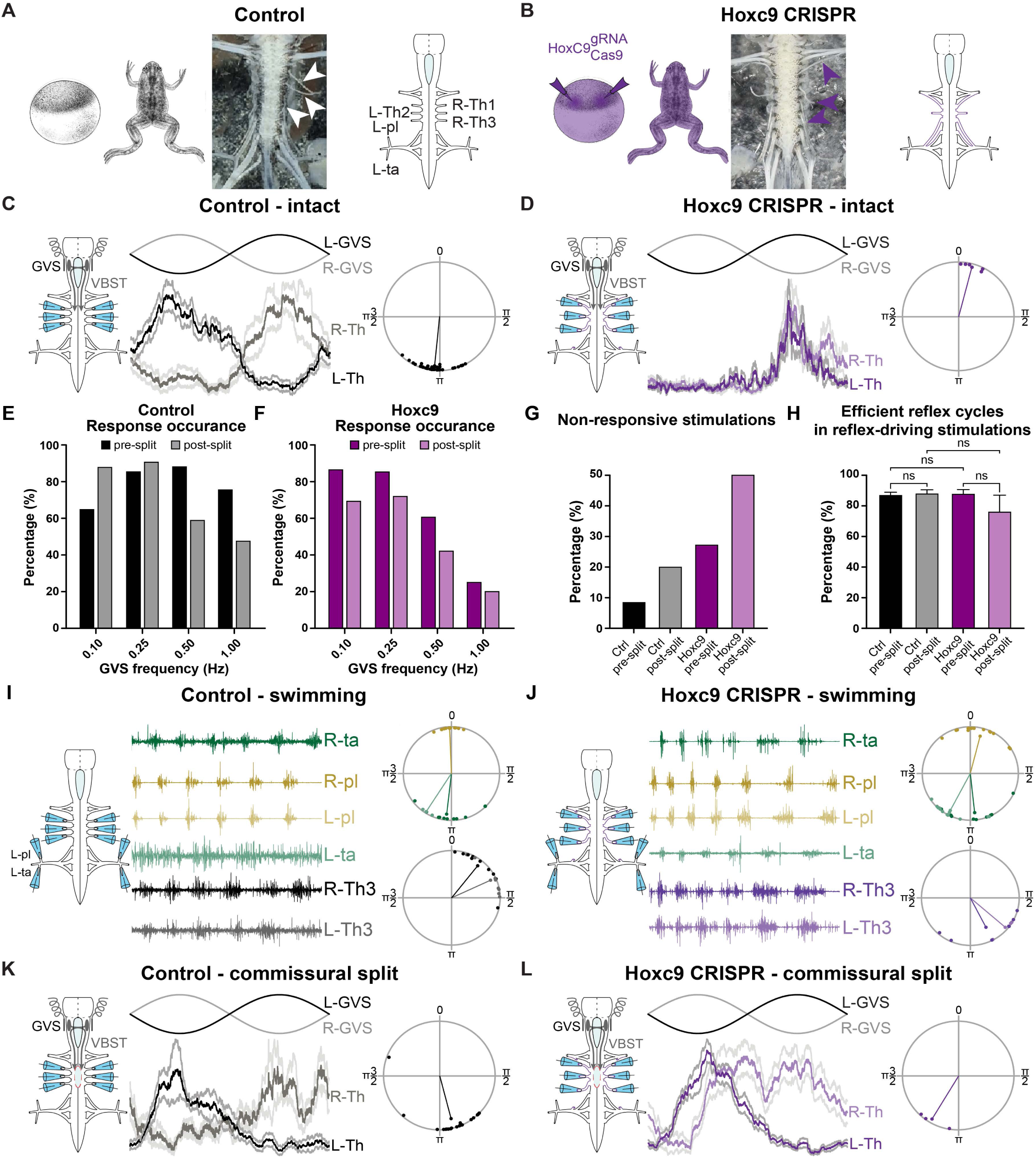
Hoxc9 transformation of electrophysiological output recapitulates model predictions. **A, B.** Control and whole-body *hoxc9* CRISPR animals with anatomical preparations and schematics, highlighting the three thoracic roots (arrowheads), where Th1 and Th3 in *hoxc9* CRISPR animals are re-directed to the brachial and lumbar plexus, respectively (**B**). Drawings adapted from *Zahn et al. 2022*.^182^ **C**, **D.** Alternating galvanic stimulation (left-right) of the vestibular nucleus in the hindbrain causes alternating (right-left) in control (**C**), but synchronous thoracic root responses in *hoxc9* CRISPR animals (**D**). Data shown for Th2 with 0.5 Hz GVS as mean ±SD root recordings, and phase difference in polar plots between left-right. For both control and *hoxc9* CRISPR animals, N=6. Polar plots, control: ma=185.9±3.1°, r=0.99, Rayleigh test p=0.001; HoxC9: ma=14.6±6.3°, r=0.99, Rayleigh test p=0.002. **E, F.** Percentage of galvanic stimulation frequencies that triggered a response in control (**A**) and *hoxc9* CRISPR mutant (**B**) animals, pre- (dark shade) and post-split (light shade) of the thoracic spinal cord commissural projections. Only animals with both pre- and post-split responses were included, N=5 for wildtype and N=7 for mutant animals; and only galvanic stimulations that did not evoke locomotor-like hindlimb responses/activity were included in the analysis, all three thoracic root pairs (L-/R-Th1, L-/R-Th2, L-/R-Th3) were sampled resulting in n=294 distinct GVS sequences for wildtype and n=279 for mutant animals. **G.** Percentage of stimulations that failed to trigger a response in control (black) and *hoxc9* CRISPR mutant (purple) animals, pre- and post-split of commissural connections. All three thoracic root pairs (L-/R-Th1, L-/R-Th2, L-/R-Th3) were sampled. **H.** GVS-evoked responses successfully triggered thoracic reflex bursts equally pre- and post-split, in both control (black) and *hoxc9* CRISPR mutant (purple) animals. The percentage of successful reflex cycles in stimulation sequences that evoked reflex responses are shown for three thoracic root pairs (L-/R-Th1, L-/R-Th2, L-/R-Th3) at both 0.25 and 0.5 Hz GVS. Data reported as mean ± SEM. **I**, **J.** Root recordings and phase differences of hindlimb flexor-extensor rootlets (R-/L-ta, right/left *tibialis anterior*; R-/L-pl, right/left *plantaris longus*) show normal swimming, but strikingly swim-evoked thoracic response (R-/L-Th3, right/left 3^rd^ thoracic root) varies between control (**I**) and *hoxc9* animals (**J**). Raw nerve and root recordings, and phase difference in polar plots against the left *plantaris longus* are shown. N=3 and N=5, for control and *hoxc9* CRISPR animals, respectively. **K**, **L.** Following a surgical split of the thoracic commissural projections, alternating galvanic stimulation (left-right) of the vestibular nucleus causes alternating (right-left) responses in both control (**K**) and *hoxc9* CRISPR animals (**L**). Data shown for Th2 with 0.5 Hz GVS as mean ±SD root recordings, and phase difference in polar plots between left-right. N=5 and N=2, for control and *hoxc9* CRISPR animals, respectively. Polar plots, control: ma=165.1±31.7°, r=0.86, Rayleigh test p<0.001; HoxC9: ma=212.0±15.0°, r=0.97, Rayleigh test p=0.046. GVS, galvanic stimulation; VBST, vestibulospinal track

We capitalized on the frog’s vestibulospinal reflex to directly measure the effects of *hoxc9* brachialization on thoracic function. In *Xenopus* frogs, natural stimulation of the vestibular horizontal canals evokes a bilaterally alternating thoracic reflex response.^37^ To mimic this response, in both wildtype and *hoxc9* mutant frogs, we excited the vestibulospinal tract using left-right alternating galvanic vestibular stimulation (GVS).^96^ In wildtype animals, right-side stimulation triggered left-side responses; whereas left-side stimulation triggered right-side responses (**Figure 7C**). This thoracic GVS-evoked alternating reflex thus consisted of left-right bursts, phase-locked to the stimulus sinewave and in anti-phase with each other (**Figure 7C**). In striking contrast, in *hoxc9* mutant animals, the thoracic responses were bilaterally synchronous, locked to the same GVS phase and in phase with each other (**Figure 7D**). Furthermore, in *hoxc9*-mutant thoracic networks compared to wildtype, lower frequencies of GVS stimulation were more efficient in evoking responses than higher frequencies, and there was a two- to three-fold higher failure rate of stimulus-evoked responses (**Figure 7E-H**), showing that the responsiveness of the circuit was tied to the transformation of neural architecture. Overall, as predicted by our computational model, Hoxc9 spinal reorganization produces a functional network that is bilaterally synchronized, as opposed to alternating, in response to vestibular stimulation.

During swimming, thoracic networks receive ascending input from the lumbar spinal cord to coordinate hindlimb kick bouts and postural control, which is necessary for proper locomotor behavior.^100^ To test the effects of Hoxc9-driven thoracic transformation on swimming, we performed electrophysiological recordings concomitantly from thoracic ventral roots and the nerves innervating individual hindlimb flexor and extensor muscles. Under normal conditions, hindlimb flexor and extensor -innervating roots alternate, and thoracic-innervating roots fire with a short delay.^47,94^ During such swimming bouts, *hoxc9* mutants, like wildtype frogs, exhibited highly coordinated flexor/extensor alternating responses (**Figure 7I-J**), suggesting that hindlimb networks were resistant to the *hoxc9* mutation and consistent with the specific transformation of the thoracic spinal cord. However, coordination of lumbar and thoracic circuits, as measured by the delay of thoracic bursts following extensor firing, was perturbed, with a greater delay in *hoxc9* mutant as compared to wildtype frogs (**Figure 7I-J**). These findings show that the Hoxc9-driven thoracic transformation broadly affects rostrocaudal networks, not only for sensory-motor but also for locomotor networks.

Our model predicts that aberrant commissural projections are necessary for the synchronization of vestibular-evoked thoracic responses. To directly test this hypothesis, we surgically split the thoracic spinal cord longitudinally from Th1 to Th3 in both wildtype and *hoxc9* mutants. Severing the commissural projections did not change the alternating nature of the GVS-evoked reflex response in wildtype animals (**Figure 7K**). In *hoxc9* mutants, however, it largely restored left-right alternation (**Figure 7L**). Following commissural split in both wildtype and *hoxc9* preparations, GVS stimulation in general failed to evoke a response as reliably, and responses became more variable (**Figure 7G, 7K-L**). This suggested that the thoracic circuitry as a whole actively participates in vestibular-driven postural reflexes, with local bilateral interactions lowering the threshold for reflex expression and shaping its output. Together, these data demonstrate that synchronicity resulting from the Hoxc9 transformation is mediated by a change in the balance of commissural excitation/inhibition.

Our electrophysiological findings thus indicate that loss of Hoxc9 results in a dramatic morphological and functional transformation, validating our model’s predictions.

## DISCUSSION

The limb-torso-limb regionalization of the vertebrate *bauplan* saw an explosion of locomotor patterns, but how changes in spinal interneuron architecture correspondingly diversified behavior is surprisingly still unknown. We found that frog and mouse interneuron circuits are rostrocaudally regionalized, exhibiting both species-conserved and -divergent local features that align with their movement patterns and developmental stage. We demonstrate that these neural distributions are orchestrated on a progenitor-level by a Hoxc9 on/off switch. They play a key role in shaping regional output, as their disruption perturbed both descending reflex and locomotor responses. Our findings thus provide an axis to uncover the developmental, cellular and functional basis of motor diversity produced by spinal circuits through the lens of an organism’s body plan.

### Regionalized spinal architecture mirrors body plan and function

Despite decades of work on the spinal cord and its component neurons, it still remained unclear whether differences in the number and distribution of interneurons according to body region is a ubiquitous feature of spinal circuits and how these differences might influence regional motor output. Here, we establish: (i) early tadpole swim circuits are uniform in their rostrocaudal architecture; (ii) regional differences arise with the emergence of limbs during frog metamorphosis; and (iii) four-limb vertebrate species have both species-conserved and -divergent neuron distribution patterns across rostrocaudal levels.

Simple undulatory swimming, requiring rostrocaudal wave propagation, thus arises from a largely even distribution of excitatory and inhibitory neurons.^63,64,101–105^ In such a circuit, rostrocaudal biases in projection pattern shape network dynamics, as observed in lamprey, larval zebrafish and tadpoles.^5,14,62,64,106–109^ While this axial architecture is the primary contributor to locomotion, regionalized circuits in zebrafish control the pectoral fins.^110–112^ In addition, in the cartilaginous little skate, fin regions were expanded and further specialized at pectoral levels for synchronous, and at pelvic levels for alternating, movement.^2^ And, as we demonstrate here, frog metamorphosis exemplifies that a uniform axial distribution must become regionalized for limb-locomotion.

Until now however, it was unclear whether the addition of limbs and the diversification of motor neuron pools to innervate their musculature required a new compartmentalized, regional interneuron circuit. Supporting the existence of local microcircuitry, early transplantation experiments in chick demonstrated swapping of lumbar with brachial spinal segments was sufficient to change the pattern of locomotion from alternating to synchronous.^113^ The cellular and circuit differences that underlay this functional transformation however were still unknown. More recently, rabies tracing experiments from fore- and hindlimb musculature found the majority of premotor networks were local,^114–116^ with the exception being propriospinal networks which connect the two limbs.^117^ Our rostrocaudal atlas of spinal interneurons shows regionalization of the spinal cord is a ubiquitous feature of four-limb circuits that does not simply match the enlargement of the spinal cord at limb levels.

Interneurons that closely regulate motor neuron output,^5,7,8^ namely V0v, V1 and V3v, were enriched at limb levels, arguing for an optimal regulatory-to-motor neuron ratio per class – a rhythm generation unit for limb movement conserved across species. Similarly, our finding that dI1 neurons were higher in number at limb levels, aligns with the dI1 population comprising a proprioceptive circuit that predominantly relays positional information of limbs to the cerebellum, with the forelimb bias perhaps reflecting a tighter coordination of hand-to-mouth feeding.^118–121^

Dorsal sensory-related interneurons deviate from the high-limb, low-torso patterns observed for motor neurons and motor-related interneurons. For dI5 interneurons, a high-brachial pattern is observed for the MafA subpopulation, implicated in mechanosensation,^69,122^ which could facilitate forelimb feeding and exploration via sensory-motor networks across species.^34^ A clear thoracic enrichment however was observed for the larger, parental Lmx1b/Cpne4 dI5 class. Given the reported role of dI5 neurons in transmitting somatosensory input ^56,69,123–125^, such enrichment may be due to prioritization of the trunk containing vital organs, or the convergence of sensory signals in the thoracic spinal cord to coordinate cross-level locomotor responses. Neuron class numbers can thus spatially mirror region-specific functions.

Three interneuron populations – the commissural inhibitory dI6 and the ipsilateral excitatory V2a and dI3 – differed in their rostrocaudal patterns between both tadpole/frog stages and frog/mouse. dI6 interneurons coordinate left-right alternation in mice and V2a interneurons regulate the rostral-to-caudal propagation of tail circuit activity in zebrafish.^32,61,126,127^ In tadpoles, assuming these functions are conserved, the rostral enrichment of both populations could facilitate the initiation and caudal propagation of alternating waves to drive tail undulation.^47,128,129^ This function would be unnecessary at frog stages: with the tail now lost and limbs driving locomotion, this enrichment is no longer observed.^47,129^ In addition, unlike in frogs in which synchronous hopping dominates, in mice hindlimb alternation is prominent, for which high lumbar dI6 and V2a populations are essential.^32,51,68,126,130^ Moreover, mice involve all four limbs in their gaits and are characteristically dexterous,^5,8,66^ while *Xenopus* frogs rely mainly on their hindlimbs for locomotion and exhibit only crude forelimb functions.^44,119,131^ The proportional enrichment of dI3 neurons in the mouse forelimb region, compared to the frog, could reinforce these species-specific differences in fore-hindlimb coordination and finer grasp control.^23,24,132,133^ Finally, while the highly conserved molecular identity of cardinal classes indeed supports such conserved functions across four-limb species,^43,131^ more subtle differences in gene expression or connectivity between cardinal subtypes, consistent with the spatial differences we observe for V1 and V2a subpopulations, might also confer species-specific functions.

The degree of overall similarity in spinal architecture between mouse and frog, despite these species-specificities, supports a prominent conserved role for individual neuron types in locomotion across distantly related species. Within this however, variation in regional distributions of otherwise conserved neuronal populations is a ripe source of locomotor diversity, and a previously underappreciated foundation for resolving circuit evolution.

### Variation in the Timing of Neurogenesis

Previous studies have described spinal neurogenesis as exhibiting a characteristic caudal lag, with brachial developing earlier than lumbar regions.^77,78,134^ In species like the mouse, spinal neurogenesis happens in tandem with rostral-to-caudal organogenesis.^77^ In the *Xenopus* frog however, limb-circuit neurogenesis takes place within an already formed spinal cord, allowing us to examine the mechanisms driving regionalization independent of neural tube development.^39,57^ Indeed, we found that neurogenesis, separated from any embryonic directional pull, proceeds in synchrony across all rostrocaudal levels.

Yet, despite the broad temporal synchronization of limb regions, the thoracic spinal cord consistently diverged. Considering that birthtiming of spinal neuron classes has been previously linked to subtype differentiation,^25,32,135^ our findings provide a potential developmental basis for torso-specific subpopulation specification. We further showed that the sequential progression of neurogenesis – first motor neurons; then ventral; followed by dorsal interneurons – described in mice and rats –^77,78,136,137^ was less pronounced in the frog. Given the lower neuron numbers of the developing frog spinal cord and its characteristically wider neurogenic window, the punctuated waves of neurogenesis of the mouse spinal cord could thus represent an evolutionary adaptation to underlying spatial or metabolic constraints.^138–140^

Our study additionally reveals neurogenesis of the limb-innervating LMC motor neurons was synchronized in *Xenopus* between brachial and lumbar levels in its initiation, cessation and spatial order – LMC_m_ before LMC_l_. However, in frogs, forelimb development and function are delayed relative to that of the hindlimb.^39^ This disconnect between limb and spinal cord development thus suggests independent neuronal and mesodermal clocks. This is further supported by limb and spinal cord transplantation experiments in both frog and chick, where despite the resulting temporal misalignment limb innervation was still functional.^141–148^ Interestingly, parallel Hox codes have been described for the central nervous system and the neighbouring mesoderm,^18,22,149–151^ conferring positional information in both tissues and perhaps ensuring the formation of proper neuromuscular units,^22,151^ rather than aligning the developmental timing across tissues.

Neurogenesis is thus an essential means of central nervous system regionalization. Within this framework, each neuron population exhibits a distinct spatiotemporal birth signature, resulting in a unique combinatorial pattern for each rostrocaudal region.

### Hox genes regionalize spinal neurogenesis

As neurogenesis is a key means to regionalize the spinal cord, it must be regulated with high precision. Yet, while previous studies have highlighted the involvement of Hox genes in rostrocaudal patterning,^9,152^ their potential role in patterning neurogenesis has received comparatively little attention. We found that this role was realized by amplifying progenitor pool size and cellular output in a switch-like manner. In this way Hoxc9 expression, or lack thereof, generates the right number of neurons, at the right region and at the right time – a prerequisite for their subsequent molecular and thus functional subdivision. Hoxc9-mediated downregulation of progenitor expansion and neurogenesis at thoracic levels could be due to a combination of: (i) decreased proliferation rate; (ii) increased progenitor cell death; and (iii) earlier cell cycle exit. Indeed, while multiple studies have shown Hoxc9 to directly restrict proliferation,^88–90^ it has also been implicated in driving both apoptosis and differentiation.^88,90^ In the spinal cord, the absence of genetic redundancy for such a conserved and crucial process marks *hoxc9* as a critical regionalization tool and a prime evolutionary target,^2,9,21^ but also suggests an underlying robust regulatory network.

While the conservation of Hox gene expression across vertebrates and of rostrocaudal patterning of motor neurons and V1 interneurons is well established,^2,9,21,22,29,153^ here we discovered a near-ubiquitous role of Hox genes in patterning interneurons. Based on these data, we suggest a determinant role of Hoxc9, and by extension of Hox genes in general, on all interneuron distributions along the rostrocaudal axis that applies broadly across vertebrates. Such a role has also been proposed for invertebrates: in the fly larvae, differences in the neuronal expression of Hox genes have been linked to changes in circuit architecture and in turn, both neuronal function and behavioral output.^154,155^ Our study in frogs also exemplifies the potent role Hox-dependent neuronal specification has not only on the architecture of the spinal cord but also its function, as we demonstrated both *in silico* and *in vitro*. We found Hox-dependent neuronal changes specify regional circuit output. Hox genes thus provide a cross-phyla mechanism to parcellate circuit function via region-specific changes in not only motor neurons but also regulatory interneurons.

### Neuronal basis of circuit output

Whereas previous studies have highlighted the molecular diversity of spinal neurons and the role of Hox genes in fate determination,^15,16,18,20,29,31^ the functional implications of neural regionalization were an open question. Our study found that the *hoxc9* mutation induced such a pervasive phenotype that thoracic ventral roots were redirected from the torso muscles to the proximal limbs. Electrophysiological recordings from all three thoracic root pairs were equally aberrant. These data demonstrate that while target-finding may be crucial for sustaining motor neuron numbers,^144,156^ it does not determine motor output. Hox manipulation thus results in a network-level transformation of circuit function.

Computational models have had a central role in helping neuroscientists understand diverse motor output.^157–160^ Here, we adopted a recently proposed theoretical framework, wherein spinal motor rhythms emerge from population-level interactions within spatially distributed recurrent networks organized by probabilistic connectivity rules.^161–163^ A strength of this approach is that it allows us to recapitulate dorsoventral and rostrocaudal differences in circuit architecture, and model the continuous and flexible integration of motor demands characteristic of multifunctional, often regionalized, spinal networks.^91,161^ The same spatial and network principles which underlie flexible lumbar locomotor dynamics in a mouse model,^91^ when combined with thoracic neuronal distributions, captured thoracic motor responses in mouse and frog. In both species, different stimuli efficiently generated specific, stereotyped output, further reinforcing the versatility of this method. In addition, global tonic drive, in both species’ models, generated rostral-to-caudal propagating waves. Such caudally-directed waves have been found in electrophysiological recordings of newborn rat thoracic spinal cord during fictive locomotion.^164,165^ The mammalian thoracic spinal cord, as we observed in our model, was also capable of producing synchronous responses *in vitro*^166,167^, although these were overridden by lumbar alternation during fictive locomotion.^164,165,168^

Interaction between the divergent neuronal abundances and regional distributions conferred by the loss of Hoxc9 also transformed thoracic output within our model, matching our electrophysiological data. Proportional variance across thoracic networks with the same projection motifs affected the output by altering connectivity and consequently, emergent dynamics. At a cell type level, changing just the number and distribution of only three neuron classes was sufficient to change motor output. In dictating motor output, this suggests that more granular features such as subtype identity or strict, non-probabilistic connectivity rules play a secondary role to cardinal neuron distributions and their broad projection patterns.^29,32,61,63,64^ Spatial differences can affect motor output by altering the neurons targeted by descending inputs,^30^ as descending tracts tend to invade predetermined areas independent of the transversal cell distribution.^169,170^ In brachialized networks, descending drive was predominantly distributed over V2a and V3v neurons compared to wildtype networks, in which motor neurons were mainly recruited. Together, these spatial network phenomena constitute an integrated mechanism by which spinal regionalization affects motor dynamics, despite the uniformity of single neuron response properties we assumed here. In addition, the physiological appropriate responses we observed in our frog model, which has the connectivity of a mouse circuit, predict species-shared projection and connectivity patterns of cardinal classes and descending vestibular input, both experimentally testable and partially supported by shared ipsi- and contralateral projections of cardinal populations across species.^7,14^

On a locomotor level, vestibular-to-thoracic dysfunction can greatly affect *Xenopus* behavior.^94^ As such, the *in vitro* and *in silico* transformation of thoracic output we observe in *hoxc9* mutants, along with the potential loss of trunk and axial muscle innervation due to the redirection of thoracic roots, could impair balance, posture, breathing and even locomotion itself. Furthermore, it has been argued that lack of Hoxc9 expression plays a fate-determinant role,^2,22^ and could thus be sufficient to produce neurons primed to respond to limb-derived, axon guidance cues that direct motor axons to the nearest limb. Alternatively, signals from the neighbouring limb levels,^171–173^ might differentially influence their adjacent Hoxc9-lacking regions endowing them with a fore- or hindlimb bias. Given the intrinsic propensity of limb-innervating motor neurons to establish functional synapses with limb muscles, even on the opposite side of the body, and their dependence on such synapses for their survival,^146,147,174,175^ it is likely that the ectopic LMC-like, thoracic motor neurons synapse onto fore- and hindlimb muscles. Hoxc9 mutation could thus interfere with normal locomotion and feeding.

### Neural regionalization and the diversification of circuit output

Our findings show that studying neural regionalization is paramount to understand circuit output. This raises questions about which core principles shape regional output, and how development locally diversifies not just cell type but also circuit architecture. We find that spinal regions are in many respects – developmentally, cellularly, and even functionally – separate units, yet they coordinately generate complex behaviors. Our study identifies regulation of neurogenesis via Hox genes as a simple yet versatile tool for both development and evolution to generate these regions and mold local circuits. Such a generalizable principle for local diversification of neural output via regional differences in cell number and distribution can be applied more broadly to other areas of the CNS, from the hindbrain to the motor cortex.^9,176^

### Limitations of the study

Neuron architecture in the spinal cord was largely characterized using immunohistochemistry for canonical markers of each neuron class. This approach was employed due to the fact that Cre-lines to lineage trace each population are not available across species. To account for this, the choice of embryonic and tadpole developmental stage was aligned with the timepoint of maximal marker gene expression,^42,43,131^ and notably, in mice, the same rostrocaudal pattern was observed between developmental and adult stages (**Figure 2**). In addition, in mice when possible, both Cre-lines and antibody staining were utilized and similar patterns were observed with both approaches (*see* **STAR Methods**). A caveat however to our approach is that markers may be downregulated and/or mark different neuron types across species, although cross-species single cell analysis argues against this possibility.^43^

We used first-generation (F0) CRISPR-generated mutant *Xenopus laevis* animals to examine the molecular, developmental and functional effects of *hoxc9* loss-of-function (**Figure 3, 5, 7**). In frogs, F0 CRISPR is a highly efficient method to generate whole-body mutation.^131,177,178^ For *hoxc9*, we show that mutation coincides with near total loss of Hoxc9 protein expression and gain of ectopic Hoxc6 expression and furthermore, that *hoxc9* CRISPR mutant frogs have the same motor neuron phenotype as Hoxc9^-/-^ mutant mice (**Figure 3**).^22^ The phenotypes we observe are also of high penetrance and expressivity, the misdirection of the ventral roots was even more severe in Hoxc9 mutant frogs than in mutant mice (**Figure 7B**). Supporting phenotype specificity, the molecular phenotypes induced by *hoxc9* mutation were not observed in mutant tadpoles or frogs for either *foxp1* or *en1* CRISPR mutant animals that were generated concurrently.^131^

Interneuron projection patterns and connectivity were not explicitly measured in this study, due to the technical difficulty in frogs of tracing neuronal connectivity patterns in a cell- and class-specific manner, despite recent advances in population-level viral labeling.^179^ Instead, for computational modeling, coarse projection patterns in frogs were assumed to be similar to those of mice. The validity of this assumption is supported by overall similarities of ipsi-/contralateral projections of analogous cardinal types in zebrafish, tadpoles and mice.^14,180,181^ The assumption of similar connectivity allows us to specifically isolate the effects of changing cellular number and distribution. Species-specific connectivity is however probable and to account for this, connectivity was sampled probabilistically, thus providing intra-class variability and limiting top-down decision biases (*see* **STAR Methods**). We however cannot rule out the existence of frog- or mouse-specific projections and/or connectivity that our model fails to reproduce. Future analysis of wildtype and *hoxc9*-mutant circuits across species may reveal conserved and divergent interneuron projection patterns, as well as their potential Hox-dependent regulation.^15,20,155^

## RESOURCE AVAILABILITY

### Lead contact

Further information and requests for resources and reagents should be directed to and will be fulfilled by the lead contact, Lora B. Sweeney.

### Materials availability

All materials will be made available upon request to the lead contact.

### Data and code availability

- Code used for spatial comparison and neurogenesis prediction analyses is available at GitHub (https://github.com/stpapadop/Papadopoulos2026etal). Code used for model simulations is available at GitHub (https://github.com/EarendilTheMariner/Papadopoulos2026etal_ModelCodebase).
- Any additional information required to reanalyze the data reported in this paper is available upon request to the lead contact.

## ACKNOWLEDGEMENTS

We thank Anna Kicheva, Thomas Minchington, and members of Sweeney Lab for feedback on the manuscript. We also thank Marito Hayashi, Sophie Gobeil, Giulia Silvestrelli for assistance with selected mouse experiments; Filip Knop, Matthijs Smits and the Aquatics and Imaging and Optics Facilities (ISTA) for technical support. In addition, we thank our funding sources for providing resources to carry out this work: Horizon Europe ERC Starting Grant 101041551 (S.P., F.A.T., L.B.S.); Special Research Program (SFB) of the Austrian Science Fund (FWF) F7814-B (S.P., L.B.S.); Austrian Academy of Sciences (ÖAW) DOC Fellowship 27229 (S.P.); Novonordisk foundation NNF23OC0082192 (A.W.); Natural Sciences and Engineering Research Council of Canada Undergraduate Student Research Award (K.E.); ISTfellow postdoctoral fellowship (C.J.); Wellcome Trust Early Career Award 225674/Z/22/Z (R.R.); Canadian Health Institute of Research PJT-173547 and Natural Sciences and Engineering Research Council of Canada RGPIN 04880 (Y.Z.); ERC Consolidator Grant 819603 SYNAPSEEK (T.P.V.); ISTA Interdisciplinary Project Committee (IPC) Grant 61 (T.P.V., L.B.S); and French Centre National des Études Spatiales (CNES) and Centre National de la Recherche Scientifique (CNRS) (D.L.R.).

## AUTHOR CONTRIBUTIONS

L.B.S. led the project. L.B.S. and S.P. devised and coordinated the project. S.P., C.J., F.A.T. and L.B.S. performed genetic and molecular experiments. D.L.R. and S.P. performed electrophysiological experiments. S.P., R.R., K.E. and Y.Z. performed mouse experiments. A.W. in coordination with S.P., T.P.V. and L.B.S. performed model simulations and computational analysis. S.P., A.W., D.L.R. and L.B.S. wrote the manuscript. R.R., T.P.V. and Y.Z. edited the manuscript and provided critical feedback.

## DECLARATION OF INTERESTS

The authors declare no competing interests.

## DECLARATION OF GENERATIVE AI AND AI-ASSISTED TECHNOLOGIES IN THE WRITING PROCESS

During the preparation of this work, the authors used ChatGPT 5.5 and DeepSeek V3.2 to brainstorm alternative, more concise language, and coding solutions. After using these tools/services, the authors reviewed and edited the content as needed and take full responsibility for the content of the published article.

## STAR METHODS

### KEY RESOURCES TABLE

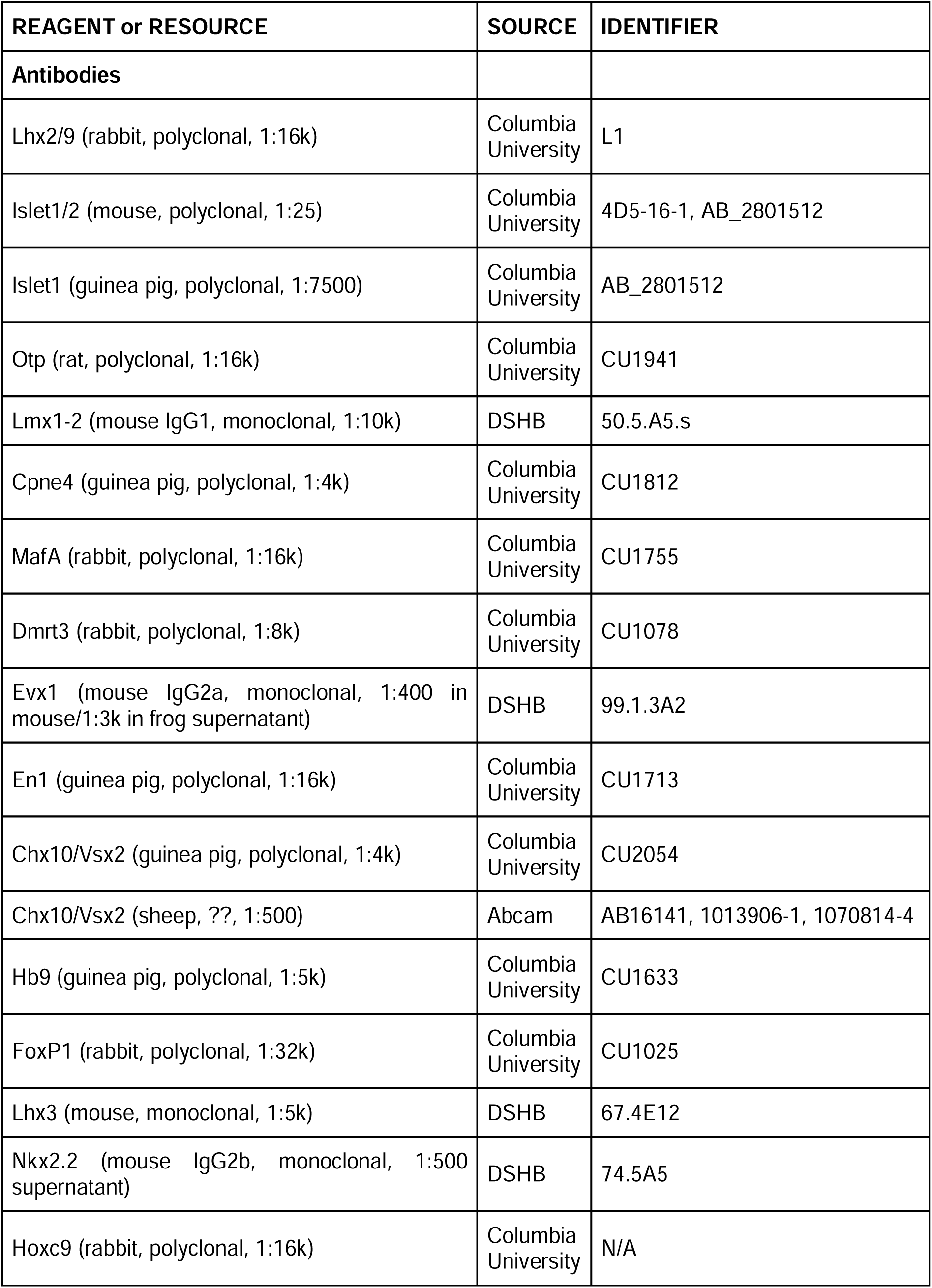
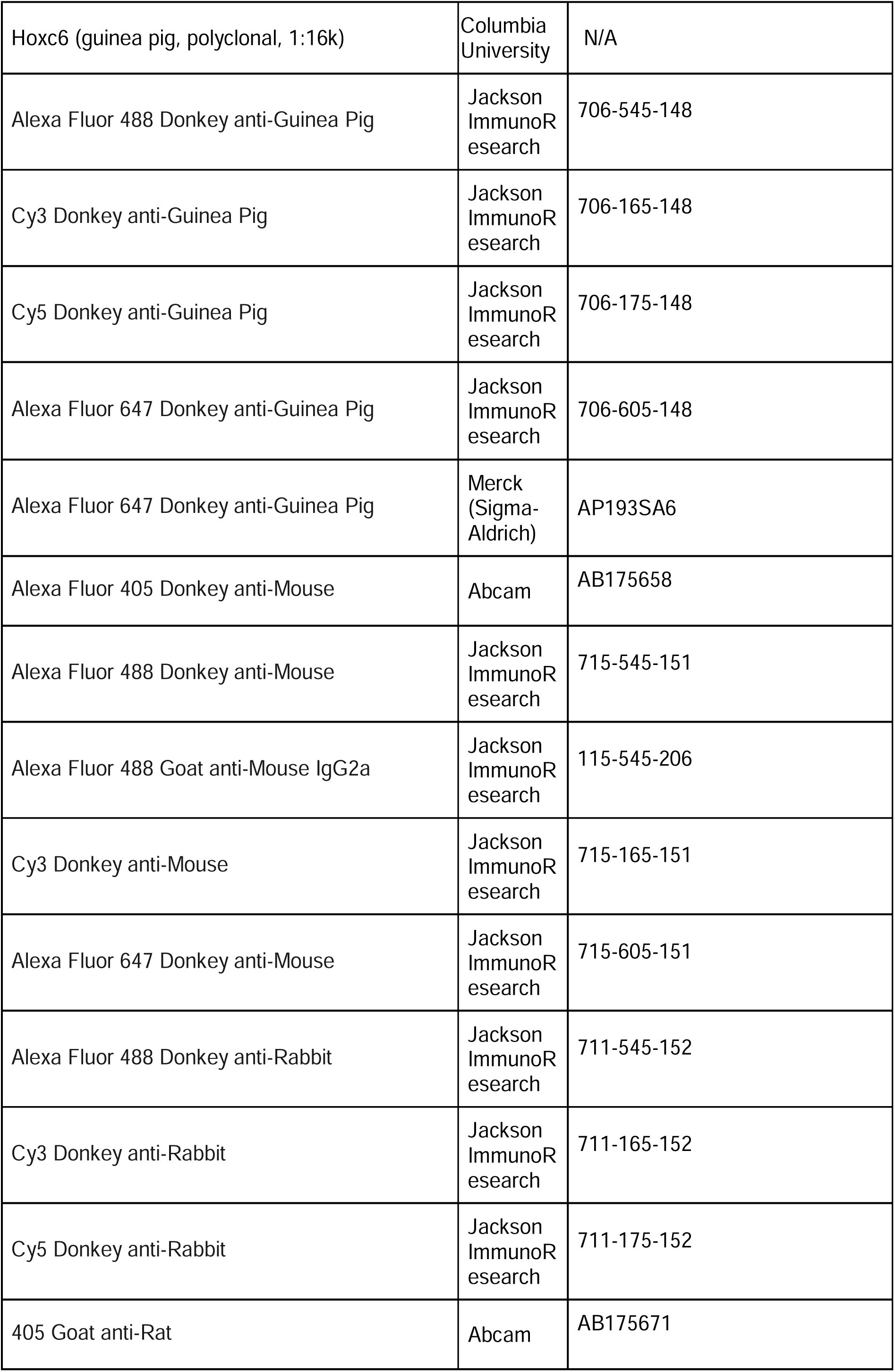
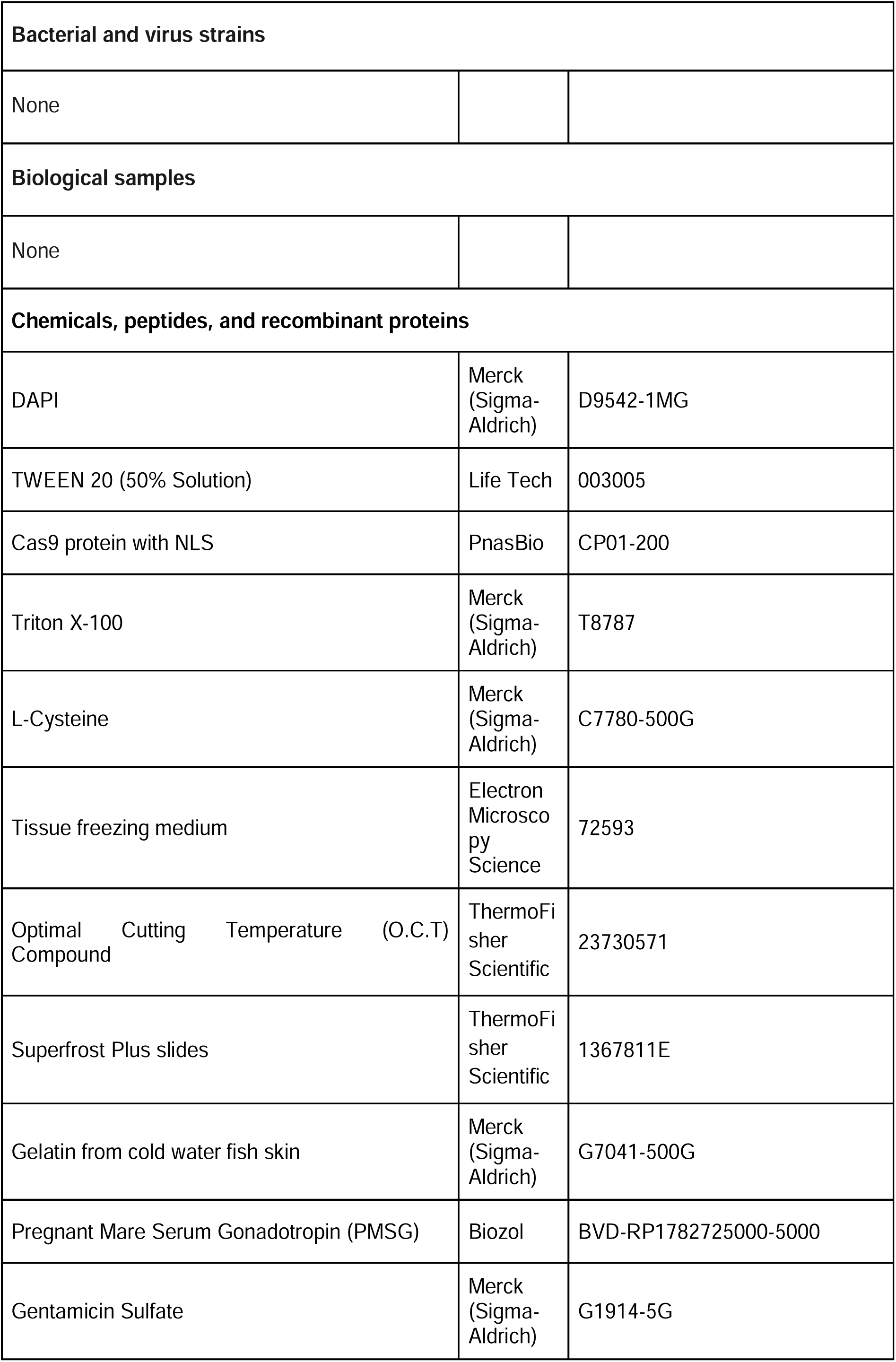
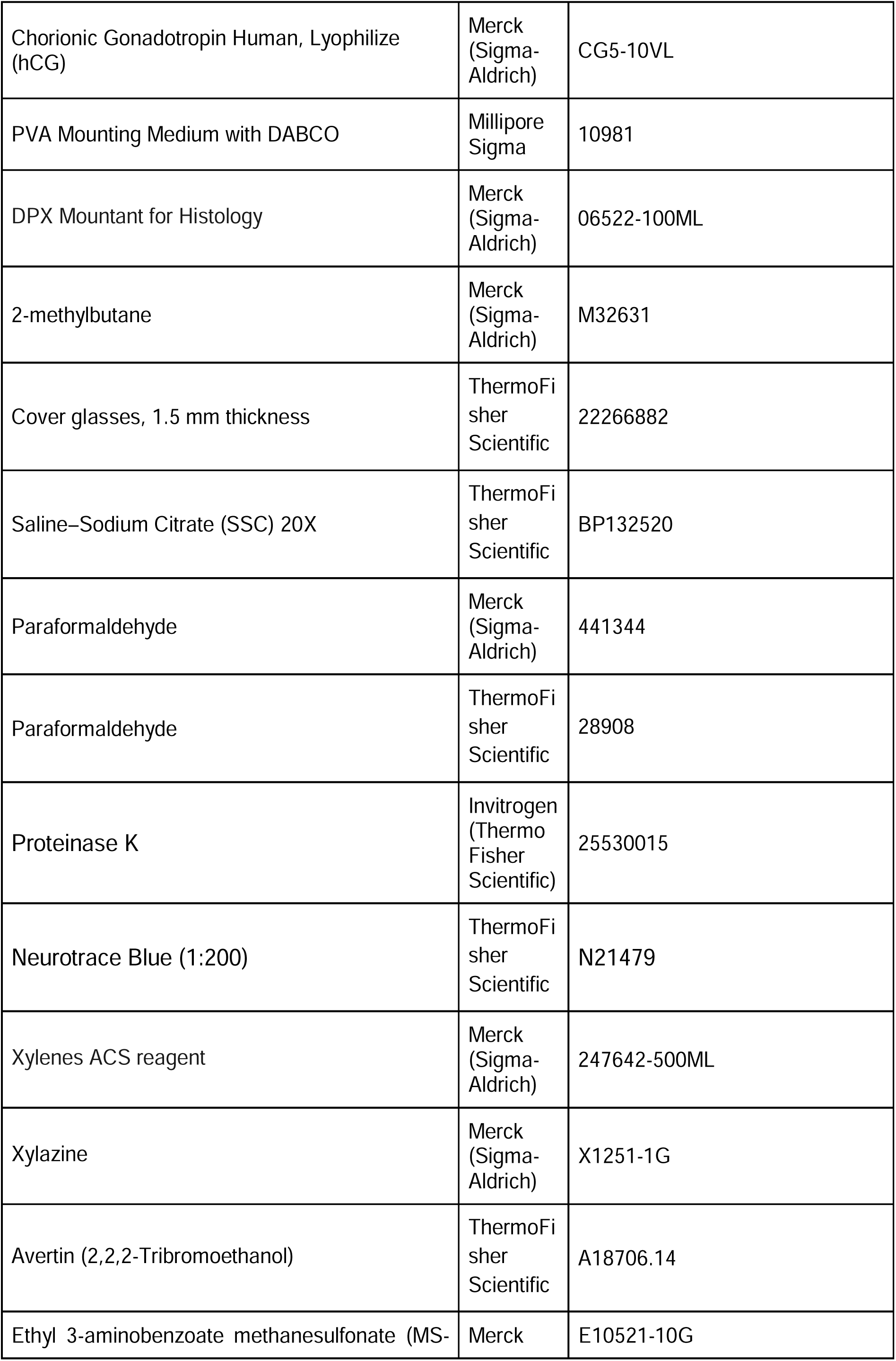
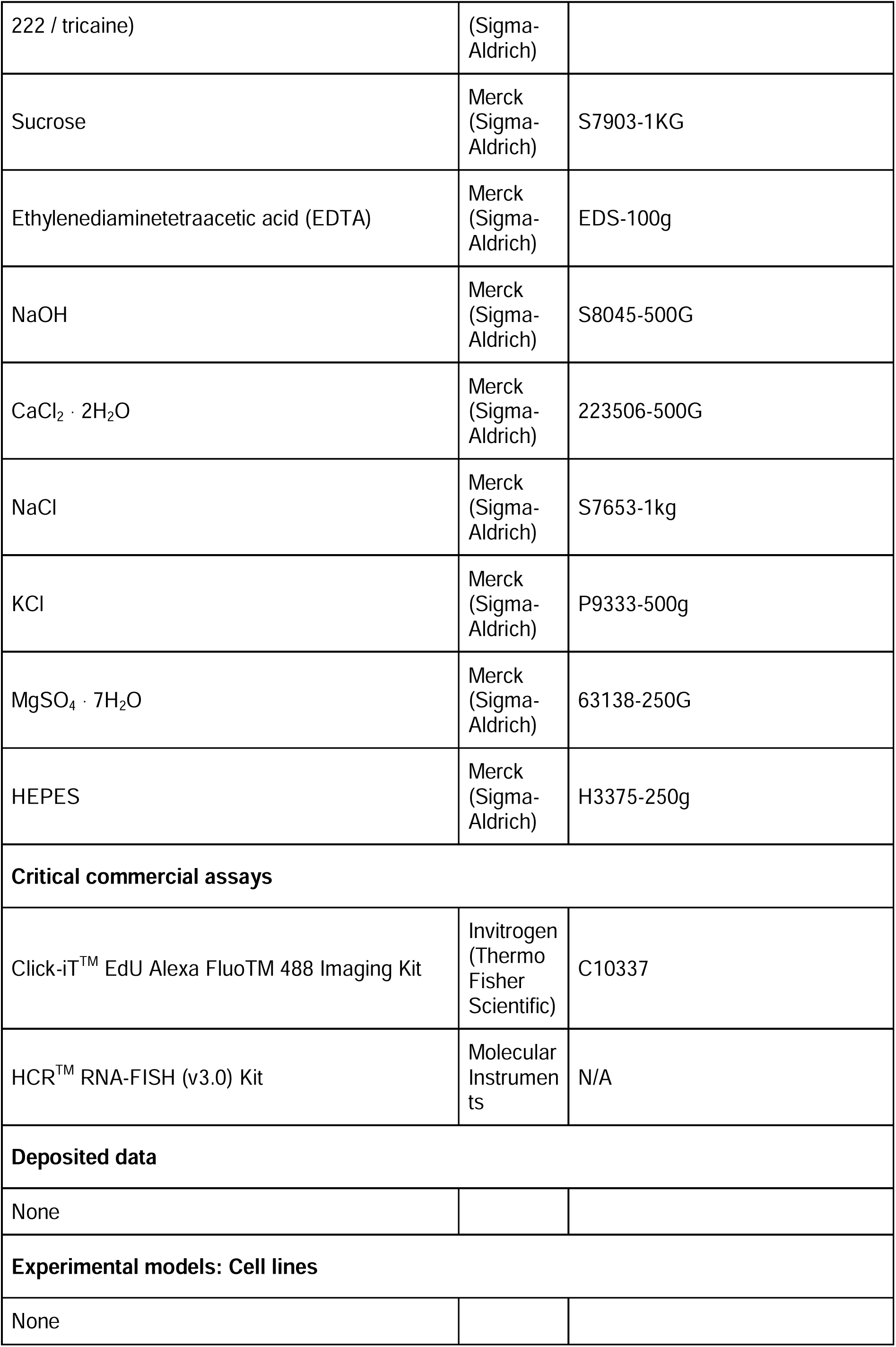
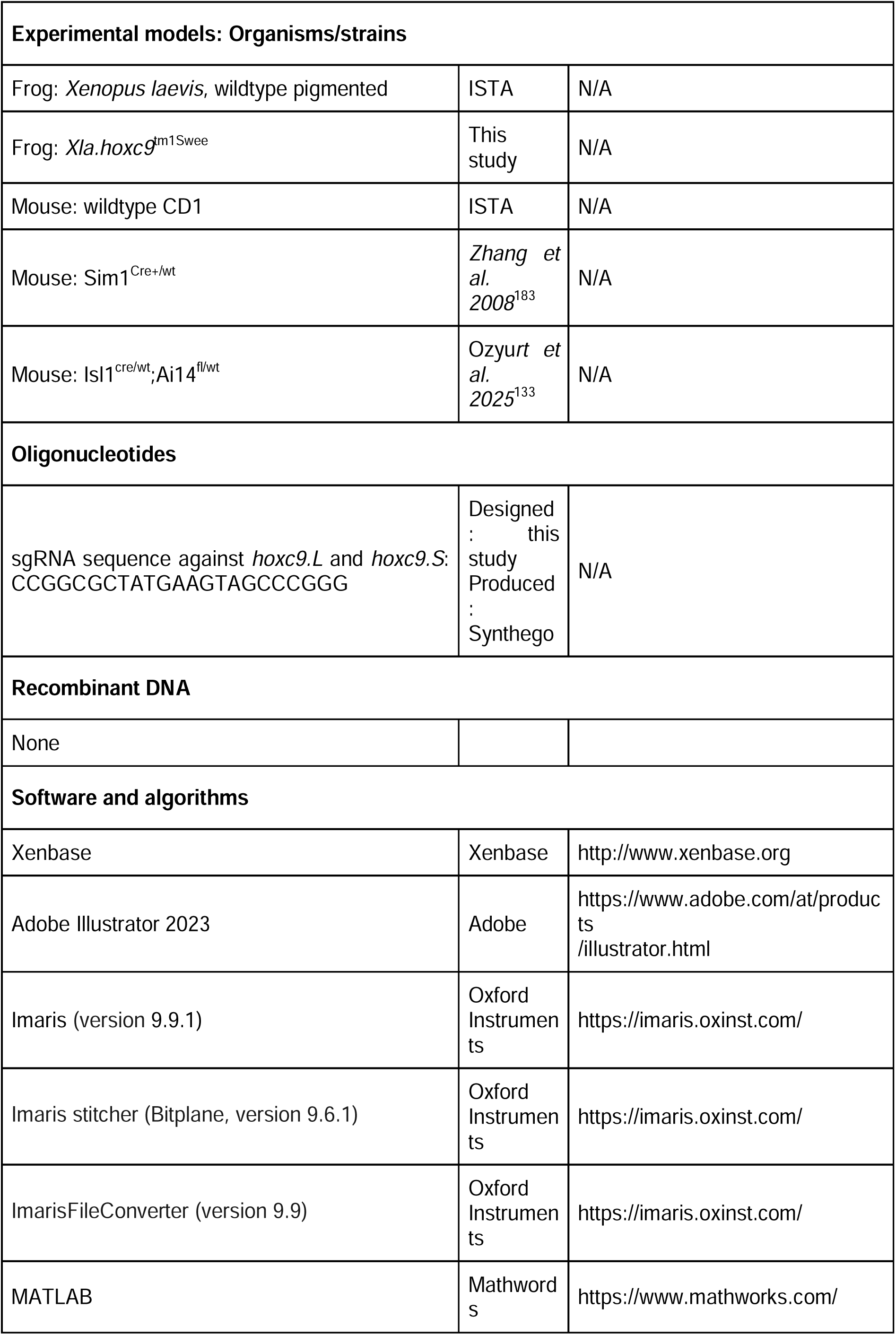
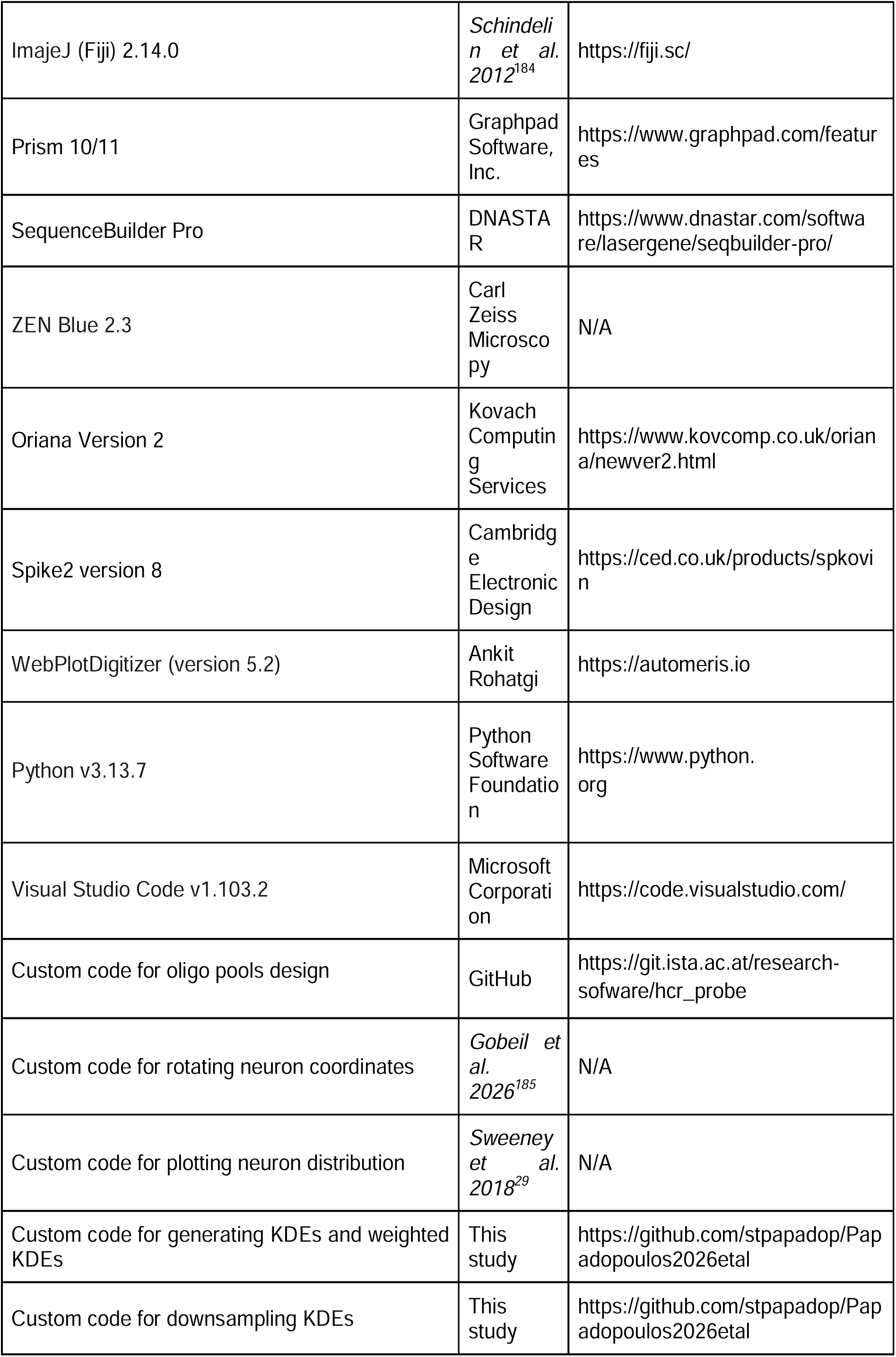
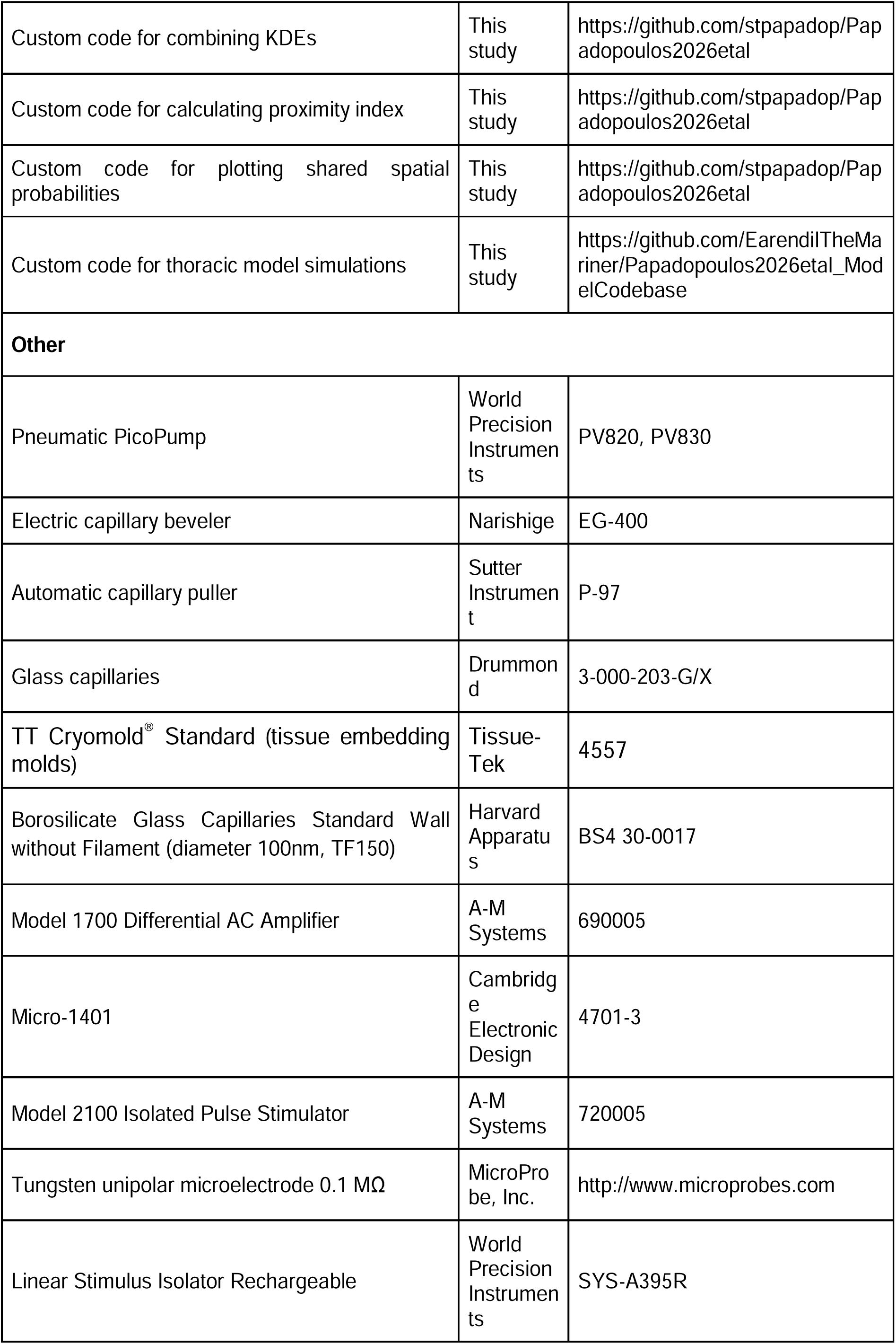

### EXPERIMENTAL MODEL AND STUDY PARTICIPANT DETAILS

#### Frog

All experimental procedures were approved by the Austrian Federal Ministry of Science and Research and carried out in compliance with the Austrian and European Union animal law (permit numbers: 2020-0.550.806, 2020-0.762.370, 2022-0.137.228, 2023-0.591.050 and 2024-0.019.606).

Wild-type pigmented *Xenopus laevis* animals were bred and raised at the Institute of Science and Technology Austria (ISTA) in specialized frog facilities under controlled conditions: 18–22 °C on a 12-hour light/12-hour dark cycle. Tadpoles and juvenile frogs of both sexes were used.

#### Mouse

Experimental procedures took place in three separate locations: ISTA, Dalhousie University, and University College London (UCL), under the approval of the Austrian Federal Ministry of Science and Research and in compliance with the Austrian and European Union animal law (license numbers: 2024-0.191.328 and 2025-0.146.987) or in accordance with the Canadian Council on Animal Care and under the approval of the University Committee on Laboratory Animals at Dalhousie University or in accordance with the Animals (Scientific Procedures) Act UK (1986) and under the approval of the UCL AWERB committee (license number: PP2688499), respectively.

Mice were kept in animal facilities under controlled conditions (21 ± 1 °C; 40–55% humidity) on a 12-hour light/12-hour dark cycle, following protocols approved by the relevant institutional animal-care and ethics committees. Wild-type CD1 mice of both sexes were used, as well as Sim1^cre/wt^ and Isl1^cre/wt^;Ai14^fl/wt^ mice to study V3 and dI3 neurons, respectively. Generation of these transgenic lines was described previously in *Zhang et al. 2008* and Ozyu*rt et al. 2025*. ^133,183^

### METHOD DETAILS

#### Xenopus laevis husbandry

*Xenopus laevis* adult animals, male and female, were purchased from the National *Xenopus* Resource – NXR (https://www.xenopusnxr.org/nxr/) and the European *Xenopus* Resource Centre – EXRC (https://xenopusresource.org). Females were induced to lay eggs by injection of 100 units Pregnant Mare Serum Gonadotropin (PMSG) and 1000 units human chorionic gonadotropin (hCG) in the dorsal lymph sac, spaced one to two weeks apart. Egg laying took place in 1x egg laying solution (1:10 dilution in MilliQ water of 10x stock: 1.2 M NaCl; 10 mM KCl; 24 mM NaHCO_3_; 8 mM MgSO_4_·7H_2_O; 300 mM Tris; 4 mM CaCl_2_·2H_2_O; 3 mM Ca(NO_3_)_2_·4H_2_O, pH adjusted to 7.4-7.5 with HCl). The eggs were then fertilized *in vitro* using freshly dissected *Xenopus* testes. Embryos were raised in 0.1x MMR^+^ on a 16°C cold plate until stage NF44, when they were moved to the filtered water-system of the frog facility. Embryonic and tadpole staging was based on Nieuwkoop and Faber’s Normal table of *Xenopus laevis*.^39^

#### CRISPR experiments

Based on the *Xenopus laevis* v10.1 genome, a single guide RNA (sgRNA) against the conserved region of the Hoxc9 locus in both short and long chromosomes was designed using ChopChop.^186^ The protospacer adjacent motif (PAM site) was chosen such that it followed a DNA sequence producing a highly conserved amino acid sequence. The *hoxc9*-sgRNA sequence can be found in the **Key Resources Table**. The sgRNA was synthesized at Synthego (https://www.synthego.com/).

To generate *hoxc9* CRISPR loss-of-function mutants injection mix was produced by mixing 5 ng of sgRNA with 3 ng of purified Cas9 protein diluted in DNA-free water at room temperature (final concentration of 10μg/μl for gRNA and 0.5mg/μl for Cas9). Fertilized eggs were collected (as described above) and dejellied using 0.1x Marc’s Modified Ringer’s solution with added 1.3mM of EDTA and 20mM of CaCl2(H_2_O)_2_ (MMR^+^) solution containing 3% cysteine (pH 8.2). Dejellied embryos were washed twice in deionized water and twice in 0.1x MMR^+^, before being transferred to RNA injection buffer (1% Ficoll, 0.5x MMR^+^) on a 16°C cold plate. One-cell stage embryos, 30min post fertilization, or two-cell stage embryos, 2h post fertilization, were injected with injection mix bilaterally (injected volume ∼16nl) or unilaterally (injected volume ∼8nl) to produce whole- and half-body CRISPR mutants, respectively. Post injection embryos were moved to 0.1x MMR^+^ with 1mg/ml gentamicin on a 16°C cold plate until stage NF44. Then they were transferred to 0.1x MMR^+^ at room temperature and cared for as wild type animals.

#### Immunohistochemistry and EdU visualization – frog

Stage NF55 animals were sacrificed in 0.1% (w/v) ethyl 3-aminobenzoate methanesulfonate (tricaine) solution (MS-222, 1L of 0.1x MMR^+^, 1g of Tricaine, 0.15g of NaHCO_3_). Spinal cord and surrounding tissue were coarsely dissected and fixed in 4% paraformaldehyde in 1x PBS for 90 min on ice. The tissue was then incubated overnight at 4°C in cryoprotectant solution: 15% (w/v) sucrose, 8% (w/v) fish jelly [50% (w/v) fish skin gelatine in MilliQ H_2_O] in 1x PBS. Afterwards, the tissue was embedded using Tissue Freezing Medium (Electron Microscopy Sciences EMS Catalog #72592) in vinyl molds (TT Cryomold® Standard #4557), and was frozen by immersion in 2-methylbutane at approximately −40°C, before storing at - 70°C.

Frozen tissue-blocks were cryosectioned at 16μm thickness using an OTF6000 (Bright Instrument) or a Cryostat NX70 (Erpedia) cryostat, and the sections were collected on glass microscope slides, which were then stored at −70°C. Before staining the frozen slides were dried for 30 min at room temperature followed by 4-5 h at 4°C, and then rehydrated by washing twice with 1x PBS with 0.2% Triton 100-X (PBST) before incubating in 0.2% PBST for 5-10 minutes. After rehydration, the slides were incubated overnight at 4°C in a dark humidified chamber with a primary antibody mix in 0.2% PBST. To remove unbound antibodies the slides were washed twice with PBST, and then incubated in 0.2% PBST for 5-10min, before a 30-60 min incubation with the secondary antibody mix, at room temperature in a dark humidified chamber. To remove unbound antibodies the slides were washed twice with PBST. If the animals whose tissue was being processed have not undergone EdU injections, the slides were dehydrated by consecutive washes with ddH2O, 25%, 50%, 75%, and twice with 100% EtOH, followed by two washes with Xylenes. Finally, a coverslip was mounted with ∼500 μL of DPX on the dehydrated slides After drying in a dark, ventilated place at room temperature, slides were stored at 4°C. Otherwise, to visualize EdU, the slides were first incubated for 5-10 min in PBST, and then in the EdU reaction solution (864 μL MilliQ H_2_O, 86 μL 10x Click-iT reaction buffer, 40 μL CuSO_4_, 2.5 μL Alexa fluor azide, 10 μL reaction buffer additive – Thermo Fisher) for 60-90 min, at room temperature in a dark humidified chamber. After which, the slides were dehydrated and a coverslip was mounted as described previously.

#### Tissue collection and immunohistochemistry – mouse

Adult mice were anesthetized via intraperitoneal injections of a ketamine and xylazine (60-80 and 10-12 mg/kg, respectively), for pregnant females Avertin (Tribromoethanol, 240 mg/kg) was used. They were then transcardially perfused with ice-cold PBS followed by 4% paraformaldehyde (PFA, Electron Microscopy Sciences or ThermoFisher 28908) in PBS. Adult spinal cords were dissected and post-fixed in 4% PFA for 2-4 h on ice and cryoprotected in 30% sucrose in PBS for a minimum of 24 h at 4°C. E18.5 embryos were anesthetized by inhalation of isoflurane and transcardially perfused with ice-cold 4% PFA, E13.5 embryos were not individually perfused. All embryos were post-fixed in 4% PFA for 90 min on ice and cryoprotected in 30% sucrose in PB mix (1:1 0.2M phosphate buffer:MilliQ H_2_O) overnight. E18.5 spinal cords were then dissected. Cryoprotected spinal cords and embryos were then embedded in PolyFreeze (Polysciences, 25113-1) or optimal cutting temperature compound (OCT, Tissue-Tek 4583 or Fisher Healthcare) and frozen. Sectioning was performed with a cryostat (Leica CM3050 S, or CM1950, or OTF6000 – Bright Instrument, or Cryostat NX70 – Erpedia) at 16-30 μm and mounted onto Superfrost Plus Microscope Slides (Fisherbrand).

Spinal cord sections were put at 4°C following cryosecting until dry. They were then rehydrated by washing with PBS or PBS with 0.1-0.2% Triton 100-X (PBST). Depending on antibody penetration adult sections were at this point blocked for 30 min in PBS 2x (0.2 M), 0.2-0.3% Triton 100-X, 10 % donkey normal serum. Sections were then incubated with primary antibody solution in 0.1-0.2% PBST and 10% heat-inactivated horse serum (Invitrogen) or blocking solution. Embryonic tissue was not blocked. Primary antibody incubation lasted 12-48 h depending on antibody efficacy and section thickness, at 4°C. Slides were then washed three times with PBS for 15 min, and incubated with secondary antibody solution in PBS for 1-12 h at room temperature (embryonic tissue) or at 4°C (adult tissue). After washing three times with PBS for 15 min, a coverslip was mounted with fluorescent mounting medium (Dako), PVA/DABCO antifade medium (10981 or 81381, Sigma), or custom Mowiol-DABCO (25% w/v glycerol, 10% 960 w/v Mowiol, 2.5% w/v DABCO in 0.2 M Tris-HCl pH 8.5), and the slides were stored at 4°C.

#### Image acquisition and processing

Images were obtained using the following confocal microscopes: Zeiss LSM510 upright, LSM800 and LSM900 inverted; Leica Sp8 upright and inverted; Nikon CSU-W1 dual-disk spinning disk. 10x and 20x air objectives were used in all cases. Whenever tiles were collected, they were stitched either using Zen Blue (ZEN Blue 2.3 software), or Imaris stitcher (Bitplane, version 9.6.1). Images are presented as z-projections. Images were analyzed using ImajeJ (version 2.14.0, 1.54k) or Imaris (version 9.9.1) after microscope files were first transformed into Imaris file format using ImarisFileConverter (version 9.9). Marker-positive cells were quantified manually using the Cell Counter plugin of ImageJ, or using the automated “Spots” function of Imaris before manual validation.

Neuron cardinal classes and subpopulations were defined based on established combinatorial marker expression.^41,43^ For all datasets at least three hemisections per level per animal were analyzed (details for each case are given in the figure legends), rostrocaudally at approximately the middle region of each of the brachial, thoracic and lumbar levels. For Sim1^cre/wt^ adult mice, only the ventral population was analyzed (V3v) for a direct comparison with the ventral Nkx2.2 expressing V3 population in frog and embryonic mouse. Likewise, in Isl1^cre/wt^;Ai14^fl/wt^ p11 and adult mice only dI3 neurons were counted – excluding motor neurons based on position, morphology and in some datasets ChAT staining.

Marker colocalization was scored using Imaris either manually or implementing a custom MATLAB script (previously described in *Sweeney et al. 2018*) followed by manual validation.^29^ To generate kernel density estimate maps (KDEs) on a common fixed rectilinear grid, we first extracted the XY coordinates of the marked cells from Imaris using another custom MATLAB script (also previously described in *Sweeney et al. 2018*).^29^ This MATLAB script then allowed us to normalize cell coordinates to a standard spinal hemisection using three reference points (central canal, most ventral point of the white matter, and most lateral point of the white matter). For populations extending to the dorsal spinal cord a fourth reference point was also used (most dorsal point of the white matter), this adapted MATLAB/Python pipeline was previously described in *Gobeil et al. 2026*.^185^

To generate the proximity index of spatial distributions we used a Python script that calculated the difference between every KDE grid cell of one distribution and the respective KDE grid cell of a second distribution. This comparison was then summed up into a single value that was added to the normalized distance between the highest density point of each KDE, thus producing a single value representing how similar two distributions were. KDE grid comparisons were also used to make the neurogenesis predictions. Here another Python script was used, which did two things: first, normalized the KDEs of EdU-positive cells for each time point between NF48 and 54 for each brachial, thoracic, and lumbar level, and second, by using these normalized EdU KDEs and the distribution KDEs of cells expressing different neuron class markers calculated the combined probability of every KDE grid cell for both EdU and neuron marker per timepoint and rostrocaudal level. These Python scripts can be found: https://github.com/stpapadop/Papadopoulos2026etal. We then calculated in which of the seven (NF48-54) combined probability KDE grids of each neuron population the top 90% of the total values belonged, and calculated the percentage for each stage. With either the experimentally calculated neurogenesis fraction for individual populations, or the dorsal or ventral average for the populations that were not directly assessed (**Figure S10H, S11B**), we calculated how many neurons of each population present at NF55 (**Figure 1D-M**) would have been captured in a theoretical EdU series between NF48 and NF54. Using this extrapolated neuron number in combination with the predicted percentage of neurogenesis per stage from our KDE analysis we were able to predict the number of neurons per class by level that would have been born at each stage of limb circuit generation (**Figure S11D, S11F-G, S11I-J**).

#### Neurogenesis characterization in *Xenopus* tadpoles

Stage NF48-54 animals were anesthetized for a few minutes (≤20min) using a 0.01% tricaine methanesulfonate solution (MS-222, 1L of 0.1xMMR^+^, 0.1g of Tricaine, 0.15g of NaHCO_3_), and then placed ventral side up on a gauze kept moist with anesthetic solution. Using a beveled glass needle, 50μg of 5-Ethynyl Uridine (EdU) per 1g of body weight diluted in MilliQ water was introduced via intracoelomic injection. The injections for all stages took place in three consecutive days, with two injections per day, spaced 6 h and 18 h apart for the same day or two consecutive days, respectively, based on a ∼2.5-3.5 h uridine analog availability, a ∼5-7 h S-phase, and a ∼35-42 h cell-cycle length reported in *Thuret et al. 2015* for the posterior hindbrain/anterior spinal cord.^76^ During the injection period the animals were kept at 23°C to counter possible deceleration of development due to stress, water was changed before the first injection and food was added after the last injection, daily. On the fourth day they were moved to a filtered water-system at 21°C, and raised to three days after stage NF55, when they were sacrificed and processed as described in the ***Immunohistochemistry and EdU visualization – frog*** and ***Image acquisition and processing*** sections.

### QUANTIFICATION AND STATISTICAL ANALYSIS

For both frog and mouse brachial, thoracic, and lumbar spinal rostrocaudal region boundaries were defined based on level-specific motorneuron columns.^22^ In post-natal mouse rostrocaudal were also identified based on ventral roots. For limb regions, in NF55 and NF66 *Xenopus* animals all neurons (motor and interneurons) were scored at the level where the LMC is at its largest, approximately two thirds caudally within the brachial and lumbar segments. In NF47 *Xenopus* tadpoles Raldh2 expression that marks limb-projecting motor neurons was used to reveal the brachial and lumbar regions, and neurons were scored approximately in the middle of these regions. Across the three stages all thoracic neurons were scored at the approximate midpoint between the caudal and rostral boundaries of the brachial and lumbar regions, respectively. Similarly, E13.5, E18 and post-natal mouse data were collected at approximately the middle of the respective brachial, thoracic, and lumbar regions. For post-natal V1, V2a, and dI6 neurons we made use of already published data adapted from *Sweeney et al. 2018*, *Hayashi et al. 2018*, *and Perry et al. 2019*.^29,32,68^ V1 and V2a were kindly supplied by the authors. dI6 data were extracted using the WebPlotDigitizer software (https://automeris.io, version 5.2). Adult (p82-149) data for dI3 for the lumbar spinal region are reused with the authors’ permission from *Ozyurt et al. 2025* and *Ronzano et al. 2026*; and additionally, from *Ronzano et al. 2026* for the thoracic region.^24,133^ For each rostrocaudal level a minimum of three hemisections per animal for two animals was quantified across species and developmental stages, with the actual N reported at the legend of each relevant figure. Images were quantified as three dimensional z-stacks with a z size of 16 μm, if the z size happened to exceed this the data were post hoc normalized to 16 μm. The numbers of V2b interneurons used in the frog model were defined as ⅓ of the V1 numbers at either the brachial or the thoracic region, based on the the V2b/V1 ratio being ⅓ in stage NF54 *Xenopus* single cell RNA sequencing data, and spatial distribution was extracted from spatial transcriptomics data of these regions for the same developmental stage.^43^ The numbers and spatial distributions of V0d and V2b used in the mouse spinal model were based on E12.5 transgenic lines of Dbx1^Lac/wt^ and Gata3^Cre/wt^, respectively.^33^ For V0d, we considered Dbx1^Lac/wt^ marked neurons that were not co-labeled with Evx1 antibody, as Dbx1^Lac/wt^ was shown to mark the whole V0 class and only that. Numbers and coordinates were extracted from the published images using Imaris 9.9.1. In the mouse wildtype thoracic model the number and spatial distribution of ‘MMC’ refer to the Isl1/2^+^FoxP1^-^ (MMC/HMC) population (**Figure S5C**).

Using Prism 10/11, cell counts underwent statistical testing as indicated in the figure legends in each case. For rostrocaudal comparisons one-way ANOVA and Tukey’s multiple comparison tests were performed. While, for neurogenesis timeline data two-way ANOVA and uncorrected Fisher’s LSD tests for multiple comparisons were performed. In all graphs mean ± SEM is shown, and **** P<0.0001, *** P<0.001, ** P<0.01, * P<0.05, and ns P>0.05. Illustrations were created using Adobe Illustrator 2023.

#### Hybridization Chain Reaction (HCR) on frozen *Xenopus* sections

Fluorescent in situ mRNA Hybridization using Hybridization chain Reaction was conducted as described in the 2026 edition of *Xenopus: Methods and Protocols* Chapter 11 – Fluorescent In Situ mRNA Hybridization (FISH) Using Hybridization Chain Reaction (HCR) in *Xenopus* Cryosections (*Vijatovic et al. 2026*^187^) Oligo pools were designed using a custom Python script available (https://git.ista.ac.at/research-sofware/hcr_probe), based on the *Xenopus laevis* genome (v10.1).^186^ Oligo pools were commercially synthesized by Integrated DNA Technologies (eu.idtdna.com/) and the oligo pool sequences against the mRNA targets used can be found in **Table S1**.

HCR images were acquired using a Leica Sp8 upright microscope with a 63x glycerol immersion objective (1.30 refractive index), and processed using ImajeJ (version, 1.54k). Progenitor domain(s) size was defined as the length of the polygonal chain that traced the border of the central canal between the ventral and dorsal edge of the relevant mRNA expression (**Figure S9G**).

#### Galvanic vestibular stimulation (GVS) and electrophysiological recordings from *in vitro* brainstem/spinal semi-intact preparations

Animal dissection for *in vitro* preparations and nerve recording procedures were conducted as previously described.^37,94^ Under deep anesthesia in buffered 0.05% MS-222 and after viscera and forebrain removal, the brainstem-spinal cord together with the thoracic ventral roots (Vr) and the motor nerves innervating ankle flexor (Flex; *tibialis anterior*) and extensor (Ext; *plantaris longus*) on both sides were dissected out from juvenile *Xenopus* frogs. *In vitro* experiments were performed in carbogen (95% O2/5% CO2)-bubbled Ringer’s saline (93.5 mM NaCl, 3 mM KCl, 30 mM NaHCO_3_, 0.5 mM NaH_2_PO_4_, 2.6 mM CaCl_2_, 1 mM MgCl_2_, and 11 mM glucose, pH 7.4). The two otic capsules were kept intact and attached to the brainstem with the VIII^th^ nerve. Extracellular motor activity was recorded bilaterally from the three thoracic Vr with borosilicate glass suction electrodes (tip diameter, 100 nm; TF150; Harvard Apparatus) filled with Ringer’s solution. Extracellular activity from Ext/Flex motor nerves was recorded using stainless *en passant* wire electrodes. Electrophysiological signals were directed to a Model-1700 amplifier (AM-Systems) and digitized at 10kHz through a Micro-1401 interface (CED: Cambridge Electronic Design). All electrophysiological signals were acquired, stored, and analyzed using Spike2 software (CED).

Spontaneous or trigeminal nerve electrical stimulation-evoked locomotion was recorded from thoracic Vr and Ext/Flex motor nerves. Trigeminal nerve stimulation was delivered as single or repeated 100-µs pulses (30-80 V) using a Model-2100 stimulator (AM-Systems), through unipolar 0.1-MΩ tungsten microelectrode (MicroProbe) placed in contact with the nerve on either side of the brainstem.

Vr and Ext/Flex motor nerve reflex bursts were recorded simultaneously in response to galvanic vestibular stimulation (GVS). In order to activate the vestibular end organs that generate the most consistent spinal responses in *Xenopus*,^37,96^ GVS was delivered using two coated iron electrodes placed in contact with the otic capsule on both sides vertically to the horizontal canal sensory cells. GVS consisted of sequences of sinusoidal current (10 cycles; 0.1-1 Hz; ±1-5 mA) applied simultaneously but in phase opposition through the left and right otic capsules, close to the canal cupula. 24.1% of GVS sequences in wildtypes and 38.9% in mutants evoked fictive swimming episodes, which were not investigated further because they dramatically interfered with vestibular-evoked spinal reflexes. The 0.25 and 0.5 Hz stimulation frequencies proved optimal to generate reproducible spinal reflex responses in wild-type animals, and all data reported here were obtained using such GVS parameters (**Figure 7E-H**).

#### Statistical analysis of electrophysiological data

GVS-evoked reflex responses were analyzed with circular statistics using Oriana 2 software (Kovacs). Single phases of responses were measured for each cycle as the time to reach maximal responses of a given ventral root or nerve referred to the cycle duration, leading to the calculation of the mean phase response for a given motor root/nerve as the mean vector angle for the stimulation sequence. Only significant mean phases were considered further, i.e. when mean vector length was larger than 0.5 and the Rayleigh distribution uniformity test was significant. Thereafter, significant left and right mean vectors were compared for each thoracic segmental Vr and limb motor nerves. Left-right analysis of limb muscle activity was used to separate GVS cycle-related reflex responses (characterized by left/right phase-locked alternation) from sensory-triggered locomotor episodes (characterized by left/right synchrony between agonistic muscles).^37,94^ GVS-evoked thoracic responses were considered for further analysis only in the absence of sensory-evoked locomotor activity. In these conditions, because no significant differences in the response phase were observed between the three thoracic Vr on the same side, all thoracic mean vectors were combined for each side, and the global circular distribution of the relative left/right thoracic coordination was then plotted. A final resulting grand mean vector was then calculated for the whole population of thoracic Vr responses to GVS, the length (r) of which indicated the robustness of the distribution (the greater, the more robust) and the mean angle (ma) illustrated the left/right coordination (0°=synchrony, 180°=alternation; **Figure 7C-D, 7K-L**). The putative difference in GVS ability to evoke thoracic responses was tested with a non-parametric t-test (Mann-Whitney) using Prism 10/11 (Graph Pad).

Locomotor activity was analyzed with circular statistics, and all single motor root/nerve bursts were compared to the left ankle extensor (*plantaris longus*) nerve burst as a reference. As above, all phase relationships for a given root/nerve in an episode were used to calculate a mean phase value representative of the swimming episode. Then, all significant mean phase values were pooled together, and a grand mean vector was calculated for each motor root/nerve and used to illustrate the population mean coordination during swimming (**Figure 7I-J**).

## Notes

### Competing Interest Statement

The authors have declared no competing interest.

