## Supplemental Material for "Spinal circuit regionalization diversifies motor output along the vertebrate body axis"

### 1 Methods

#### Frog and Mouse Segment Models

In our frog models, we populated thoracic and brachialized segment volumes with interneuron types V0v, V1, V2a, V2b, V3v, dI6 and motor neuron types MMC and LMC. Mediolateral ($x$) and dorsoventral ($y$) spatial coordinates and cell type proportions were determined by our data analysis at NF55 at thoracic and brachial levels. Assuming that transversal cell type distributions extend throughout each segment, longitudinal coordinates ($z$) were sampled uniformly for each segment, with a total length of $4125\mu m$. The three ventral roots were evenly spaced longitudinally within each model segment. Partially brachialized thoracic segment volumes were constructed assigning brachial cell counts and transversal positions to a subset of cell types in an otherwise thoracic cell composition.
The mouse thoracic segment was populated with interneuron types V0v, V0d, V1, V2a, V2b, V3v, dI6 and the motor neuron type MMC. Transversal coordinates and cell quantitites were extracted from mouse embryonal stages. Longitudinal coordinates were sampled uniformly within a total length of $8250\mu m$. The thirteen ventral roots were evenly spaced longitudinally within the segment.
For mouse and frog cell type mapping and quantification, see **STAR Methods: QUANTIFICATION AND STATISTICAL ANALYSIS**.

#### Network Connectivity

Synapses between neurons were sampled probabilistically via mouse-like spatial projection biases, as devised in *Komi et al, 2026*,[91] but reformatted here for convenience. To determine synaptic connectivity, we computed the pairwise distances between neurons in each spatial dimension independently:

$$d_{X,ij} = x_i - x_j, \quad d_{Y,ij} = y_i - y_j, \quad d_{Z,ij} = z_i - z_j$$

We then modeled the probability of a synaptic connection between neurons $i$ and $j$ using Gaussian distributions for each dimension:

$$P_X^p(d_{X,ij}) \sim \mathcal{N}(\mu_X^p, \sigma_X^p), \quad P_Y^p(d_{Y,ij}) \sim \mathcal{N}(\mu_Y^p, \sigma_Y^p), \quad P_Z^p(d_{Z,ij}) \sim \mathcal{N}(\mu_Z^p, \sigma_Z^p)$$

where $\mu_X^p$, $\mu_Y^p$, and $\mu_Z^p$ are the means, and $\sigma_X^p$, $\sigma_Y^p$, and $\sigma_Z^p$ are the standard deviations. The overall synaptic probability was computed as:

$$\mathbf{P}_{syn}(i,j) = \left(P_X^p(d_{X,ij}) \cdot P_Y^p(d_{Y,ij}) \cdot P_Z^p(d_{Z,ij})\right)^{1/3}$$

This combines the probabilities from each dimension into a single measure of connection likelihood. We then binarized this probability to create a binary adjacency matrix $\mathbf{A}$:

$$\mathbf{A}(i,j) \sim \text{Binomial}(1, \mathbf{P}_{syn}(i,j))$$

where $\mathbf{A}(i,j) = 1$ indicates a synaptic connection between neurons $i$ and $j$. The hadamard product of $\mathbf{A}$ and the synaptic weight matrix, $\mathbf{G}(i,j) \sim \mathcal{N}(1, 0.1)$ yielded the connectivity matrix:

$$\mathbf{W} = \mathbf{G} \circ \mathbf{A}$$

where excitatory (inhibitory) columns of $\mathbf{W}$ were made strictly non-negative (non-positive). Finally, a detailed excitatory/inhibitory balance was imposed on the connectivity as the values of the rows adding up to zero, i.e., $\mathbf{W}_{i,:}^T \mathbf{1}_n = 0$ for all postsynaptic neurons $i$.

#### Dynamical Model

All networks were analyzed and simulated according to the firing-rate formalism:

$$\tau\dot{\mathbf{x}} = -\mathbf{x} + \mathbf{W}\psi(\mathbf{x}) + \mathbf{I}(t) + \epsilon(t)$$

where $\tau$ is the $n$ dimensional vector of time constants, $\mathbf{x}$ is a vector with the input potentials of all neurons, $\dot{\mathbf{x}}$ its time derivative, and $\mathbf{I}$ is an external input. The noise term is Gaussian with $\epsilon(t) \sim \mathcal{N}(0,\sigma)$. $\psi(\mathbf{x})$ is a sigmoidal transfer function (using the hyperbolic tangent function), mapping the input to the firing rate:

$$\psi(x) = \begin{cases} T \cdot \tanh\left(\frac{g(x-T)}{T}\right) + T, & \text{for } x \leq T \\ f_{\max} \cdot \tanh\left(\frac{g(x-T)}{f_{\max}}\right) + T, & \text{for } x > T. \end{cases}$$

The output firing rate is non-negative and $T$ represents the inflection point of the sigmoid where $\frac{d\psi(x)}{dx} = g$. $T$ and $f_{max}$ together set the maximal asymptotic firing rate. See **Figure S13A** for parameter summary. The networks were simulated according to the Euler rule, $x_{t+dt} = x_t + \dot{x}_t dt$ with time step $dt = 1ms$.

#### Network Input Modes

In this work, we investigated our model response to global and vestibulospinal input. Vestibulospinal targets were sampled from a probability distribution defined over the transversal plane. We constructed this distribution by first binning each experimentally mapped $xy$-coordinate of thoracic MMCs. We then assigned a probability mass to each grid point using a bivariate kernel density estimator (`mvksdensity.m` in MATLAB, bandwidth = 10, kernel = 'normal'), with the MMC $xy$-coordinates as input. The output was a probability mass function, $pmf_{XY}()$, defined over the transversal plane. For each instantiated network, we fed the transversal coordinate of each neuron to the $pmf_{XY}()$, thus assigning a vestibular target probability to each neuron. From these probabilities, we used weighted random sampling of $n = 300$ targets with `datasample.m` in MATLAB. In the global input mode, every cell in the network received input.

#### Analysis: Vestibulospinal Response

We simulated the ventral root output of each network in response to left-right alternating sinusoidal input to the sampled vestibulospinal targets

$$I_{\text{left}}(t) = \sin\left(\frac{t}{T}\right) A + A$$

$$I_{\text{right}}(t) = -\sin\left(\frac{t}{T}\right) A + A$$

with period $T = 500\,\text{ms}$ and amplitude $A = 7.5$ ($A = 2.5$ for mouse). External input to all remaining neurons was zero. The output of the left and right middle thoracic roots were approximated by summing the firing rates of all motor neurons (MNs) in the rostrocaudal vicinity of the root:

$$R_{\text{right}}(t) = \sum_i r_i(t), \qquad R_{\text{left}}(t) = \sum_j r_j(t)$$

where $r_i(t)$ denotes the firing rate of motor neuron $i$ at time $t$, and indices $i$ and $j$ enumerate right and left hemicord motor neurons, respectively. The root traces were subsequently $z$-scored. To examine phase differences between left and right hemicord output, peaks were detected in each root trace. Networks for which the number of detected peaks did not equal the number of cycles in the sinusoidal input (here $n = 8$) were excluded from further analysis. Letting $\mathbf{p}^R$ and $\mathbf{p}^L \in \mathbb{R}^n$ denote the ordered peak-time vectors for the right and left sides, the normalised peak-time difference was computed as:

$$\Delta \mathbf{p} = \frac{\mathbf{p}^L - \mathbf{p}^R}{\bar{\tau}}$$

where $\bar{\tau}$ is the mean interval between consecutive peaks. The mean phase difference was then defined as:

$$\Delta \theta = \frac{2\pi}{n} \sum_{i=1}^{n} |\Delta p_i|$$

scaled to the interval $[0, 2\pi]$. For all thoracic network models introduced (wildtype thoracic, brachialized and partially brachialized for frog, wildtype thoracic for mouse) $n = 100$ networks were instantiated and $\Delta\theta$ was computed for each. Statistical comparison between $\Delta\theta$ distributions were performed with Mann-Whitney U-tests. Bonferroni significance correction at $\alpha = 0.05$ was performed to account for multiple testing.

#### Analysis: Global Tonic Drive

In the global input mode, all cells received a constant positive drive $I = 20$. Ventral root dynamics were investigated at steady state. Phase relationships across and along the length of the spinal cord were analyzed by comparing the z-scored firing rate patterns of the roots. For a given pair of roots $i$ and $j$, their phase relationship was defined as:

$$\Delta \theta_{ij} = \arccos\left( \frac{\mathbf{r}_i \cdot \mathbf{r}_j}{\|\mathbf{r}_i\| \, \|\mathbf{r}_j\|} \right)$$

where $\mathbf{r}_i \in \mathbb{R}^T$ denotes the vector of firing rates of root $i$ sampled at $T = 4000$ discrete steady-state time points, and $\cdot$ and $\| \cdot \|$ denote the Euclidean inner product and norm, respectively.

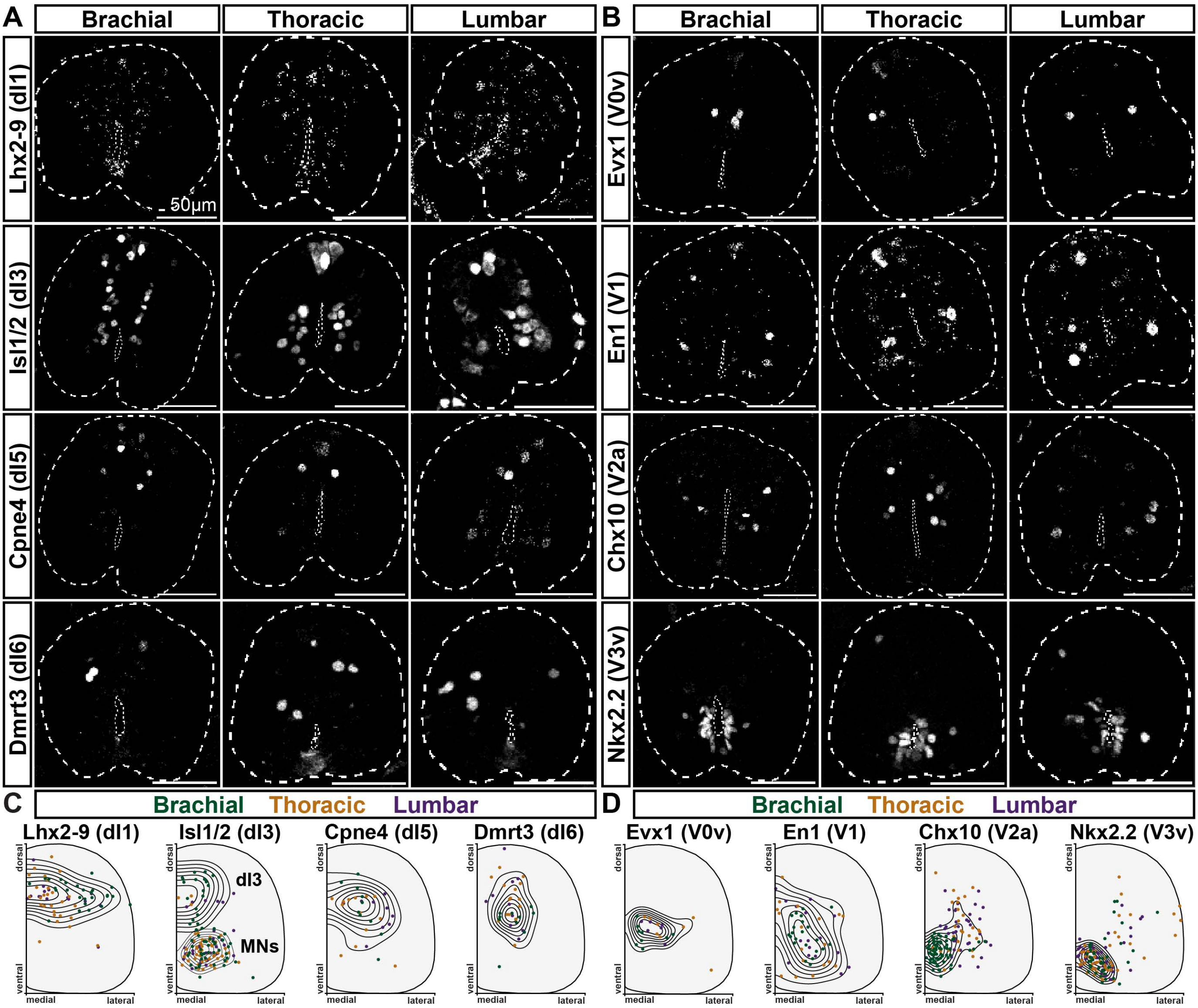
A
Brachial
Thoracic
Lumbar
B
Brachial
Thoracic
Lumbar
Lhx2-9 (dI1)
Isl1/2 (dI3)
Cpne4 (dI5)
Dmrt3 (dI6)
Evx1 (V0v)
En1 (V1)
Chx10 (V2a)
Nkx2.2 (V3v)
50µm
C
Brachial Thoracic Lumbar
D
Brachial Thoracic Lumbar
Lhx2-9 (dI1)
Isl1/2 (dI3)
Cpne4 (dI5)
Dmrt3 (dI6)
Evx1 (V0v)
En1 (V1)
Chx10 (V2a)
Nkx2.2 (V3v)
dI3
MNs
dorsal
ventral
medial
lateral

**Figure S1**

**Figure S1. Swim-circuit architecture of *Xenopus* spinal cord**

**A, B.** Spinal interneurons along the brachial-thoracic-lumbar axis of stage NF47 *Xenopus laevis*: Evx1$^+$, V0v; En1$^+$, V1; Chx10$^+$, V2a; Nkx1.1$^+$, V3v; Lhx2-9$^+$, dI1; Isl1/2$^+$(dorsal), dI3; Cpne4$^+$, dI5; and Dmrt3$^+$, dI6. Scale bars, 50μm.

**C, D.** Normalised kernel density estimate maps summarising settling positions of swim interneurons as indicated. Region-combined concentric contours depict outermost to innermost 70th to 10th percentile, dorsal and ventral Isl1/2$^+$ contours are plotted separately. Dots represent normalized settling positions for: brachial (green), thoracic (yellow), lumbar (purple)

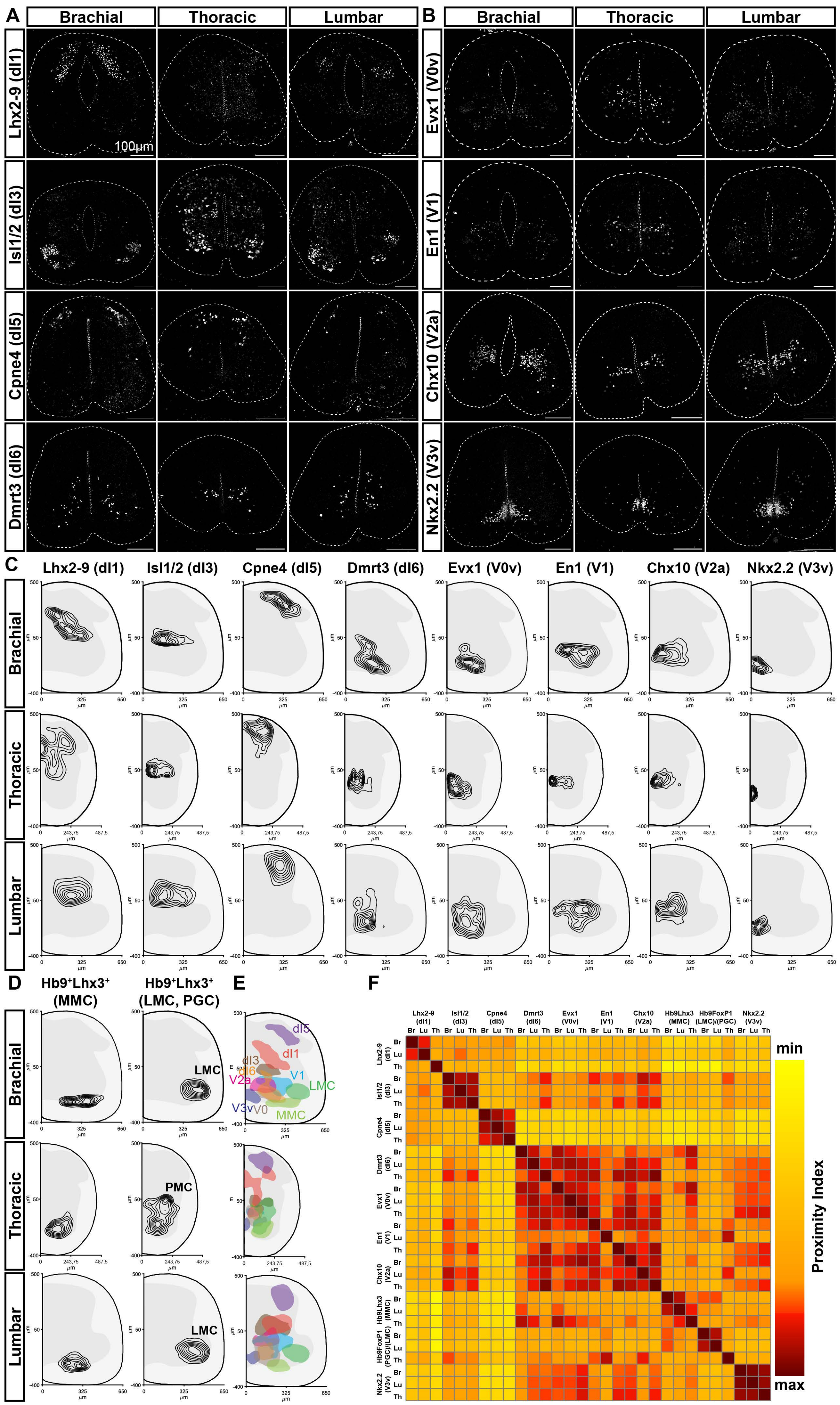
A
Brachial
Thoracic
Lumbar
Lhx2-9 (dI1)
Isl1/2 (dI3)
Cpne4 (dI5)
Dmrt3 (dI6)
100µm
B
Evx1 (V0v)
En1 (V1)
Chx10 (V2a)
Nkx2.2 (V3v)
C
Lhx2-9 (dI1)
Isl1/2 (dI3)
Cpne4 (dI5)
Dmrt3 (dI6)
Evx1 (V0v)
En1 (V1)
Chx10 (V2a)
Nkx2.2 (V3v)
D
Hb9+Lhx3+ (MMC)
Hb9+Lhx3+ (LMC, PGC)
LMC
PMC
E
dI5
dI1
dI3
dI6
V1
V2a
LMC
V3v
V0
MMC
F
min
max
Proximity Index
Br
Lu
Th

Figure S2

**Figure S2. Developing limb-circuit architecture of *Xenopus* spinal cord at NF55 tadpole stage**

**A, B.** Spinal interneurons along the brachial-thoracic-lumbar axis of stage NF55 *Xenopus laevis*: Evx1$^+$, V0v; En1$^+$, V1; Chx10$^+$, V2a; Nkx1.1$^+$, V3v; Lhx2-9$^+$, dI1; Isl1/2$^+$(dorsal), dI3; Cpne4$^+$, dI5; and Dmrt3$^+$, dI6. Scale bars, 100µm.

**C, D.** Normalised kernel density estimate maps summarising settling positions of motor and interneurons of the developing limb-circuit for brachial, thoracic, and lumbar regions, as indicated. Concentric contours depict outermost to innermost, 70th to 10th percentile.

**E.** Composite settling positions of motor and interneurons of the developing limb-circuit for brachial, thoracic, and lumbar regions. Shaded areas represent the 50th percentile of kernel density estimate map contours.

**F.** Proximity index heatmap comparing normalized kernel density estimate maps for developing limb-circuit neurons along the brachial-thoracic-lumbar axis of stage NF55 *Xenopus laevis*.

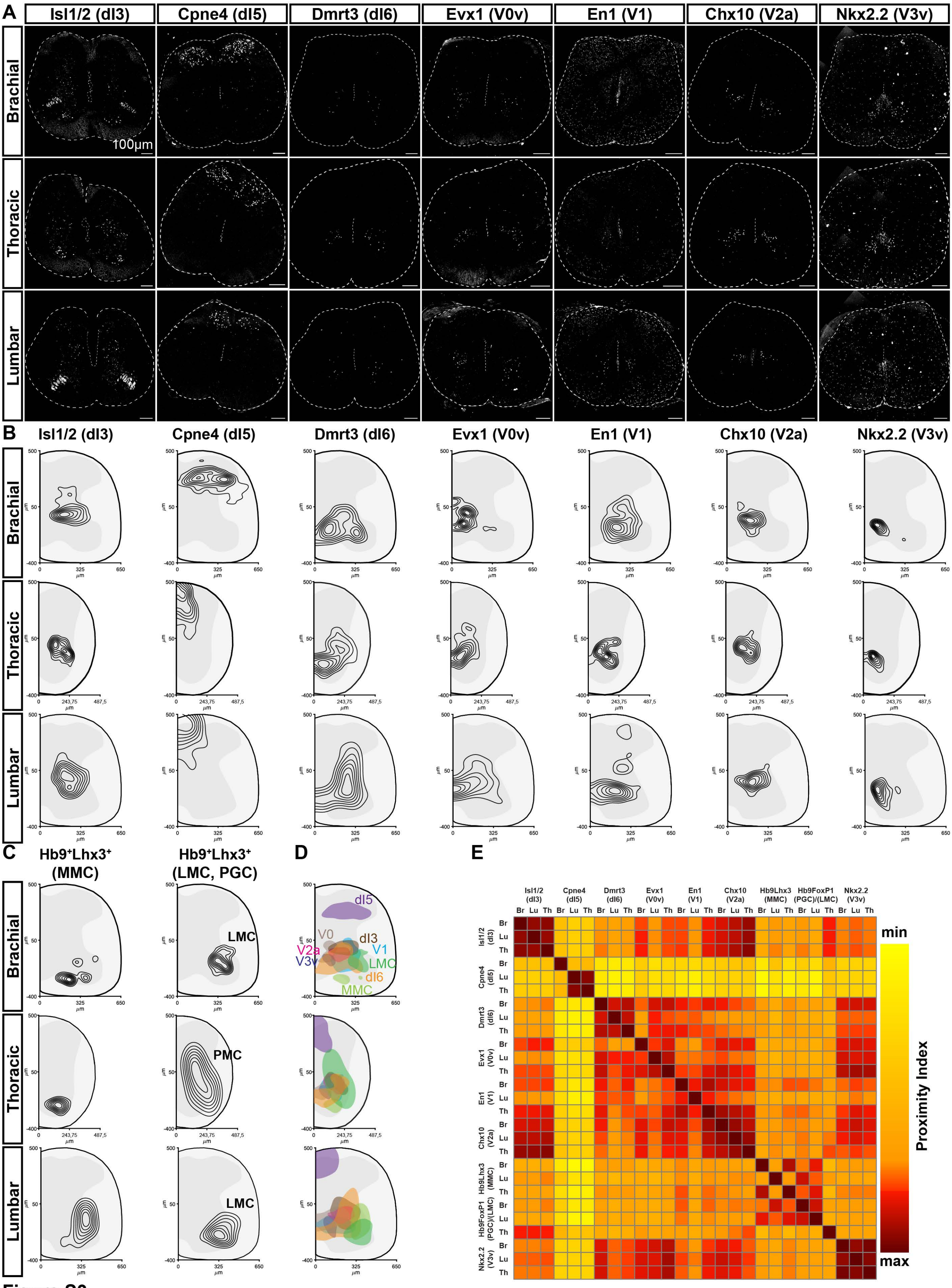

A
Isl1/2 (dI3)
Cpne4 (dI5)
Dmrt3 (dI6)
Evx1 (V0v)
En1 (V1)
Chx10 (V2a)
Nkx2.2 (V3v)
Brachial
Thoracic
Lumbar
100µm
B
C
Hb9⁺Lhx3⁺ (MMC)
Hb9⁺Lhx3⁺ (LMC, PGC)
LMC
PMC
D
dI5
V0
dI3
V2a
V1
V3v
LMC
dI6
MMC
E
Proximity Index
min
max


Figure S3

**Figure S3. Limb-circuit architecture of *Xenopus* spinal cord at NF66 frog stage**

**A.** Spinal interneurons along the brachial-thoracic-lumbar axis of stage NF66 *Xenopus laevis*: Evx1$^+$, V0v; En1$^+$, V1; Chx10$^+$, V2a; Nkx1.1$^+$, V3v; Isl1/2$^+$(dorsal), dI3; Cpne4$^+$, dI5; and Dmrt3$^+$, dI6. Scale bars, 100µm.

**B, C.** Normalised kernel density estimate maps summarising settling positions of motor and interneurons of the frog limb-circuit for brachial, thoracic, and lumbar regions, as indicated. Concentric contours depict outermost to innermost, 70th to 10th percentile.

**D.** Composite settling positions of motor and interneurons of the limb-circuit for brachial, thoracic, and lumbar regions. Shaded areas represent the 50th percentile of kernel density estimate map contours.

**E.** Proximity index heatmap comparing normalized kernel density estimate maps for developing limb-circuit neurons along the brachial-thoracic-lumbar axis of stage NF66 *Xenopus laevis*.

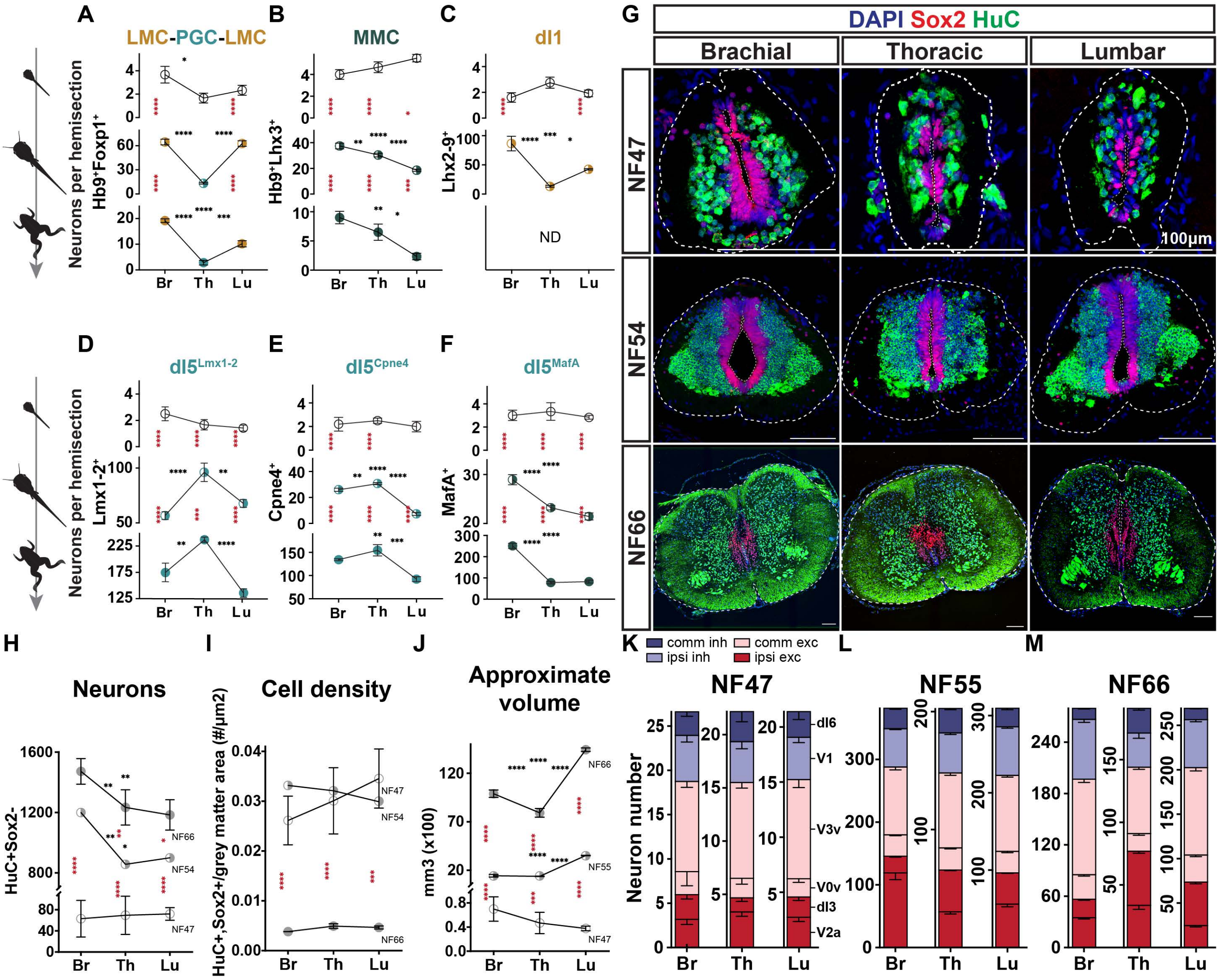

A
LMC-PGC-LMC
B
MMC
C
dI1
G
DAPI Sox2 HuC
Brachial
Thoracic
Lumbar
NF47
NF54
NF66
100μm
Neurons per hemisection
Hb9+Foxp1+
Hb9+Lhx3+
Lhx2-9+
ND
Br
Th
Lu
D
dI5Lmx1-2
E
dI5Cpne4
F
dI5MafA
Lmx1-2+
Cpne4+
MafA+
H
Neurons
HuC+Sox2-
NF66
NF54
NF47
I
Cell density
HuC+,Sox2+/grey matter area (#/μm2)
J
Approximate volume
mm3 (x100)
NF55
K
L
M
comm inh
ipsi inh
comm exc
ipsi exc
NF47
NF55
NF66
Neuron number
dI6
V1
V3v
V0v
dI3
V2a


Figure S4

**Figure S4. The swim-to-limb transition of the *Xenopus* spinal cord**

**A-F.** Neuron distribution along the brachial-thoracic-lumbar axis is uniform in the swim-circuit, but gets segmented with the emergence of the limb-circuit, acquiring a neuron class-distinct pattern. LMC/PGC, $Hb9^{+}FoxP1^{+}$; MMC, $Hb9+Lhx3^{+}$; dI1, $Lhx2\text{-}9^{+}$; $dI5^{Lmx1\text{-}2}$, $Lmx1\text{-}2^{+}$; $dI5^{MafA}$, $MafA^{+}$ ; $dI5^{Cpne4}$, $Cpne4^{+}$. For each animal 3-6 transverse hemisections per level were scored, with NF55;  N=12-14, NF47 and NF66; N=2-5 animals.

**G-I.** Cell counts of motor-related interneurons along the brachial-thoracic-lumbar axis of NF47 (**G**), NF55 (**H**), and NF66 (**I**) with regionalization within inhibitory and within excitatory neurons emerging only at the NF66 limb-circuit. Interneurons are sorted and colour-coded by projection pattern and function: blue, commissural inhibitory; light blue, ipsilateral inhibitory; pink, commissural excitatory; red, ipsilateral excitatory. All data reported as mean with SD.

**J.** Cells (DAPI), progenitors ($Sox2^{+}$) and neurons ($HuC^{+}$) along the brachial-thoracic-lumbar axis, over three locomotor modes (NF47, NF54 and NF66). Scale bars; 100μm.

**K.** Change in spinal cord volume per rostrocaudal region over three locomotor modes (NF47, NF55 and NF66), calculated by multiplying the mean transverse area at the middle of each rostrocaudal region with the mean length of that region. For each timepoint three (n=3) transverse sections per level were scored in 3 animals (N=3).

**L.** Neuron number changes per rostrocaudal region over three locomotor modes (NF47, NF54 and NF66) per transverse section at the middle of each rostrocaudal region.

**M.** Cell density changes per rostrocaudal region over three locomotor modes (NF47, NF54 and NF66) per transverse section at the middle of each rostrocaudal region. Cell density is calculated by dividing the sum of $HuC^{+}Sox2^{+}$, $HuC^{+}Sox2^{-}$, and $Sox2^{+}HuC^{-}$ cells by the area of the grey matter. Three (n=3) transverse sections per level were scored in 3 animals (N=3) for NF47 and NF66, for NF54, n=1 and N=1. (*also regarding* **L**). Black asterisks signify significant differences across rostrocaudal levels of a single developmental stage; red asterisks signify significant differences across developmental stages at the same rostrocaudal level. All data reported as mean ± SEM. One-way ANOVA and Tukey's multiple comparisons test; * p<0.1, ** p<0.01, *** p<0.001, **** p<0.0001. (*regarding* **A-F**, **K-M**).

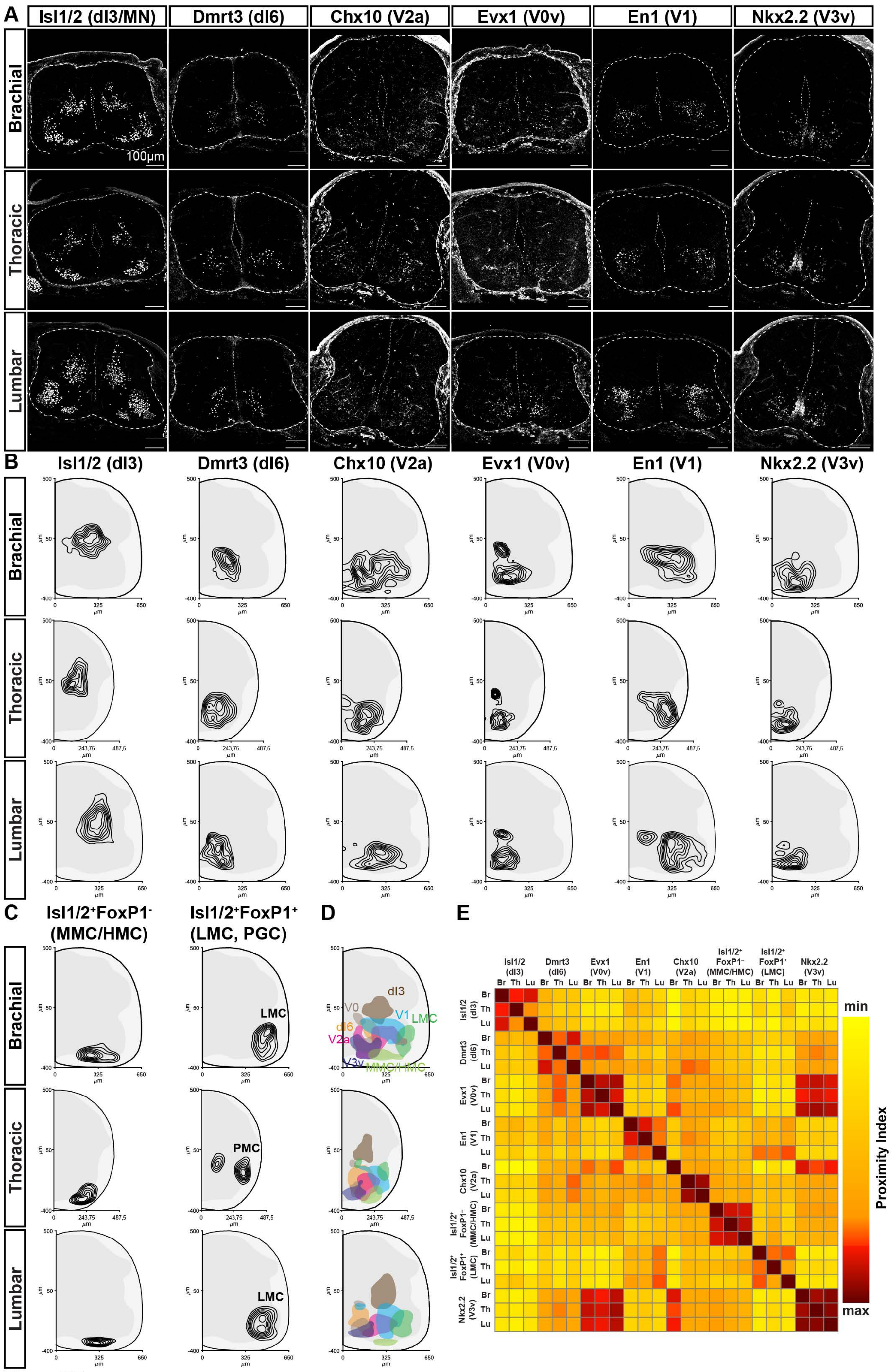
A
Isl1/2 (dI3/MN)
Dmrt3 (dI6)
Chx10 (V2a)
Evx1 (V0v)
En1 (V1)
Nkx2.2 (V3v)
Brachial
Thoracic
Lumbar
100μm
B
Isl1/2 (dI3)
Dmrt3 (dI6)
Chx10 (V2a)
Evx1 (V0v)
En1 (V1)
Nkx2.2 (V3v)
C
Isl1/2+FoxP1-
(MMC/HMC)
Isl1/2+FoxP1+
(LMC, PGC)
LMC
PMC
D
dI3
V0
V1
LMC
dI6
V2a
V3v
MMC/HMC
E
Isl1/2 (dI3)
Dmrt3 (dI6)
Evx1 (V0v)
En1 (V1)
Chx10 (V2a)
Isl1/2+ FoxP1− (MMC/HMC)
Isl1/2+ FoxP1+ (LMC)
Nkx2.2 (V3v)
Br
Th
Lu
min
max
Proximity Index

Figure S5

**Figure S5. Developing limb-circuit architecture of developing mouse spinal cord**

**A.** Spinal interneurons along the brachial-thoracic-lumbar axis of stage E13.5 mouse: Isl1/2$^+$(dorsal), dI3; Dmrt3$^+$, dI6; Chx10$^+$, V2a; Evx1$^+$, V0v; En1$^+$, V1; and Nkx1.1$^+$, V3v. Scale bars, 100μm.

**B, C.** Normalised kernel density estimate maps summarising settling positions of motor and interneurons of the developing limb-circuit for brachial, thoracic, and lumbar regions, as indicated. Concentric contours depict outermost to innermost, 70th to 10th percentile.

**D.** Composite settling positions of motor and interneurons of the developing limb-circuit for brachial, thoracic, and lumbar regions. Shaded areas represent the 50th percentile of kernel density estimate map contours.

**E.** Proximity index heatmap comparing normalized kernel density estimate maps for developing limb-circuit neurons along the brachial-thoracic-lumbar axis of stage E13.5 mouse.

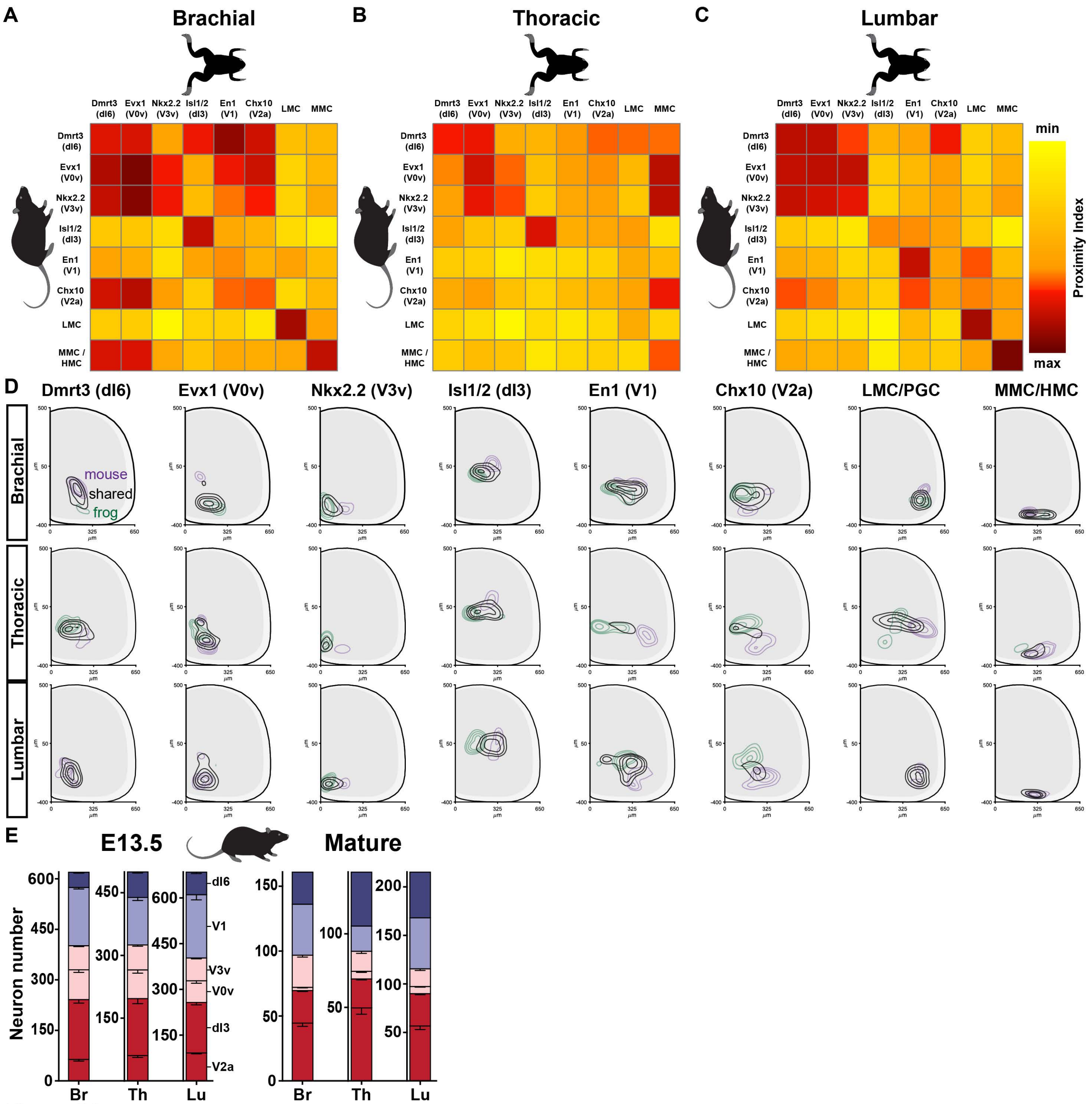

A
Brachial
B
Thoracic
C
Lumbar
Dmrt3 (dI6)
Evx1 (V0v)
Nkx2.2 (V3v)
Isl1/2 (dI3)
En1 (V1)
Chx10 (V2a)
LMC
MMC
MMC / HMC
min
max
Proximity Index
D
Dmrt3 (dI6)
Evx1 (V0v)
Nkx2.2 (V3v)
Isl1/2 (dI3)
En1 (V1)
Chx10 (V2a)
LMC/PGC
MMC/HMC
Brachial
Thoracic
Lumbar
mouse
shared
frog
E
E13.5
Mature
Neuron number
dI6
V1
V3v
V0v
dI3
V2a
Br
Th
Lu


Figure S6

**Figure S6. Rostrocaudal and cross-species differences in neural distributions in frog versus mouse**

**A-C.** Proximity index heatmaps comparing normalized kernel density estimate maps for developing limb-circuit neurons for brachial (**A**), thoracic (**B**), and lumbar (**C**) regions of stage NF55 *Xenopus* (horizontal axis) and E13.5 mouse (vertical axis). Frog kernel density estimate maps were resized so that the gray-white matter border matched that of the mouse spinal cord.

**D.** Normalised kernel density estimate maps summarising settling positions of motor and interneurons of developing limb-circuits for brachial, thoracic, and lumbar regions, as indicated. Black contours depict normalised kernel density estimate maps of shared spatial probabilities between mouse and frog; purple and green contours depict normalised kernel density estimate maps of mouse and frog spatial probabilities without those shared between them, respectively. Within each plot, black, purple and green maps are plotted on the same scale, so that concentric contours depict outermost (of all three maps) to innermost, 70th, 50th, 30th and 10th percentiles. Absent inner concentric contours signify that this map did not pass the higher/denser thresholds, of the 10th or 30th or even 50th percentile.

**E.** Cell counts of motor-related interneurons along the brachial-thoracic-lumbar axis of E13.5 and mature (E18-p149) mice, with regionalization within inhibitory neurons emerging only at the mature limb-circuit. Interneurons are sorted and colour-coded by projection pattern and function: blue, commissural inhibitory; light blue, ipsilateral inhibitory; pink, commissural excitatory; red, ipsilateral excitatory. Data reported as mean with SD (were available).

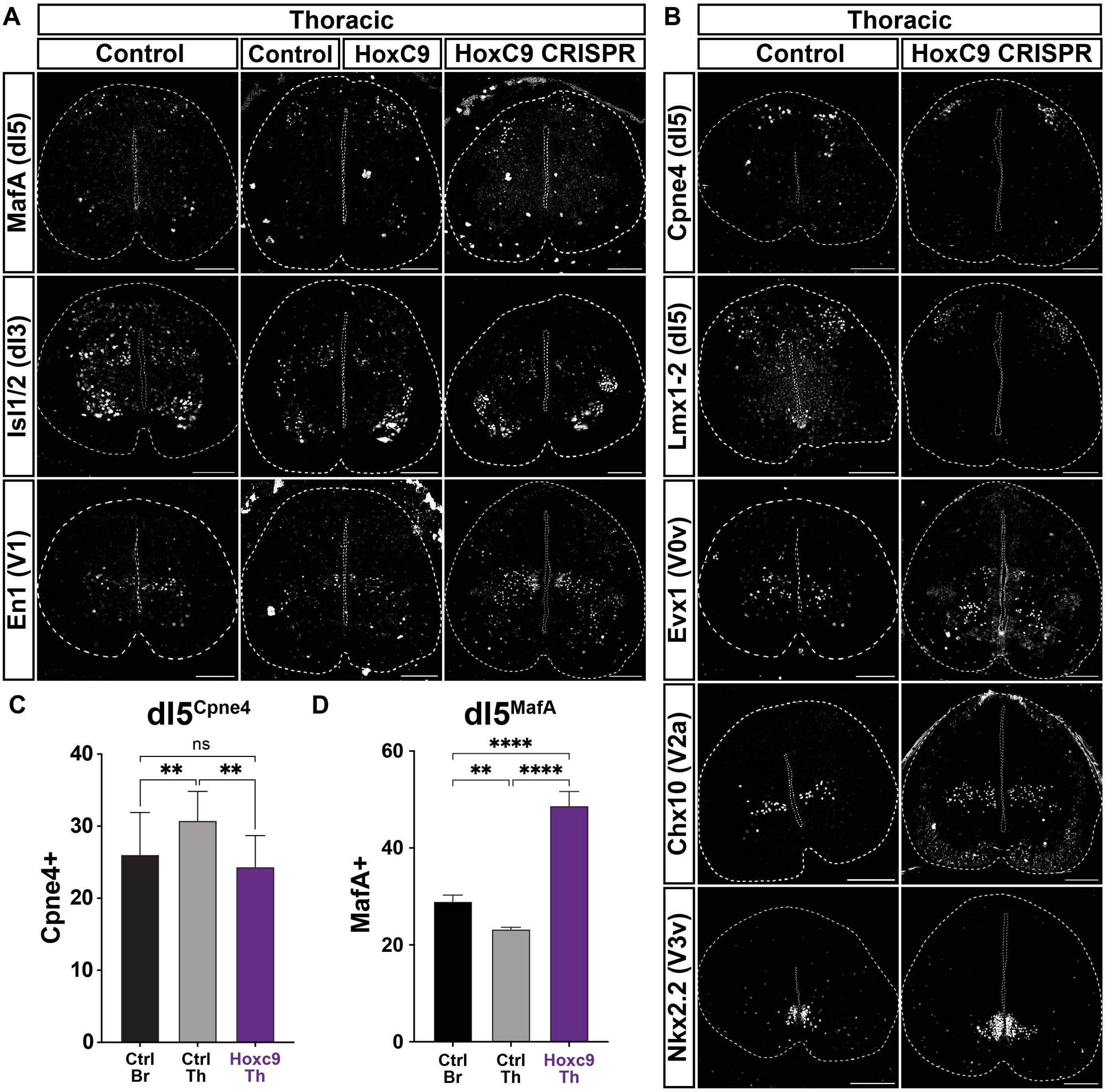
A
Thoracic
Control
Control
HoxC9
HoxC9 CRISPR
MafA (dI5)
Isl1/2 (dI3)
En1 (V1)
B
Thoracic
Control
HoxC9 CRISPR
Cpne4 (dI5)
Lmx1-2 (dI5)
Evx1 (V0v)
Chx10 (V2a)
Nkx2.2 (V3v)
C
dI5Cpne4
Cpne4+
0
10
20
30
40
ns
**
**
Ctrl Br
Ctrl Th
Hoxc9 Th
D
dI5MafA
MafA+
0
20
40
60
****
**
****
Ctrl Br
Ctrl Th
Hoxc9 Th

Figure S7

**Figure S7. Hoxc9-driven transformation of thoracic interneuron architecture**

**A.** Spinal interneurons at the thoracic region of control (left), half- (middle), and whole-body mutant (right) NF55 *Xenopus laevis* animals. Neuron distributions in the thoracic spinal cord of whole-mutants (CRISPR/Cas 9, *hoxc9* gRNA injected) and the injected side of half-mutants are increased for: MafA$^+$(dorsal), dI5; and En1$^+$, V1, but decreased for: Isl1/2$^+$(dorsal), dI3.

**B.** Spinal interneurons at the thoracic region of control (left) and whole-body mutant (right) NF55 *Xenopus laevis* animals. Neuron distributions in the thoracic spinal cord of whole-mutants (CRISPR/Cas 9, *hoxc9* gRNA injected) are decreased for: Cpne4$^+$, dI5; Lmx1-2$^+$, dI5, but increased for: Evx1$^+$, V0v; Chx10$^+$, V2a; and Nkx1.1$^+$, V3v.

Scale bars, 100μm.

**C, D.** Thoracic-to-brachial-like transformation of neuron numbers, showing two dI5 interneuron subpopulations in control animals at brachial and thoracic levels, and whole-body CRISPR animals at thoracic levels of stage NF55 animals. All data reported as mean ± SEM. One-way ANOVA; ns, $p>0.1$; **, $p<0.01$; ****, $p<0.0001$. For each animal 3-6 transverse hemisections per level were scored, with N=3-12 for control, and N=3-5 for *hoxc9* mutant animals.

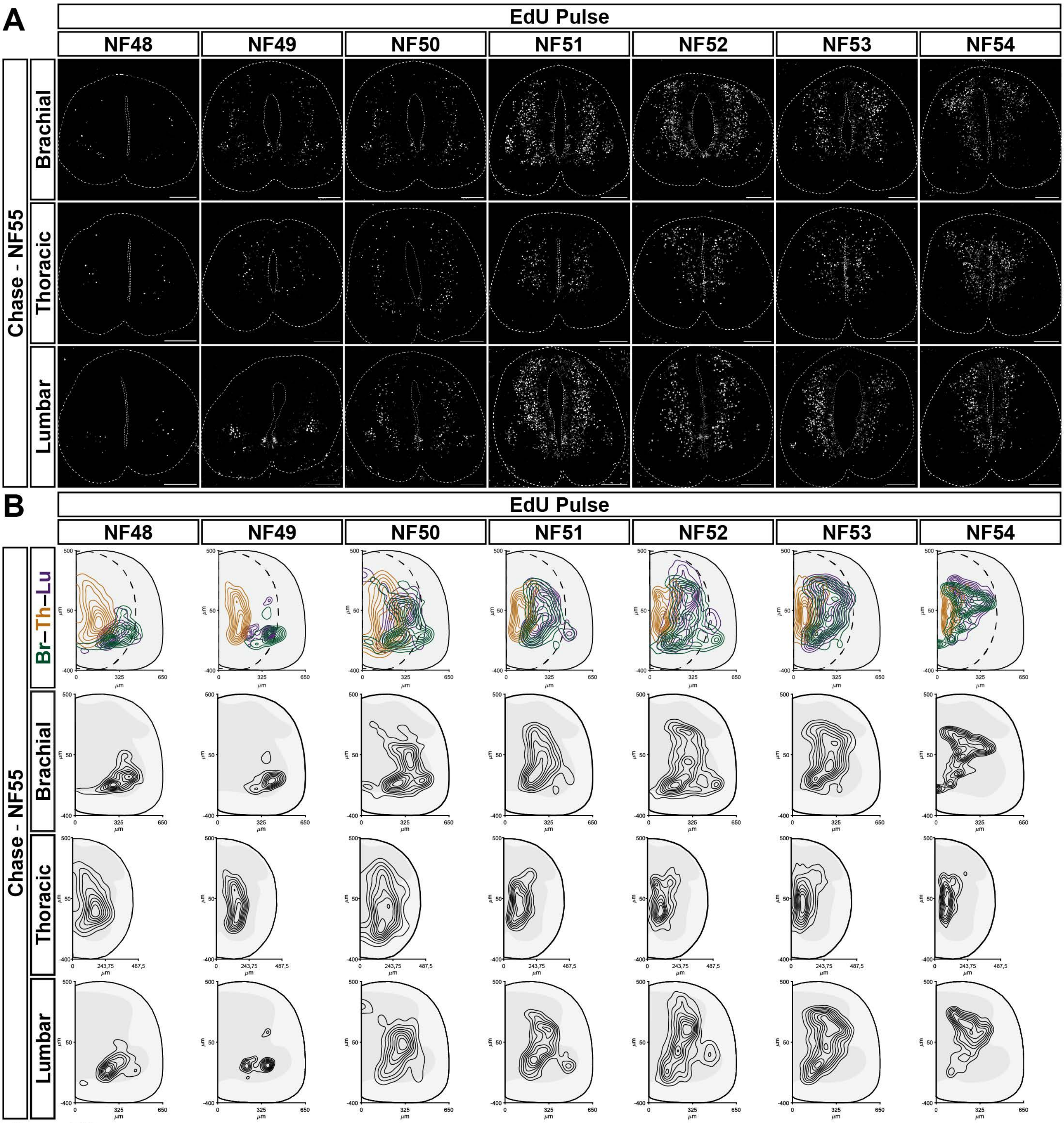
A
EdU Pulse
NF48
NF49
NF50
NF51
NF52
NF53
NF54
Chase - NF55
Brachial
Thoracic
Lumbar
B
EdU Pulse
NF48
NF49
NF50
NF51
NF52
NF53
NF54
Chase - NF55
Br–Th–Lu
Brachial
Thoracic
Lumbar
µm

Figure S8

**Figure S8. Limb-circuit neurogenesis along the rostrocaudal axis of the frog spinal cord**
**A.** $EdU^+$ cells along the brachial-thoracic-lumbar axis of *Xenopus laevis* animals that were injected at stages NF48 to 54 (columns, as indicated), and chased out to stage NF55. Scale bars, 100µm.
**B.** Normalised kernel density estimate maps summarising settling positions of $EdU^+$ cells for the brachial, thoracic, and lumbar regions, as well as composite (top row: brachial, green; thoracic, yellow; lumbar, purple), as indicated. In composite maps, thoracic settling positions are plotted on a narrower spinal cord with dotted lines indicating the thoracic and full lines the brachial/lumbar border. Concentric contours depict outermost to innermost, 70th to 10th percentile.

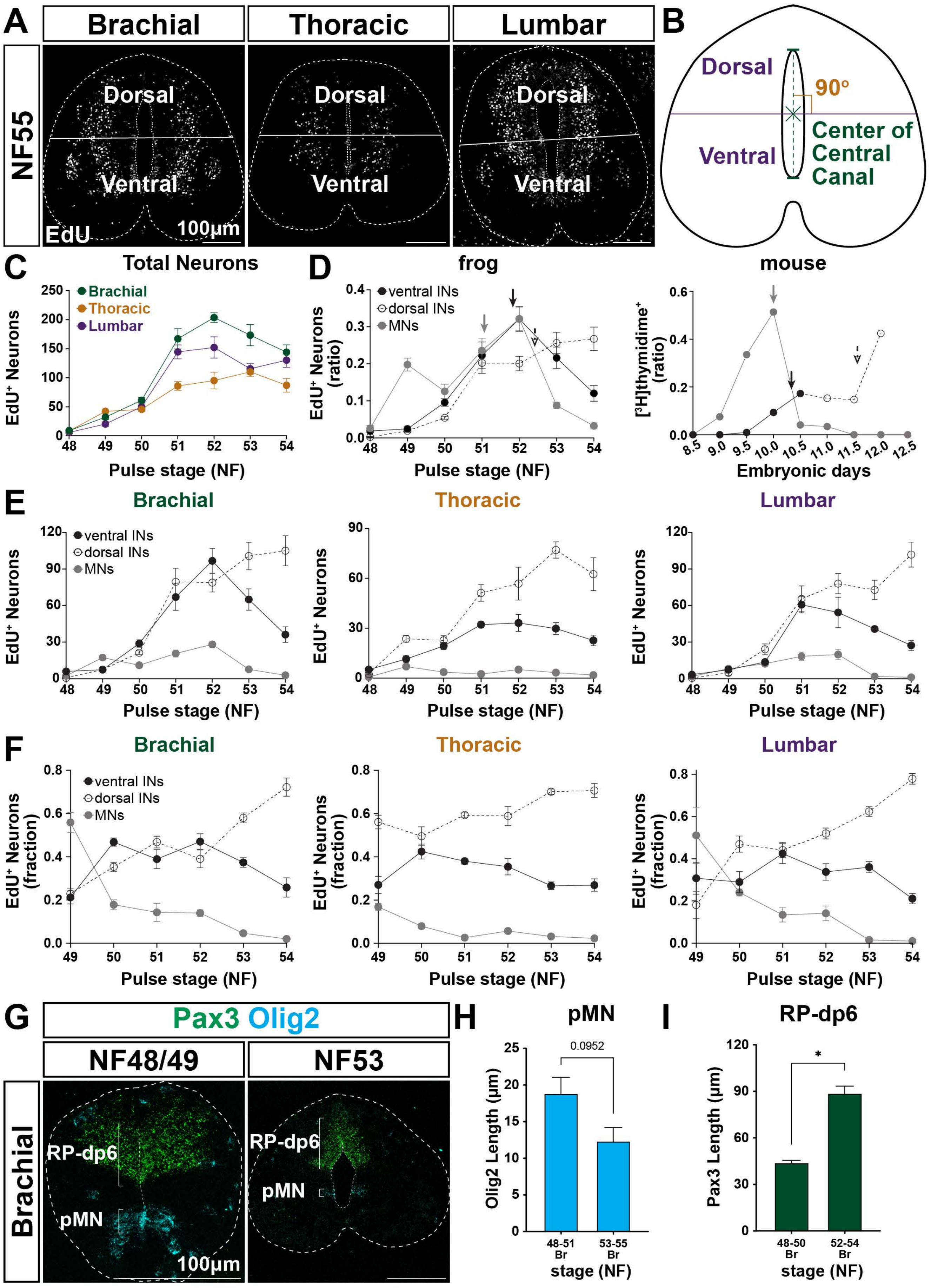
A
Brachial
Thoracic
Lumbar
NF55
Dorsal
Ventral
EdU
100µm
B
Dorsal
90°
Ventral
Center of Central Canal
C
Total Neurons
Brachial
Thoracic
Lumbar
EdU+ Neurons
Pulse stage (NF)
D
frog
ventral INs
dorsal INs
MNs
EdU+ Neurons (ratio)
Pulse stage (NF)
mouse
[3H]thymidime+ (ratio)
Embryonic days
E
Brachial
Thoracic
Lumbar
EdU+ Neurons
Pulse stage (NF)
F
Brachial
Thoracic
Lumbar
EdU+ Neurons (fraction)
Pulse stage (NF)
G
Pax3 Olig2
NF48/49
NF53
Brachial
RP-dp6
pMN
100µm
H
pMN
0.0952
Olig2 Length (µm)
48-51 Br
53-55 Br
stage (NF)
I
RP-dp6
*
Pax3 Length (µm)
48-50 Br
52-54 Br
stage (NF)

Figure S9

**Figure S9. Regionalized spinal neurogenesis of the limb-circuit**

**A.** Brachial, thoracic and lumbar examples of NF55 *Xenopus laevis* spinal cord sections with $EdU^+$ cells depicting the dorso-ventral boundary (white line). Scale bars, 100µm.

**B.** Schematic of a transverse section of the spinal cord with the dorso-ventral boundary drawn at the midpoint between the dorsal and ventral edge of the ventricle (center of central canal, marked with ‘x’) perpendicular to the dorso-ventral axis.

**C.** Synchronous rostrocaudal neurogenesis, exemplified by $EdU^+$ cells at brachial (green), thoracic (yellow), and lumbar (purple) levels of the spinal cord. All data reported as mean ± SEM. For each timepoint three (n=3) transverse hemisections per level were scored in six (N=6) animals.

**D.** Overlapping versus distinct phases of limb-circuit generation in the brachial region of *Xenopus* and mouse, respectively, depicting motor (gray), ventral (black, full circles) and dorsal (black, open circles/dotted line) interneurons. Mouse data are adapted from *Sims & Vaughn 1979*.[136] Arrows indicate $50^{th}$ percentile of respective neurogenesis. Ratios are calculated dividing by the sum of all timepoints.

**E.** Absolute brachial, thoracic and lumbar neurogenesis of the limb-circuit of *Xenopus* split in motor (gray), ventral (black, full circles) and dorsal (black, open circles/dotted line) interneurons.

**F.** Fractional brachial, thoracic and lumbar neurogenesis of the limb-circuit of *Xenopus* split in motor (gray), ventral (black, full circles) and dorsal (black, open circles/dotted line) interneurons. Fractions are calculated dividing by the sum of each timepoint.

(**C-F**) All data reported as mean ± SEM. For each timepoint three (n=3) transverse hemisections per level were scored in two (N=2) animals.

**G.** Detection of Pax3 and Olig2 mRNA at brachial levels of stage NF48/49 and NF53 *Xenopus laevis* animals using in situ hybridization indicates the dorsal (roof plate-to-dp6), and motor (pMN) progenitor domains, respectively. Scale bars, 100µm.

**H-I.** Progenitor domain size changes between early and late limb-circuit generation, based on Pax3 and Olig2 mRNA detection at brachial levels of *Xenopus laevis* animals at stage NF48-50 versus NF52-54 (dorsal, RP-dp6) and NF48-51 versus NF53-55 (motor, pMN). All data reported as mean ± SEM. Mann-Whitney; *, $p<0.05$. For each time window two (n=2) transverse sections were scored in four to five (N=4-5) animals.

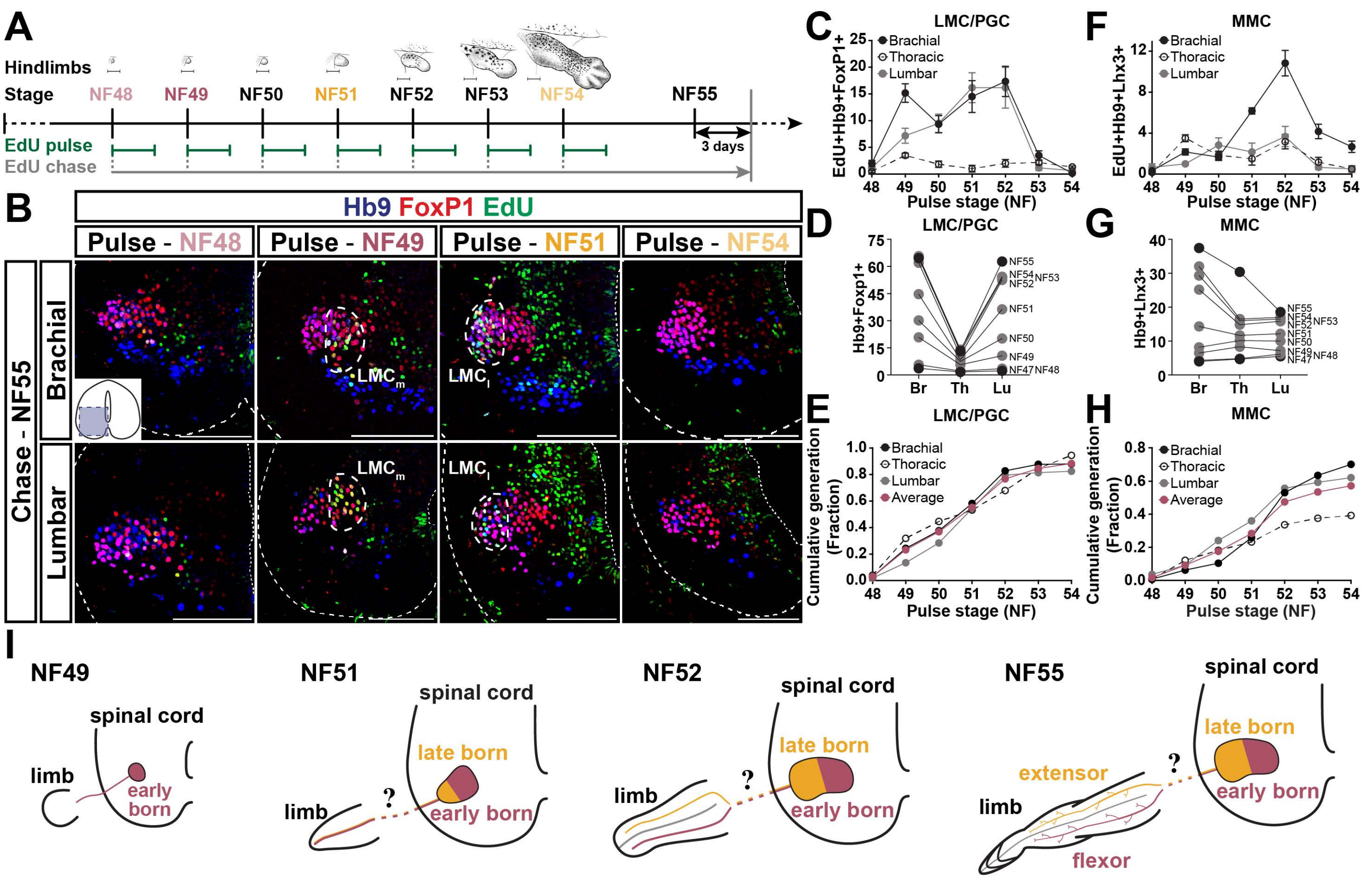

A
Hindlimbs
Stage
NF48
NF49
NF50
NF51
NF52
NF53
NF54
NF55
EdU pulse
EdU chase
3 days
B
Hb9 FoxP1 EdU
Pulse - NF48
Pulse - NF49
Pulse - NF51
Pulse - NF54
Chase - NF55
Brachial
Lumbar
LMCm
LMCl
C
LMC/PGC
EdU+Hb9+FoxP1+
Brachial
Thoracic
Lumbar
Pulse stage (NF)
D
LMC/PGC
Hb9+Foxp1+
Br
Th
Lu
E
LMC/PGC
Cumulative generation (Fraction)
Average
F
MMC
EdU+Hb9+Lhx3+
G
MMC
Hb9+Lhx3+
H
MMC
I
NF49
spinal cord
limb
early born
NF51
late born
NF52
NF55
extensor
flexor


Figure S10

**Figure S10. Regionalized generation of motor neurons**

**A.** Schematic of EdU pulse-chase experiment. Each pulse represents a 3-day injection regiment. Hindlimb drawings adapted from *Zahn et al. 2022*.[182]

**B.** $LMC_m$ neurogenesis precedes that of $LMC_l$ synchronously for brachial and lumbar regions in the *Xenopus* frog. Images depict the ventral horn, as indicated in the inset, of stage NF55 animals pulsed with EdU at NF48, 49, 51, and 53. Scale bars, 100μm.

**C, F.** Number of LMC/PGC and MMC neurons born at each indicated pulse based on EdU uptake and $Hb9^+FoxP1^+$ and $Hb9^+Lhx3^+$ colabeling, respectively. LMC neurons exhibit a burst-like generation pattern at limb, as do brachial MMC, but PGC, and thoracic and lumbar MMC neurogenesis is maintained at a low and stable rate. All data reported as mean ± SEM. For each timepoint three (n=3) transverse hemisections per level were scored in two (N=2) animals.

**D, G.** Regionalization of rostrocaudal patterns. Black points depict the mean for the indicated stages (see **Figure S1**), and gray points represent the NF47 mean plus the mean of neurons born up to the indicated stage (shown in **C**, **F**).

**E, H.** Cumulative fraction of LMC/PGC and MMC neuron generation between stages NF48 and NF54 for brachial, thoracic, and lumbar regions and their average, colour-coded as indicated. Fractions are calculated by adding the generation means up to the indicated timepoint and dividing by the mean at stage NF55.

**I.** Schematic of the sequential $LMC_m$ and $LMC_l$ generation and hypothesized birth time-based limb innervation split into flexor (ventral) and extensor (dorsal) compartments.

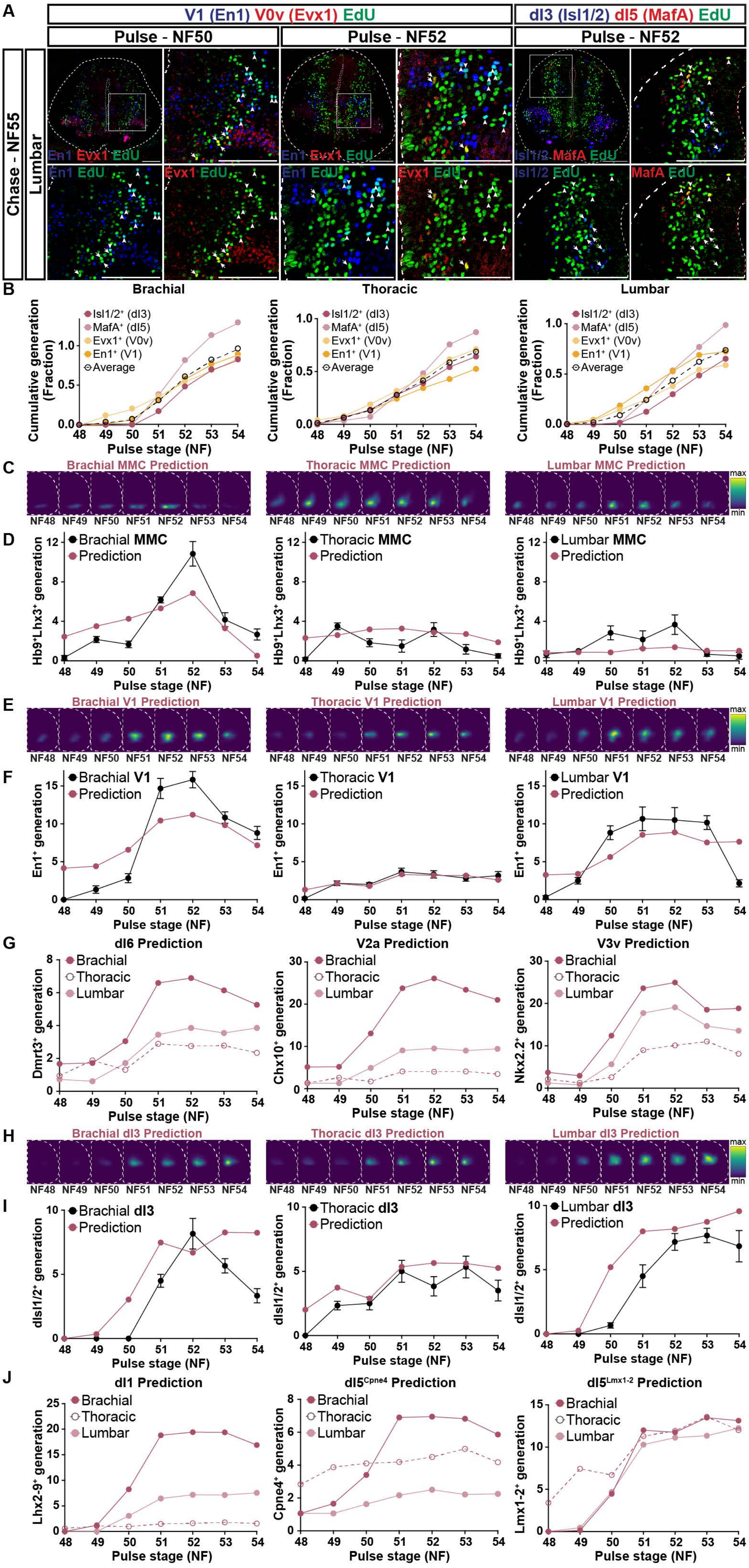

A
V1 (En1) V0v (Evx1) EdU
dI3 (Isl1/2) dI5 (MafA) EdU
Pulse - NF50
Pulse - NF52
Pulse - NF52
Chase - NF55
Lumbar
En1 Evx1 EdU
En1 EdU
Evx1 EdU
Isl1/2 MafA EdU
Isl1/2 EdU
MafA EdU
B
Brachial
Thoracic
Lumbar
Isl1/2+ (dI3)
MafA+ (dI5)
Evx1+ (V0v)
En1+ (V1)
Average
Cumulative generation (Fraction)
Pulse stage (NF)
C
Brachial MMC Prediction
Thoracic MMC Prediction
Lumbar MMC Prediction
NF48 NF49 NF50 NF51 NF52 NF53 NF54
max
min
D
Brachial MMC
Thoracic MMC
Lumbar MMC
Prediction
Hb9+Lhx3+ generation
E
Brachial V1 Prediction
Thoracic V1 Prediction
Lumbar V1 Prediction
F
Brachial V1
Thoracic V1
Lumbar V1
En1+ generation
G
dI6 Prediction
V2a Prediction
V3v Prediction
Brachial
Thoracic
Lumbar
Dmrt3+ generation
Chx10+ generation
Nkx2.2+ generation
H
Brachial dI3 Prediction
Thoracic dI3 Prediction
Lumbar dI3 Prediction
I
Brachial dI3
Thoracic dI3
Lumbar dI3
dIsl1/2+ generation
J
dI1 Prediction
dI5Cpne4 Prediction
dI5Lmx1-2 Prediction
Lhx2-9+ generation
Cpne4+ generation
Lmx1-2+ generation


Figure S11

**Figure S11. Predicted regionalization of interneuron generation**

**A.** Different numbers of V0v, V1, dI3, and $dI5^{MafA}$ interneurons are born at NF50 and NF52 stages at lumbar levels, marked by EdU and colabeled with Evx1 (arrows), En1 (arrowheads), $Isl1/2_{dorsal}$ (arrows), and $MafA_{dorsal}$ (arrowheads), respectively. Scale bars, 100µm.

**B.** Cumulative fraction of $Isl1/2^{+}$(dorsal), dI3; $MafA^{+}$(dorsal), dI5; $Evx1^{+}$, V0v; and $En1^{+}$, V1 interneuron generation between stages NF48 and 54 for brachial, thoracic, and lumbar regions and their average, colour-coded as indicated. Fractions are calculated by adding the generation means up to the indicated timepoint and dividing by the mean at stage NF55.

**C, E, H.** Normalised kernel density estimate maps of MMC (**C**), V1 (**E**), and dI3 (**H**) predicted to be generated at each timepoint between NF48 and 54, as indicated, for brachial, thoracic and lumbar regions, by combining normalized KDEs of EdU uptake per timepoint and of the respective neuron populations at stage NF55 (see **Figures S3, S8**).

**D, F, I.** Predicted (dark pink) and actual (black, see **Figures 4, S10**) number of MMC (**D**), V1 (**F**), and dI3 (**I**) neurons born at each indicated pulse. Predicted numbers are calculated based on normalized KDEs of predicted neurogenesis per region and per timepoint (shown in **C**, **E**, **H**), the total cumulative generation between NF48 and 54 (see **B**, and also **Figure S10H**) and the mean population size at stage NF55 (see **Figure 1E, 1G** and **S1B**).

**G, J.** Predicted number of $Dmrt3^{+}$, dI6; $Chx10^{+}$, V2a; $Nkx1.1^{+}$, V3v (**G**) and $Lhx2\text{-}9^{+}$, dI1; $Cpne4^{+}$, dI5; $Lmx1\text{-}2^{+}$, dI5 (**F**) neurons born at each indicated pulse, for brachial (dark pink), thoracic (open circles), and lumbar (pink). Predicted numbers are calculated based on normalized KDEs of predicted neurogenesis per region and per timepoint, the average cumulative generation between NF48 and 54 and the mean population size at stage NF55.

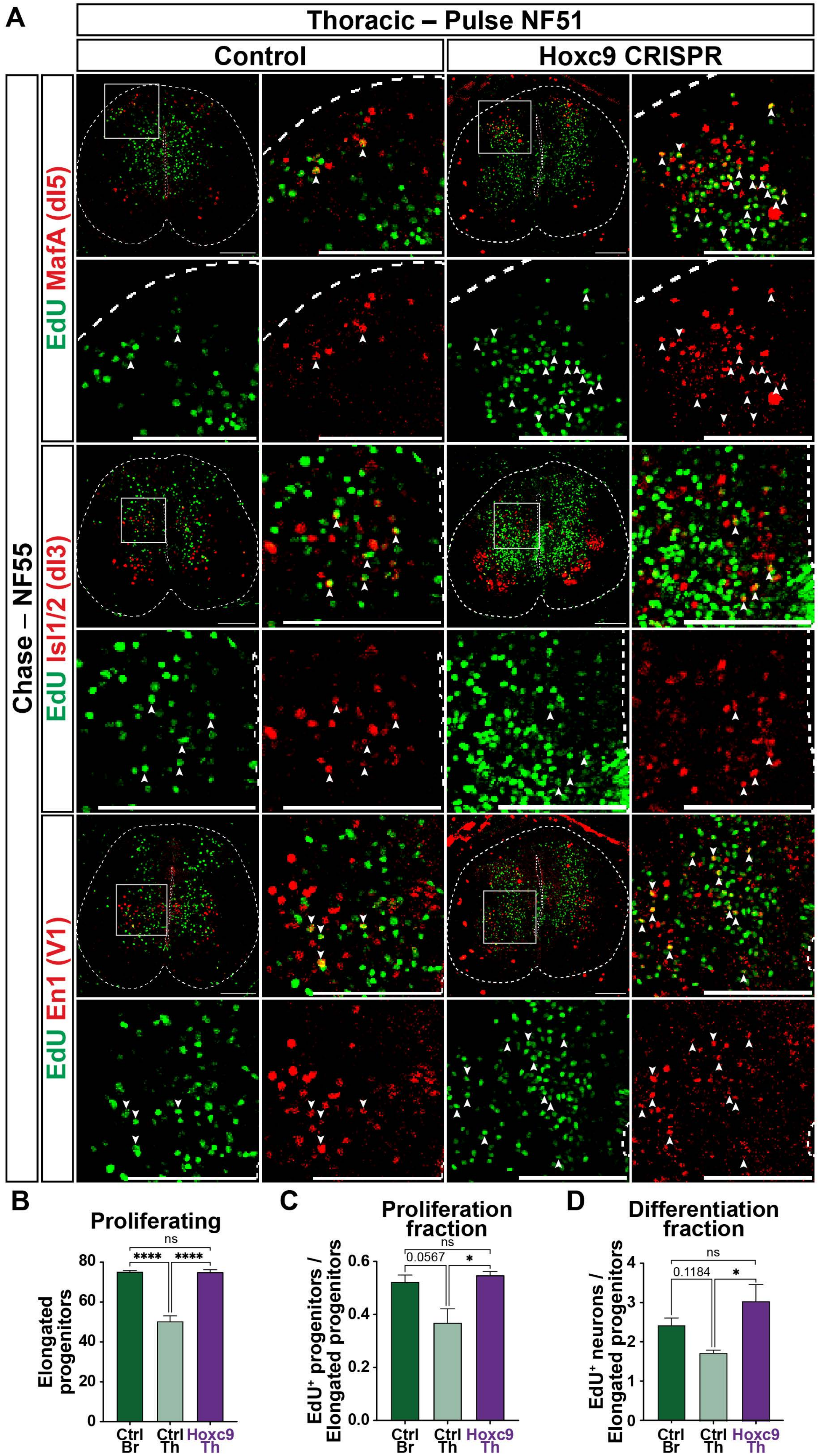
A
Thoracic – Pulse NF51
Control
Hoxc9 CRISPR
Chase – NF55
EdU MafA (dI5)
EdU Isl1/2 (dI3)
EdU En1 (V1)
B
Proliferating
Elongated progenitors
Ctrl Br
Ctrl Th
Hoxc9 Th
ns
****
****
C
Proliferation fraction
EdU⁺ progenitors / Elongated progenitors
0.0567
ns
*
D
Differentiation fraction
EdU⁺ neurons / Elongated progenitors
0.1184
ns
*

Figure S12

**Figure S12. *hoxc9* loss-of-function transforms neurogenesis of thoracic interneurons**

**A.** Thoracic-to-brachial-like transformation of progenitor expansion, neurogenesis and neural architecture at NF55 stage frogs. Cells born at stage NF51 at thoracic level, in control and whole-body *hoxc9* CRISPR animals, are marked by EdU and colabeled with $MafA^+$, dI5; (dorsal)$Isl1/2^+$, dI3; $En1^+$, V1 interneuron markers. Scale bars, 100μm.

**B.** The subset of elongated progenitors lining the central canal.

**C.** The proliferation fraction between NF51-55 is calculated by dividing the number of $EdU^+$ progenitors at NF55 by the number of progenitors lining the central canal (mitotically active/elongated progenitors).

**D.** The differentiation fraction between NF51-55 is calculated by dividing the number of $EdU^+$ neurons at NF55 by the number of progenitors lining the central canal (mitotically active/elongated progenitors).

**A**

##### Network Parameters

| | | Neurons | Excitatory fraction | Vestibular Targets | Sparsity | $f_{max}$ | threshold | gain |
|---|---|---|---|---|---|---|---|---|
| Frog Thoracic | Wildtype | 4096 | ~ 69% | 300 | ~ 90% | 60 | 20 | 1 |
| | Brachialized | 9169 | ~ 76% | 300 | ~ 90% | 60 | 20 | 1 |
| Mouse Thoracic | Wildtype | 4096 | ~ 49% | 300 | ~ 94% | 60 | 20 | 1 |

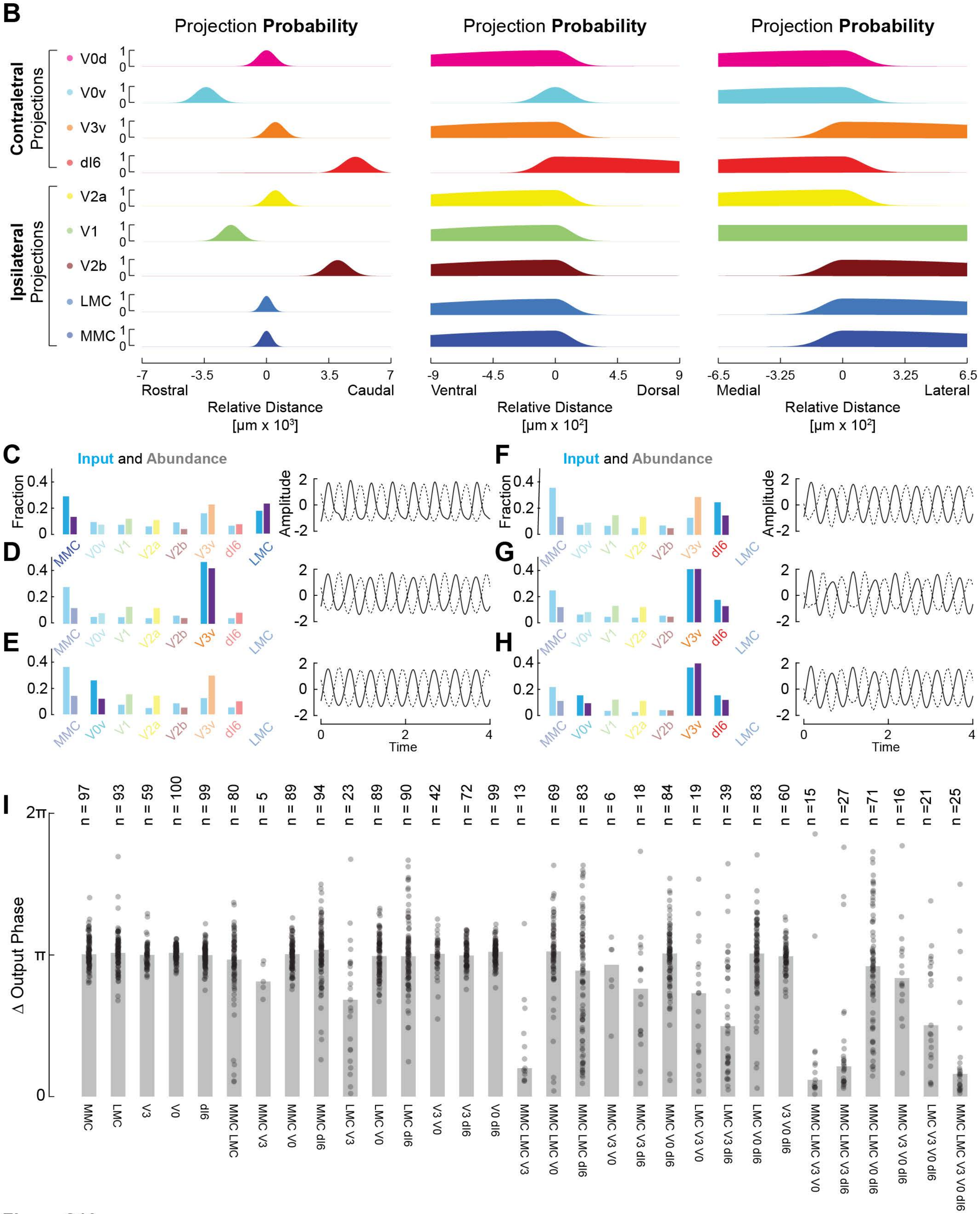


**Figure S13**

**Figure S13. Thoracic network model of thoracic *hoxc9* loss-of-function**

**A.** Model parameters for frog and mouse networks. $\mathbf{f_{max}}$, **threshold** and **gain** refer to parameters of the network equation used in simulations (*see* **STAR Methods**). For tonic drive simulations, the wildtype thoracic network neuron number used was 2*4096.

**B.** Spatial projection probabilities across rostrocaudal-, mediolateral and dorsoventral dimensions for all motor- and interneuron types used across networks. The final synapse probability emerges as the geometric mean of these projection biases (*see* **STAR Methods**).

**C-H.** Examples of partial network brachializations that do not induce bilateral synchrony in response to vestibulospinal input. Wildtype cell types are color coded as in **B,** brachialized cell types in purple, with bars denoting their numerical fraction in the particular brachialization. Blue bars denote the fractional distribution of vestibulospinal targets.

**I.** Distribution of left-right phase differences for 31 partial brachializations (all combinations of motor neurons and commissural neurons). Bar height denotes median. n is the number of data points out of 100 instantiations/brachialization.

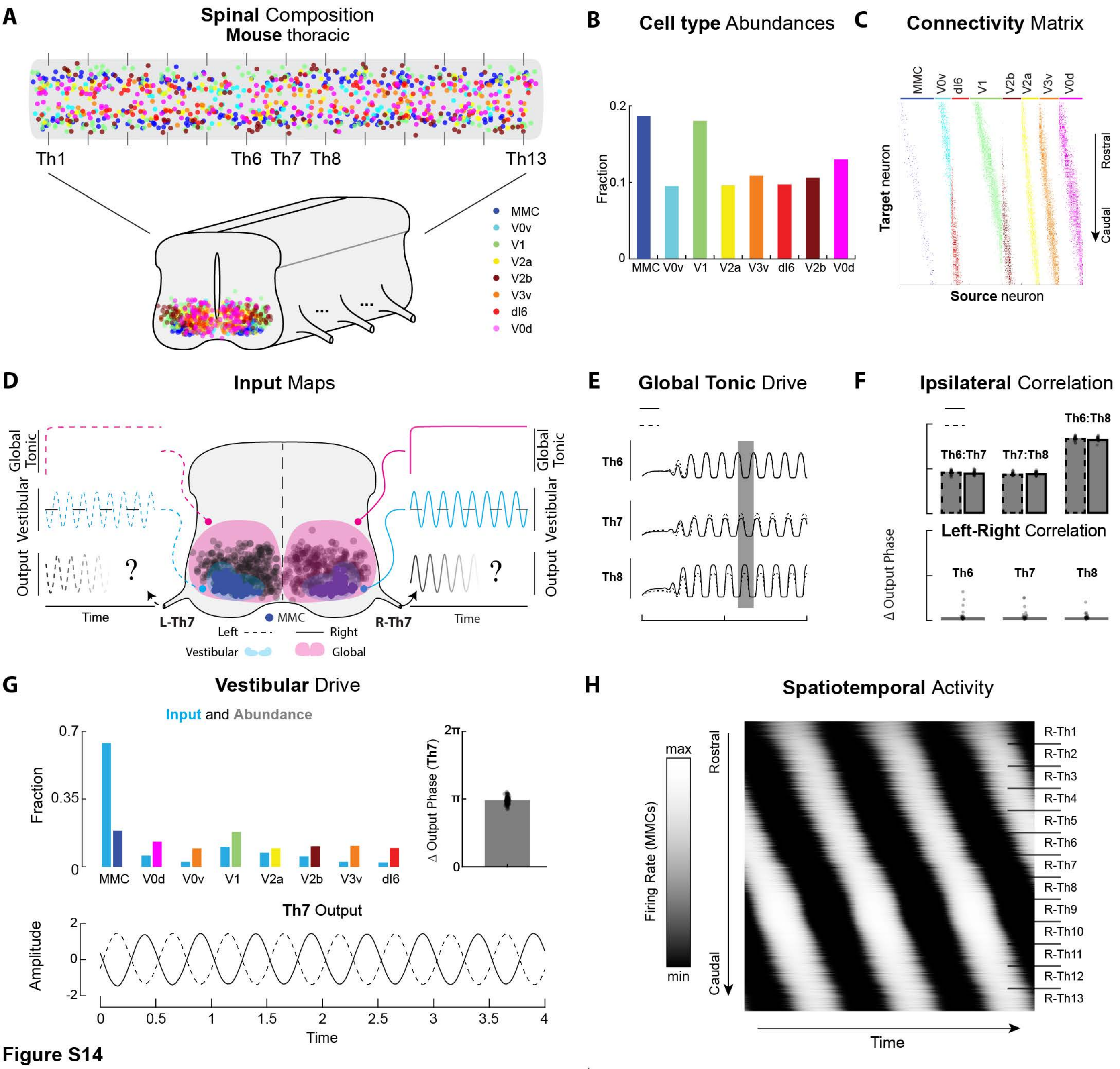
A
Spinal Composition
Mouse thoracic
Th1
Th6 Th7 Th8
Th13
MMC
V0v
V1
V2a
V2b
V3v
dI6
V0d
B
Cell type Abundances
Fraction
0.2
0.1
0
MMC V0v V1 V2a V3v dI6 V2b V0d
C
Connectivity Matrix
MMC
V0v
dI6
V1
V2b
V2a
V3v
V0d
Target neuron
Source neuron
Rostral
Caudal
D
Input Maps
Global Tonic
Vestibular
Output
Time
L-Th7
R-Th7
MMC
Left
Right
Vestibular
Global
E
Global Tonic Drive
Th6
Th7
Th8
F
Ipsilateral Correlation
Th6:Th7
Th7:Th8
Th6:Th8
Left-Right Correlation
Δ Output Phase
Th6
Th7
Th8
G
Vestibular Drive
Input and Abundance
Fraction
0.7
0.35
0
MMC V0d V0v V1 V2a V2b V3v dI6
2π
π
0
Δ Output Phase (Th7)
Th7 Output
Amplitude
2
0
-2
0 0.5 1 1.5 2 2.5 3 3.5 4
Time
H
Spatiotemporal Activity
max
min
Firing Rate (MMCs)
Rostral
Caudal
R-Th1
R-Th2
R-Th3
R-Th4
R-Th5
R-Th6
R-Th7
R-Th8
R-Th9
R-Th10
R-Th11
R-Th12
R-Th13
Time

**Figure S14**

**Figure S14. Mouse thoracic network model predicts synchronous or alternating responses depending on input**

**A.** Mouse thoracic spinal volume, populated by eight premotor interneuron classes and MMC motor neurons. Thirteen ventral roots are evenly spaced along the thoracic length.

**B**. Fractional distribution of thoracic cell types.

**C.** Emergent thoracic network connectivity matrix after combining thoracic cell type composition with mouse-like projection biases, as in **Figure 6.**

**D.** Transversal view of the thoracic mouse model with highlighted MMCs (dark blue, enlarged). Remaining cells in gray. Two input modalities are illustrated: Tonic global (pink) and vestibulospinal (light blue) drive. In the input/output curves, dashed and solid denote left and right hemicords, respectively. Network response was measured through the middle thoracic roots, L-Th7 (left) and R-Th7 (right).

**E.** Steady state activity traces at ventral root 6, 7 and 8 in response to global tonic drive. Right, solid. Left, dashed.

**F**. Phase correlation between steady state rhythms along (ventral root 6, 7 and 8) and across (right, solid; left; dashed) the cord in response to global tonic drive. Bar plots show data points from n=20 network samples. Network rhythms are predominantly left-right synchronous but propagate rostro-caudally along the cord.

**G**. Vestibular drive input statistics and output dynamics. Top left bar plot shows fractional distribution of cell types (colors) and vestibular input (blue). Top right bar plot shows distribution of left-right phase differences at Th7 in response to vestibular drive. N=100 network samples. Bar height shows median phase difference. Bottom panel shows Th7 activity traces (right, solid; left, dashed) of the median network sample. Amplitudes are in arbitrary, standardized units.

**H**. Steady state spatiotemporal activity of right hemicord MMC motor neurons in response to global, tonic network drive. Neurons are sorted rostrocaudally (vertical axis). Horizontal axis shows time. Position of the thirteen right thoracic roots displayed on the right-hand side.

| Table S1. HCR probe set sequences | | | |
|---|---|---|---|
| **Target Gene** | **Amplifier** | **Pool Pair** | **Sequence** |
| *pax3* | B2 | 1 | CCTCGTAAATCCTCATCAaaCAGCTCCAGCCAAGCTGGTCATCAT |
| *pax3* | B2 | 1 | CTGGGCATGGTCTCATCATTCTTGGaaATCATCCAGTAAACCGCC |
| *pax3* | B2 | 2 | CCTCGTAAATCCTCATCAaaGGAATCCAGTCCGGGGATAGTTCTG |
| *pax3* | B2 | 2 | GTCCTAAAGGAGTGGACACTTCCAGaaATCATCCAGTAAACCGCC |
| *pax3* | B2 | 3 | CCTCGTAAATCCTCATCAaaTGCCATCTCAACTATCTTGTGCCTT |
| *pax3* | B2 | 3 | GATGACGCAGGGTCGGATGCCATGGaaATCATCCAGTAAACCGCC |
| *pax3* | B2 | 4 | CCTCGTAAATCCTCATCAaaGCCGTGCGACACTCTGAGCTGGCGC |
| *pax3* | B2 | 4 | GTAGCGACACAGGATCTTGGAGACGaaATCATCCAGTAAACCGCC |
| *pax3* | B2 | 5 | CCTCGTAAATCCTCATCAaaTTTCCACTTCTGGAGTAGTCACCTG |
| *pax3* | B2 | 5 | TGTCTCTTTTAAATTCTTCGATCTTaaATCATCCAGTAAACCGCC |
| *pax3* | B2 | 6 | CCTCGTAAATCCTCATCAaaGAATTTCCCAGCTGAACATACCCGG |
| *pax3* | B2 | 6 | ATACTCCGTCTTTAAGTAATTTATCaaATCATCCAGTAAACCGCC |
| *pax3* | B2 | 7 | CCTCGTAAATCCTCATCAaaTTACTGAAGGCACAGTATTGCGATC |
| *pax3* | B2 | 7 | TGCTCCTCAGAATTCGGCTGATGGAaaATCATCCAGTAAACCGCC |
| *pax3* | B2 | 8 | CCTCGTAAATCCTCATCAaaTGTCCTCCTCATCCCCTTTCCCAAA |
| *pax3* | B2 | 8 | CTTCTTGCTCCTTTCGATCCAGCTCaaATCATCCAGTAAACCGCC |
| *pax3* | B2 | 9 | CCTCGTAAATCCTCATCAaaTGCTATGCTTTGCTCTCTTCTCGCT |
| *pax3* | B2 | 9 | GAGCTCTTTCCCTTAGGATCCCATCaaATCATCCAGTAAACCGCC |
| *pax3* | B2 | 10 | CCTCGTAAATCCTCATCAaaAGCCTTCCTCAGACTCAGGGGATGC |
| *pax3* | B2 | 10 | GCAGGTCTGGTTCAGAGTCAATATCaaATCATCCAGTAAACCGCC |
| *pax3* | B2 | 11 | CCTCGTAAATCCTCATCAaaTGCTCCTGCGCTGCTTCCTCTTCAG |
| *pax3* | B2 | 11 | CCAGTTGCTCTGCAGTGAATGTGGTaaATCATCCAGTAAACCGCC |
| *pax3* | B2 | 12 | CCTCGTAAATCCTCATCAaaTTCTCTCGAATGCTCTCTCTAACTC |
| *pax3* | B2 | 12 | CTCGTGTATAAATATCCGGGTAGTGaaATCATCCAGTAAACCGCC |
| *pax3* | B2 | 13 | CCTCGTAAATCCTCATCAaaTGAGCTTGGCTCTCTGGGCCAGTTC |
| *pax3* | B2 | 13 | TAAACCACACCTGAACTCGCGCCTCaaATCATCCAGTAAACCGCC |

| Table S1. HCR probe set sequences | | | |
|---|---|---|---|
| *pax3* | B2 | 14 | CCTCGTAAATCCTCATCAaaGCTTTCTCCATCTAGCGCGTCGGTT |
| *pax3* | B2 | 14 | ATGCCATGAGCTGGTTGGCTCCTGCaaATCATCCAGTAAACCGCC |
| *pax3* | B2 | 15 | CCTCGTAAATCCTCATCAaaGAAAGCCCCTGGGATCAAGTGGTT |
| *pax3* | B2 | 15 | TTGGCAGAGCTGGCATAGCTGTAGGaaATCATCCAGTAAACCGCC |
| *pax3* | B2 | 16 | CCTCGTAAATCCTCATCAaaGGTAAGAGGTCTCAGATAACTGGTA |
| *pax3* | B2 | 16 | ACACAGCTTGTGGTATAGAAGTGGGaaATCATCCAGTAAACCGCC |
| *pax3* | B2 | 17 | CCTCGTAAATCCTCATCAaaGCCTGTGAACTGTGTTGCTTGGATC |
| *pax3* | B2 | 17 | GCACACTGCTTGGAGGGAGCGGCTGaaATCATCCAGTAAACCGCC |
| *pax3* | B2 | 18 | CCTCGTAAATCCTCATCAaaTGTCGGGGTTGGAAGGAAGACTTTG |
| *pax3* | B2 | 18 | TGCTGGGCAGGCAATAGGCCGAACTaaATCATCCAGTAAACCGCC |
| *pax3* | B2 | 19 | CCTCGTAAATCCTCATCAaaACAAAGCTGTCTGTGTAGCTGGAAA |
| *pax3* | B2 | 19 | ATAGGGTTGGAAGGCCCAGATGGGGaaATCATCCAGTAAACCGCC |
| *pax3* | B2 | 20 | CCTCGTAAATCCTCATCAaaGAAAGGCCGTTGCCAATGGCTGGGT |
| *pax3* | B2 | 20 | TTAGTCAGGAGACCCATAACCTGAGaaATCATCCAGTAAACCGCC |
| *pax3* | B2 | 21 | CCTCGTAAATCCTCATCAaaTGAGGCTGATGAGGAACCCCACCAT |
| *pax3* | B2 | 21 | GTTAAAGGAGATAAGGCATAATCCGaaATCATCCAGTAAACCGCC |
| *pax3* | B2 | 22 | CCTCGTAAATCCTCATCAaaACAGCTGTGGGAGGCTCAAGGCCTC |
| *pax3* | B2 | 22 | TCCAGTCTCTGGCTGCAACTTGCTGaaATCATCCAGTAAACCGCC |
| *pax3* | B2 | 23 | CCTCGTAAATCCTCATCAaaGATAGACTGTCCAAGCTCTTCATAT |
| *pax3* | B2 | 23 | GTTGGTGGGCAATAAGACTGTGATGaaATCATCCAGTAAACCGCC |
| *pax3* | B2 | 24 | CCTCGTAAATCCTCATCAaaTCCATGCTGTAACCTGAGGTGCTGT |
| *pax3* | B2 | 24 | TGTGGGTATTGGTATCCTGTCATAGaaATCATCCAGTAAACCGCC |
| *olig2* | B5 | 1 | CTCACTCCCAATCTCTATaaAAGGAGCAGCATCGCATTGTGACAT |
| *olig2* | B5 | 1 | TCTTGGCAGGGTGTGCCAGCGACCAaaCTACCCTACAAATCCAAT |
| *olig2* | B5 | 2 | CTCACTCCCAATCTCTATaaTAGGCAGCGCTGAGAACAATGTCAT |
| *olig2* | B5 | 2 | CAATTTTCCCTCTGCCTCCGCTTTTaaCTACCCTACAAATCCAAT |
| *olig2* | B5 | 3 | CTCACTCCCAATCTCTATaaCTCCAATGCCTTGTGTGACAGTGAT |

| Table S1. HCR probe set sequences | | | |
|---|---|---|---|
| *olig2* | B5 | 3 | GCAGGTGCAGTGGAGGCGCTTGTGTaaCTACCCTACAAATCCAAT |
| *olig2* | B5 | 4 | CTCACTCCCAATCTCTATaaAGGGCCTGTTGGCAGTGCTGAGATA |
| *olig2* | B5 | 4 | CGTCAGAATCCATGGGGCAGAGAGAaaCTACCCTACAAATCCAAT |
| *olig2* | B5 | 5 | CTCACTCCCAATCTCTATaaAGGAGGCTCTGCTGGAGCCCAGGCT |
| *olig2* | B5 | 5 | GTAGGAACAGATCGTCTGTCTCCGGaaCTACCCTACAAATCCAAT |
| *olig2* | B5 | 6 | CTCACTCCCAATCTCTATaaCGCCCGAGAAACCACCTTTCCGCGA |
| *olig2* | B5 | 6 | AGTCGCTCATGGTGGAAGAAGACACaaCTACCCTACAAATCCAAT |
| *olig2* | B5 | 7 | CTCACTCCCAATCTCTATaaGTTGCGCAACTCGGCGCTCAGCTCC |
| *olig2* | B5 | 7 | TCCCGCCGCCGCGGCCAAAGCCATAaaCTACCCTACAAATCCAAT |
| *olig2* | B5 | 8 | CTCACTCCCAATCTCTATaaGCCGCCGCTGCTTTAAAAGCGCCCA |
| *olig2* | B5 | 8 | TTCTTGGCCGCCACTGCCGCCGCAGaaCTACCCTACAAATCCAAT |
| *olig2* | B5 | 9 | CTCACTCCCAATCTCTATaaTCCGACTCGTTCATCTGCTTCTTGT |
| *olig2* | B5 | 9 | TTTATCTTCAGCCTCAGGTGCTGCAaaCTACCCTACAAATCCAAT |
| *olig2* | B5 | 10 | CTCACTCCCAATCTCTATaaTCGTGCATCCGCTTGCGCTCCCGGC |
| *olig2* | B5 | 10 | CGCAGCCCGTCCATTGCGATATTCAaaCTACCCTACAAATCCAAT |
| *olig2* | B5 | 11 | CTCACTCCCAATCTCTATaaGGTCCATGCGCGTAAGGCATCACCT |
| *olig2* | B5 | 11 | GCGATCTTGGACAGTTTGCGCACTGaaCTACCCTACAAATCCAAT |
| *olig2* | B5 | 12 | CTCACTCCCAATCTCTATaaATGTAATTGCGCGCCAGCAGCAGAG |
| *olig2* | B5 | 12 | TCCTCCAGCGAGTTGGTGAGCATGAaaCTACCCTACAAATCCAAT |
| *olig2* | B5 | 13 | CTCACTCCCAATCTCTATaaCCCCGTAGATCTCGCTGACCAGGCG |
| *olig2* | B5 | 13 | CTGCGGCTGCCGCATGGTGGTGGTGaaCTACCCTACAAATCCAAT |
| *olig2* | B5 | 14 | CTCACTCCCAATCTCTATaaACCATGGGAGGCGGGTGGAGCAGGA |
| *olig2* | B5 | 14 | AGCCCCTGGACTGGGAGACTTGAGGaaCTACCCTACAAATCCAAT |
| *olig2* | B5 | 15 | CTCACTCCCAATCTCTATaaAGGACAGGGCATTCCGCTCCAATGG |
| *olig2* | B5 | 15 | AGCAGATGGCACTTGGCACATGCTAaaCTACCCTACAAATCCAAT |
| *olig2* | B5 | 16 | CTCACTCCCAATCTCTATaaGGTGGCAGTTGTGAGATGGTGATGG |
| *olig2* | B5 | 16 | CTTGGCATCTGTGCTGAGCCTGGACaaCTACCCTACAAATCCAAT |
